# Clonal dynamics and somatic evolution of haematopoiesis in mouse

**DOI:** 10.1101/2024.09.17.613129

**Authors:** Chiraag D. Kapadia, Nicholas Williams, Kevin J. Dawson, Caroline Watson, Matthew J. Yousefzadeh, Duy Le, Kudzai Nyamondo, Alex Cagan, Sarah Waldvogel, Josephine De La Fuente, Daniel Leongamornlert, Emily Mitchell, Marcus A. Florez, Rogelio Aguilar, Alejandra Martell, Anna Guzman, David Harrison, Laura J. Niedernhofer, Katherine Y. King, Peter J. Campbell, Jamie Blundell, Margaret A. Goodell, Jyoti Nangalia

## Abstract

Haematopoietic stem cells maintain blood production throughout life. While extensively characterised using the laboratory mouse, little is known about how the population is sustained and evolves with age. We isolated stem cells and progenitors from young and old mice, identifying 221,890 somatic mutations genome-wide in 1845 single cell-derived colonies, and used phylogenetic analysis to infer the ontogeny and population dynamics of the stem cell pool. Mouse stem cells and progenitors accrue ∼45 somatic mutations per year, a rate only about 2-fold greater than human progenitors despite the vastly different organismal sizes and lifespans. Phylogenetic patterns reveal that stem and multipotent progenitor cell pools are both established during embryogenesis, after which they independently self-renew in parallel over life. The stem cell pool grows steadily over the mouse lifespan to approximately 70,000 cells, self-renewing about every six weeks. Aged mice did not display the profound loss of stem cell clonal diversity characteristic of human haematopoietic ageing. However, targeted sequencing revealed small, expanded clones in the context of murine ageing, which were larger and more numerous following haematological perturbations and exhibited a selection landscape similar to humans. Our data illustrate both conserved features of population dynamics of blood and distinct patterns of age-associated somatic evolution in the short-lived mouse.

## Introduction

The haematopoietic system sustains mammalian life through the continuous generation of oxygenating red blood cells, an array of immune cells, and platelets that course through all tissues. In a large animal such as the human, blood production accounts for 86% of daily cellular turnover, generating ∼280 billion cells per day^1^. This process relies on a hierarchy of progenitors that successively amplify cellular output towards fully differentiated blood cells. All are believed to ultimately derive from rare haematopoietic stem cells (HSCs), a heterogeneous pool^2–4^ maintained in a relatively protected state to support blood production throughout life.

HSCs are the best-studied and utilised of somatic stem cells. They are the basis for life-saving bone marrow transplantation and have been purified from humans and mice for studies on their molecular regulation. HSCs are capable of being activated by various stimuli, such as infection and bleeding^5–8^, in order to rapidly replenish differentiated blood cells as needed, and concomitantly undergo controlled self-renewal to sustain the stem cell pool over time.

Like all somatic cells, HSCs accumulate somatic mutations with age^9–13^. In humans, some mutations promote cellular fitness, driving clonal outgrowth during normal ageing^9^. Such ‘clonal haematopoiesis’ (CH), while remaining at very low levels in younger individuals, is ubiquitous in the elderly where it results in a dramatic loss of clonal diversity^9^. CH is a known risk factor for blood cancers and age-associated non-cancerous disease, and may encode other ageing phenotypes^9,14–17^. Extensive clonal expansions have also been described across human tissues where they are associated with ageing, cancer and other diseases, reflecting the consequences of lifelong somatic evolution^18–22^. Whether these patterns of somatic evolution are also features of ageing in other species is unknown.

Within *Mammalia*, the rate of somatic mutation accrual in colonic epithelium inversely scales with lifespan; that is, species acquire a similar magnitude of mutations by the end-of-life independent of lifespan^23^. However, it is unclear if this pattern extends to other tissues beyond the colon and whether the consequences of somatic evolution over human life also scale to shorter-lived species.

The inbred laboratory mouse is used ubiquitously across biomedical research. It has been used extensively to study haematopoiesis, leading to fundamental tenets of somatic stem cell biology. The most commonly used strain, C57Bl/6J, has a median lifespan of 28 months^24^, 1/35th that of humans, and broadly recapitulates many phenotypic features of human ageing, with some preliminary data suggesting a lower rate of CH^25^. Here, we study the ontogeny, clonal dynamics, and selection landscapes of murine HSC populations *in vivo* to understand the evolutionary processes shaping the maintenance and ageing of blood production.

## RESULTS

### Whole genome sequencing of hematopoietic stem and progenitor cells

To study somatic mutagenesis and clonal dynamics in the haematopoietic compartment within the laboratory mouse, we purified HSCs from three young (3-months) and three aged (30-months) healthy C57BL/6J female mice (Fig.1A, Extended Data Fig.1A), ages estimated to be human lifespan equivalents of ∼20 and 85-90 years respectively (Supplementary Note 1). A longstanding consensus is that haematopoiesis is supported by long-term stem cells (LT-HSCs, henceforth, referred to as HSCs) which give rise to multipotent progenitors (MPPs, sometimes called short-term HSCs). Extensive functional analysis established that both HSCs and MPPs, distinguished on the basis of their cell-surface markers, can support haematopoiesis and produce all differentiated blood cell types, but HSCs can engraft hosts following multiple rounds of serial transplantation, whereas MPPs cannot ^26–31^. Therefore, we examined both these populations *in vivo*. Single HSCs and MPPs were expanded *in vitro* to produce colonies (Fig.1A, Extended Data Fig.1B) for whole-genome DNA sequencing at an average depth of 14X. From individual 3-month and 30-month animals (n=6), we sequenced 61-235 HSC-derived colonies and 70-191 MPP-derived colonies (Fig.1B). We also purified fewer HSCs from 17 additional mice aged 1 day to 30 months (total 242, ranging from 9-24 cells per animal). Following exclusion of 139 colonies due to low sequencing coverage or lack of clonality (Extended Data Fig.1C and Methods), 1547 whole genomes (908 HSCs, 639 MPPs) were taken forward for somatic mutation identification and phylogenetic reconstruction.

**Figure 1.**
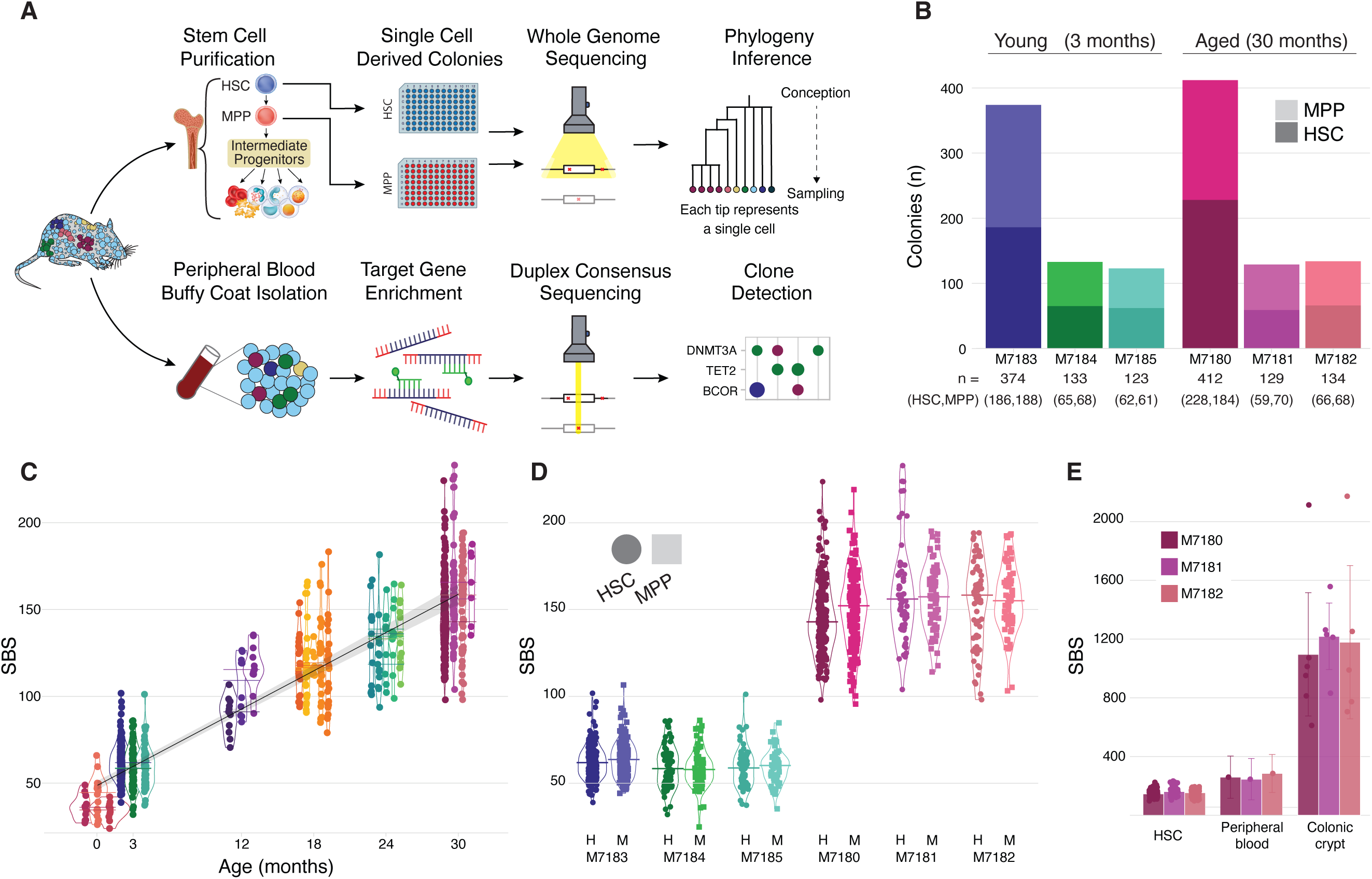
Somatic mutations in murine stem cell-derived haematopoietic colonies. **A)** Study approach. Single-cell derived colony whole genome sequencing (WGS) of long-term haematopoietic stem cells (HSC) and multipotent progenitors (MPP) to study somatic mutations, lineage relationships and population dynamics, top; targeted duplex-sequencing of peripheral blood to identify small clonal expansions and fitness landscapes, bottom. **B)** Number of whole genomes (n=1305) of HSC- and MPP-derived colonies that underwent phylogenetic construction for each female mouse (n=6). Plots are coloured according to HSC- or MPP-derived colonies, darker and lighter shades, respectively. **C)** Burden of individual single base substitutions (SBS) observed in HSCs (n=908) from each donor. Points are coloured as in panel B. Line shows linear mixed-effect regression of mutation burden observed in colonies. Shaded areas indicate the 95% confidence interval. **D)** Comparison of SBS burden between HSC- and MPP-derived colonies from the same mice. SBS burden from HSCs are shown as circles and burden from MPPs are shown as squares. H, HSC; M, MPP, shown above animal ID. **E)** SBS burden across HSCs (data as in panel C), whole blood, and individual colonic crypts in the three aged mice. Error bars denote 95% confidence interval. Peripheral blood and colonic crypt somatic mutation burdens were measured with nanorate sequencing and WGS, respectively.

### Somatic mutation accumulation in murine haematopoietic stem cells

Comparison of HSCs from young and old mice revealed a constant rate of somatic mutation accumulation with age (Fig.1C). Mice aged 3 months had an estimated 59.5 single base substitutions (SBS) (95% confidence interval, CI, 57.3-61.7), and by 30 months had acquired 161.4 SBS per HSC (CI 155.1-167.8), corresponding to 45.3 SBS per year (CI 42.2-48.4) or a somatic mutation being acquired every 8-9 days within HSCs. Across the diploid mouse genome, this reflects a mutation rate of 8.3⋅10^-9^ bp per year (CI 7.7-8.9⋅10^-9^ bp/year). Few insertions-deletions were captured per colony (median 1, range 0-4) with no chromosomal changes observed. Previous studies suggest that MPPs are a more rapidly cycling population^5,32,33^ thought to amplify cell production from HSCs, which could result in a greater mutation burden. However, there was no difference in mutation burden between HSCs and MPPs (Fig.1D), consistent with observations from human blood wherein no appreciable differences in somatic mutation burdens between HSCs and more differentiated blood cells are apparent^11,12^.

The murine HSC SBS rate is about twice that of humans (14-17 SBS per year)^9–11,34^, given their similar genome sizes. This is consistent with the concept that somatic mutation rates are negatively correlated with lifespan^23^ such that short-lived species have higher rates of somatic mutation accumulation than longer-lived animals. However, the ∼10-fold difference in ultimate mutation burden (∼150 in HSCs from 30-month-old mice vs >1,500 in human HSCs of an equivalent age of 85-90 years) is much greater than expected given that total end-of-life somatic mutation numbers in mammalian intestinal crypts show low variation regardless of life-span^23^. Thus, we wished to validate the lower-than-expected somatic mutation burden observed in aged murine stem cells.

First, we compared genome-wide mutation burdens in HSCs with that of matched intestinal crypts from the same three aged mice. Following WGS of individually microdissected clonal crypts (n=16, range 5-6 per sample, Extended Data Fig.1D, Methods), we confirmed that colonic epithelium exhibited substantially higher mutation burdens, similar to that reported previously^23^ (Fig.1E), confirming that we were not underestimating mutations in HSCs. Secondly, we undertook independent nano-error rate whole-genome duplex sequencing^12^ of matched whole blood from the three aged animals. This method identifies mutations in single DNA molecules and, thus, can orthogonally estimate mutation burden from peripheral blood. The mutation burden was not statistically different from that of haematopoietic colonies (Fig.1E). We did note a non-significant trend towards higher mutation burden estimates from whole blood than HSC colonies – this is likely due to whole blood including lymphoid cells which have higher mutation burdens^35^. Despite whole blood having a mixture of mature cell types and the different sequencing technologies used, these data confirm that somatic mutation rates in blood do not inversely scale with lifespan to the same degree as observed in colon.

### Aetiology of mutational processes in haematopoietic stem cells in mouse

The pattern of sharing of somatic mutations across individual colonies can be used to reconstruct a phylogenetic tree that depicts their ancestral lineage relationships (Methods). We use the term ‘lineage’ here to represent the direct line of descent rather than different blood cell types. Figure 2 shows the phylogenetic trees for a 3-month- and 30-month-old mouse, with additional young and aged phylogenies in Extended Data Figure 2. At the tips of the trees are individual HSC-(blue) and MPP-derived colonies (red); the branches that trace upwards from each colony to the root of the tree reflect the somatic mutations present in that individual colony and how these mutations are shared across other colonies. Individual branchpoints (“coalescences”) represent ancestral cell divisions wherein descendants of both daughter cells were captured at sampling. Colonies that share a common ancestor on the phylogeny are referred to as a clade.

**Figure 2.**
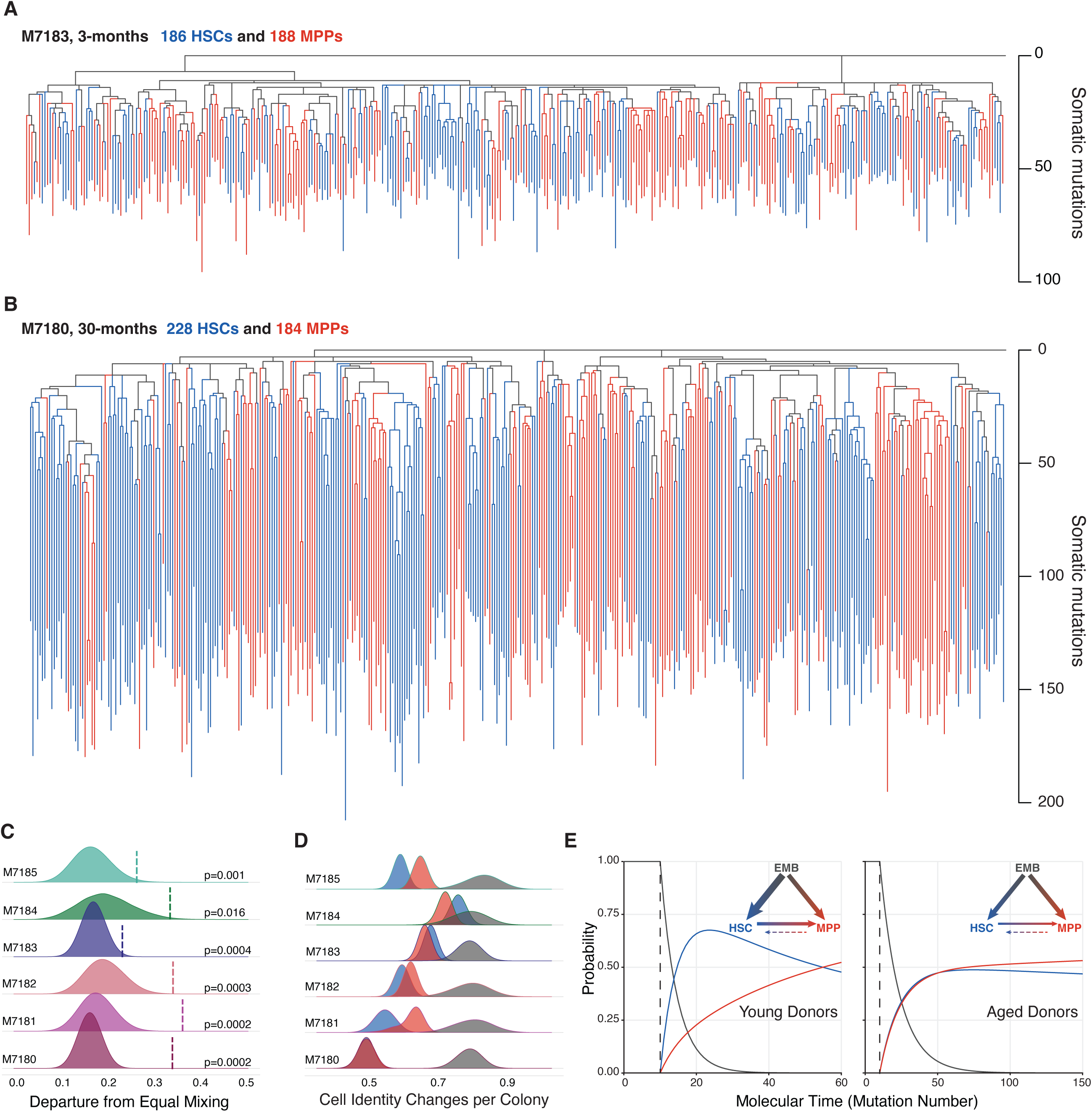
Phylogenetic trees of HSCs and MPPs from a young and old mouse. **A-B)** Phylogenies were constructed from young (3-months, A) and aged (30-months, B) female mice using the pattern of sharing of somatic mutations among HSC (blue) and MPP (red) colonies. Each tip represents a single colony. Branch lengths represent mutation number, corrected for varying sequencing depth of descendant colonies. Branches and coalescence colours reflect the identity of descendent colonies with HSCs in blue and MPPs in red, respectively. Branches where we are unable to infer the established cell type for one or more lines of descent are coloured black. **C)** To determine the degree of phylogenetic relatedness between HSC and MPP, we measured the amount of HSC-MPP mixing within clades. If an MPP had a recent HSC ancestor, clades should contain both cell types. We thus compared the “observed” versus “expected-by-chance” clade mixing behaviour. The mixing metric for a clade is the absolute difference between the proportion of HSCs in a clade and the expected value under equal sampling, 0.5; this metric is then averaged for all clades in a phylogeny. The vertical bar reflects the observed average clade mixing metric within the constructed phylogenies. The filled distributions reflect average clade mixing metrics that would be expected by random chance or more frequent intermixing of HSCs and MPPs, and were generated by reshuffling the tip cell identities within the tree. HSC or MPP colonies are designated as being in the same clade if they share a most recent common ancestor after 25 mutations, corresponding to early foetal development. Only clades with more than 3 colonies are considered. **D)** Distributions of the number of cell identity changes required per colony to capture the observed tip states. The number of cell identity changes assuming an ‘HSC-first’ model (HSCs first give rise to MPPs) is shown in blue. The required cell identity changes for the opposite ‘MPP-first’ model, in which MPPs first give rise to HSCs, is shown in red. The null distribution, in which tip states are randomly reshuffled is shown in grey. **E)** Cell-type probability trajectories displaying specification to HSC or MPP states under a simple 3-state ontogeny model (Methods). In 30-month donors (right), we observe equal generation of HSC and MPP from embryonic progenitors (EMB) and can reject an “HSC-first” model. In 3-month donors (left), we observe relatively increased generation of HSCs from EMB and can not reject an “HSC”-first model. The displayed trajectories are based on iterating the maximum likelihood based Markov chain starting at the embryonic state. Thickness of arrows reflect the proportion of overall transitions from the EMB state to HSC and MPP states, and between HSC and MPP states. The cell identity transition rates are derived in Supplementary Note 2.

First, we wished to understand the aetiology of the higher rate of mutation accumulation in murine HSC and MPPs compared to human HSCs. DNA replication during cell division is one source of mutations reflecting DNA polymerase base incorporation errors. To estimate the rate of DNA replication-associated somatic mutation accumulation in mice, we studied the distribution of nodes on the phylogenies with more than two descendant lineages, termed polytomies.

These are evidence of ancestral cell divisions which were not associated with the acquisition of a somatic mutation and can be used to infer the average number of mutations that are acquired per cell division^10^ (Extended Data Fig.3). We focused on the roots of the trees where we capture the greatest number of coalescences, both due to the small population size and the rapid divisions at this point in life. We observed 266 lineages by 12 mutations of molecular time in five donors that had adequate (>10 lineages) diversity. Of the 265 symmetrical self-renewing cell divisions that would have required, 44 were mutationally silent, leading to a mutation rate estimate of 1.80 (95% CI: 1.46-2.19) mutations per cell division during early life (Extended Data Fig.3). This estimate is not significantly different from that previously observed in humans (1.84 mutations/cell division, p=0.5)^13^, suggesting that excess mutation accumulation is not occurring due to poorer fidelity during DNA replication in murine stem cells.

Mutagenic biological processes yield distinguishable patterns of base substitutions at trinucleotide sequence contexts, termed mutation signatures. We identified three mutational processes (Extended Data Fig.4A, Methods): i) SBS1 reflecting the spontaneous deamination of methylated cytosines, ii) SBS5 likely produced by cell-intrinsic damage and repair processes, and iii) SBS18 characterised by C>A transversions potentially linked to oxidative damage. Substitutions attributed to SBS1 and SBS5 increased with age (8.64 SBS1/month and 32.52 SBS5/month), keeping with their clock-like nature across species; indeed, these processes account for most mutations in healthy human HSCs. Mutations attributed to SBS18 (mean 5.3, range 1.5-18 per colony, Extended Data Fig.4B), were previously identified in murine colorectal crypts^23^, but did not appear to accumulate with age. To explore the timing of SBS18 mutations, we deconvoluted branch-specific mutations (Extended Data Fig.4C). SBS18 accrued before three months of life, followed by a plateau (Extended Data Fig.4D), suggesting a specific early-life vulnerability to these mutations, reminiscent of their presence in human placenta and human foetal HSCs^13,36^. Taken together, the higher relative somatic mutation accumulation rate in mice is underlaid by context-specific mutational processes (SBS18) and a higher rate of endogenous DNA damage and reduced repair (SBS1 and 5).

### Origin and parallel establishment of HSC and MPP populations

We next sought to examine the lineage relationships between the HSCs and MPPs. Classical models of the haematopoietic differentiation hierarchy propose that MPPs derive from HSCs^37,38^. In recent years, a more nuanced and dynamic picture has emerged, with the identification of additional self-renewing progenitor compartments^2,4^. Using our phylogenetic approach, stem cell ontogeny can be retraced *in vivo* during unperturbed haematopoiesis. Working up from the phylogenetic “tip” states of HSC (blue) or MPP (red), we infer the identity of ancestral branches and coalescences based on the identity of their nearest sibling cell (detailed in Supplemental Note 2). Branches where we are unable to infer the established cell type for one or more lines of descent are coloured black. We observed clear vertical bands of HSC-only (all “blue”) and MPP-only (all “red”) ancestral lineages across the trees representing independent clades (Fig.2A,B, Extended Data Fig.2), with a minority of HSCs (“blue” tips) being sampled from MPP (“red” clades) and vice versa. The clear separation of MPPs and HSCs suggests that most HSCs are derived from HSC self-renewing divisions, and most MPPs are derived from MPP self-renewing divisions, with each population independently self-renewing in parallel throughout life. If HSCs and MPPs were closely related to one another, as might be the case if MPPs were recently generated from HSCs, then one would expect the two cell types to be intermixed across the phylogenetic tree, with individual clades (cells derived from a common ancestor) containing cells of both types. However, we observed that clades were largely uniform in composition, containing more cells of the same type than would be expected if the population of HSCs and MPPs were intermixed (Fig.2C). This phylogenetic separation of HSC and MPPs provides strong evidence that these two populations independently contribute to blood production in the mouse.

The lack of intermixing between HSCs and MPPs on the phylogenetic trees suggests long-term inheritance of the HSC or MPP ‘state’, presumably encoded epigenetically. Therefore, we next explored when such sustained MPP and HSC cell state commitments may have occurred during life. Coalescences near the top of a phylogeny (near the root) reflect cell divisions that occur soon after conception. Most branches and coalescences here are ‘black’ (Fig.2A,B, Extended Data Fig.2), as no established HSC or MPP lineages could be inferred at this time. Stable heritable identity of either HSC or MPP appears established at similar times – by around 25 mutations of molecular time suggesting that a substantial proportion of the HSC and MPP populations appear to diverge early in life. The mixed effects regression model of mutation rate suggests that ∼50 somatic mutations may be acquired before birth (intercept of mixed effects model, 48.2, CI 45.61-50.8, Methods). Thus, HSCs and MPPs are likely established in parallel during foetal development.

To explore when *in utero* this was occurring, we evaluated somatic mutations present in both HSCs and colonic crypts from the same animals – by definition, mutations shared between these tissues arose in a common ancestor whose progeny contributed to both blood and colonic epithelium. As blood is derived from mesoderm and colonic epithelium is derived from endoderm, any shared mutations must have occurred in embryonic cells prior to gastrulation. Mutations on the haematopoietic phylogeny were observed in sampled colonic crypts (n=4-6 crypts per 30-month-old mouse) down to 9-11 mutations of molecular time (Extended Data Fig.3), with decreasing representation of mutations further from the phylogeny root, timing these shared mutations to have occurred during gastrulation. Indeed, branches with an inferred HSC or MPP identity did not share mutations with the colon, consistent with these lineages being established after germ layer specification.

Given the likely embryonic establishment of distinct HSC and MPP pools, we next considered the simplest series of cell state changes (eg HSC to MPP, or HSC to MPP, etc) that might be required to capture the observed cell identities. We first considered the prevailing view, that MPPs are generated from HSCs, such that HSC fate occurs prior to specification of MPP. We counted the number of cell identity changes required to reach the sampled cell identity. Surprisingly, the HSC-to-MPP model was equivalently parsimonious (requiring a number of cell state changes that was not statistically different) to a model where all cells start as MPP (with HSCs able to arise from MPPs), an ontogeny not generally considered likely (Fig.2D, also see Supplementary Note 2). Overall, our data suggest that many long-term HSC and MPP lineages are established independently and in parallel during early development, and that MPPs do not always arise from HSCs, contrary to classical haematopoiesis models.

### Modelling HSC and MPP establishment and transitions

To formalise the above ideas and develop an ontogeny model for HSCs and MPPs that fits with our observed data, we developed a hidden Markov tree model. The Markov approach allows estimation of the rates at which a cell state makes transitions as it evolves through time. We defined three unobservable ancestral states: embryonic precursor cell (EMB), HSC, and MPP. We then used the observed outcomes of HSC versus MPP tip states to infer both the sequence and the transition rates between these states during life (Methods, Supplementary Note 2). We considered all cells prior to gastrulation (<10 mutations) as EMB, and then assumed that in each unit of molecular time, there is a fixed probability of transitioning out of this state to either an HSC or MPP state (with subsequent fixed probabilities of further transitions such as HSC-to-MPP). In addition to characterising the most feasible model parameters that fit the observed data using a maximum likelihood approach (Methods), we also estimated the rate of HSC to MPP (and vice-versa) transitions during life, to account for any HSC/MPP mixed clades (Supplementary Note 2).

We fitted the above model to i) each donor, ii) each age group, and iii) the whole cohort. Based on a nested likelihood ratio test analysis, we found that the model fitted to each age group (young and old mice separately) was most consistent with our data (Supplementary Note 2). Across the whole cohort, we found that a model in which EMB can transition to either HSC or MPP was a significantly better fit than an “HSC-first” model, where all EMB must transition to HSC prior to any MPP specification. However, when testing young and old mice separately, we were only able to reject an “HSC-first” ontogeny model in older mice. We could not reject an “HSC-first” model in younger mice as our data suggested more frequent HSC to MPP transitions earlier in life (Fig.2E). This apparent inconsistency in the results between young and old mice could perhaps be explained if the HSCs that produce the MPPs early in life are extinguished by old age, and thus could not be sampled for inclusion in the phylogeny. Alternatively, the rate of HSCs that transition to MPPs may be greater earlier in life. Further work is required to explore this. Interestingly, our model indicates that 50% of all HSC and MPP lineages in young and aged mice had committed to their cell state before 50 mutations of molecular time, likely before birth. As might be expected, HSC to MPP transitions were more frequent than MPP to HSC transitions, which were extremely rare (1 in 1000 transitions) and within the plausible limitations of cell-sorting accuracies. (Supplemental Note 2).

### Haematopoietic population dynamics over life

The pattern of coalescences (branch points) in a phylogenetic tree reflects the ratio (N/λ) of the overall population size (N) and the HSC self-renewing cell division rate (λ) over time – both smaller populations and more frequent cell divisions decrease the interval between coalescences. Mice haematopoietic phylogenies show a different pattern of coalescences (Fig.2A-B, Extended Data Fig.2) to equivalently-aged humans^9^. Human haematopoietic phylogenies have a profusion of early coalescences, reflecting the period of rapid cell division during embryonic growth. Coalescences are then infrequently observed due to the presence of both a large and stable HSC population by early adulthood, reappearing only in elderly human phylogenies within clonal expansions when clonal diversity dramatically collapses.

By comparison, murine haematopoietic phylogenies display coalescences continuing down the tree (see Extended Data Fig.5 for side-by-side human-mouse comparisons). These time intervals between coalescences can be used to infer the HSC population trajectory (N/λ) using a phylodynamic Bayesian framework (Methods). We observed an early period of exponential HSC growth followed by progressively increasing N/λ over the murine lifespan (Fig.3A), consistent with the observed increase in total HSC number with age by flow cytometry (Fig.3B), and other studies^39,40^. Our findings contrast with hematopoietic progenitor population trajectories in humans^9,10^ which exhibit a population growth plateau during adulthood followed by stable population size for the remainder of life. Interestingly, we infer entirely overlapping N/λ trajectories for HSCs and MPPs. Together with their similar mutation burdens and lineage independence, these data suggest that murine HSC and MPP clonal dynamics during steady state *in vivo* haematopoiesis are indistinguishable.

**Figure 3.**
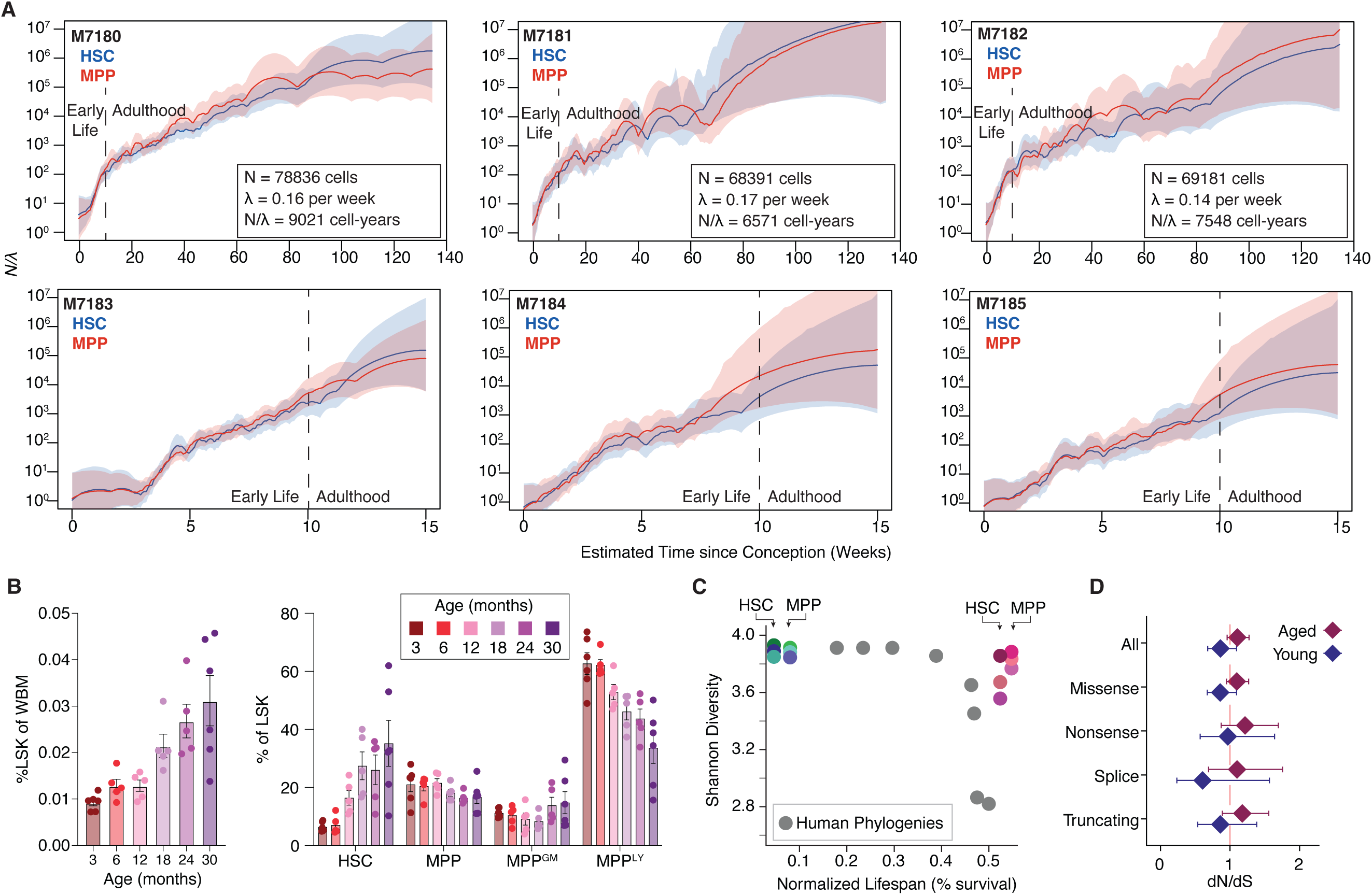
Population dynamics and selection in the murine stem cells. **A)** Population trajectories estimated separately in HSCs and MPPs using Bayesian phylodynamics for the six samples shown in Fig 2.A-B and Extended Data Fig.2. The dark blue (HSC) and red lines (MPP) indicate the mean effective population trajectory; shaded areas are 95% confidence intervals. Vertical dashed lines separate trajectories into early life and adulthood age periods, in which different population size behaviour are observed. Inset values indicate posterior density estimates of population size (N), symmetric cell division rate per week (λ), and their ratio in (N/λ) in HSC-years, as derived from approximate Bayesian computations. **B)** Haematopoietic stem and progenitor cell (HSPC) prevalence during murine ageing. The relative abundance of total HSPCs (left, defined as the LSK compartment) and individual HSPC subpopulations (right) are compared. MPP^Ly^ are lymphoid-biased progenitors, MPP^GM^ are myeloid-biased progenitors, based on current immunophenotypic definitions^55^. **C)** Shannon diversity index for each phylogeny calculated using the number and size of unique clades present at 50 mutations molecular time. Mouse points are coloured as in Fig.1B. Grey dots depict results from data published in Mitchell et al^9^. **D)** Normalised ratio of non-synonymous to synonymous somatic mutations (dN/dS) for somatic mutations observed across aged and young animals overlaps with 1 suggesting no departure from neutrality.

We next developed a joint HSC/MPP population dynamics model (given our data suggests both populations contribute equivalently to haematopoiesis), in which the population of stem cells grows towards the target population size, taking into account loss of HSC and MPP cells via cell death or differentiation (Methods). We then applied approximate Bayesian computation^41^, which generates simulations of phylogenetic trees to estimate the most likely posterior distributions of population size and symmetrical self-renewing division rates. Using this approach, we estimate that the murine HSC-MPP population grows to around 70,000 cells (median 72,414, CI 25,510-98,540). Symmetric cell divisions occur approximately every 6 weeks (median 6.4 weeks, CI 1.8-13.2 weeks). Stem cells exit the population, by either death or differentiation, about once every 18 weeks (CI 2.3-357 weeks). Posterior density estimates for each mouse are shown in Fig.3 and Extended Data Fig.6.

### Stem cell contribution to progenitors and mature blood cells

Given the observed lineage independence of HSC and MPP, and their overlapping growth trajectories, we wondered what difference *in vivo* might exist between the two populations. Therefore, we studied if HSCs and MPPs might differentially contribute to downstream lineage-biased progenitors and mature blood cells.

We isolated single cells from a mixed progenitor compartment (LSK cells) that includes granulocyte/macrophage-biased MPPs (MPP^GM^) and lymphoid-biased MPPs (MPP^Ly^) from the three aged animals. We performed whole genome sequencing on colonies from 298 LSK cells and performed phylogenetic analysis. Within the extended phylogenetic trees, we observed no discernable bias in MPP^GM^ or MPP^Ly^ emerging preferentially from HSC versus MPP (Extended Data Fig.7A-C). To evaluate this more formally, we separately examined clades that contained MPP^GM^ or MPP^Ly^ and evaluated if they were more phylogenetically linked to HSCs or MPPs than expected by chance. Neither MPP^GM^ nor MPP^Ly^ preferentially derived from either HSC or MPP beyond random chance (Extended Data Fig.7D-E), confirming that both HSCs and MPPs produce these downstream progenitors at seemingly similar proportions. However, these data are limited by a relatively low number of sampled MPP^GM^ and MPP^Ly^.

We next evaluated if peripheral blood cells were preferentially derived from HSCs or MPPs. We performed deep targeted sequencing on peripheral blood DNA from the three aged mice for a subset of mutations displayed on the corresponding phylogenetic trees (Methods). The fraction of cells in peripheral blood harbouring a mutation present on the phylogenetic tree can be used to estimate how much that lineage contributes to blood production. For example, if a single cell or lineage contributed avidly to differentiated progeny, then its mutations would be seen at high proportion (variant allele frequency, VAF) in peripheral blood. We recaptured mutations in the peripheral blood that were acquired in both ancestral HSCs and ancestral MPPs, suggesting that both these cell types actively contribute to mature blood production. Mutations private to single cells on the phylogeny were subclonal, occurring below 0.1% VAF in peripheral blood (Extended Data Fig.8A) in line with each HSC/MPP contributing only a small amount of overall blood production. While both HSC and MPP ancestral lineages gave rise to peripheral blood, we observed a slight bias toward increased representation of ancestral MPP lineages compared to HSCs, though this difference was subtle (Extended Data Fig.8B). This subtle difference may be due to increased proliferation of MPP descendants, or differences in compartment population size earlier in life; we cannot distinguish between these possibilities.

### Absence of large clonal expansions in aged mice

A striking feature of the phylogenetic trees in aged mice is the uniform distribution of long branches with no expanded clades (Fig.3C, Extended Data Fig.5). This indicates mouse haematopoiesis maintains clonal diversity instead of collapsing into an oligoclonal state as observed in elderly humans^9^. Indeed, our population dynamics simulations confidently recapitulated observed phylogenies under a model of neutral growth in the absence of selection. Concordantly, no colonies (n=1305) displayed mutations in murine orthologues of genes associated with human clonal haematopoiesis (CH), which could act as potential driver events, aligning with a topology devoid of observable late-life clonal exponential growth. Among 49,849 SNVs observed across young and aged samples, the relative rate of nonsynonymous mutation acquisition also did not significantly depart from neutrality (Fig.3D), with no novel genes identified as being under selection. This indicates that positive selection does not explain the catalogue of somatic mutations observed.

Given that mutation entry, which furnishes a population with phenotypic variation and substrate for selection, is occurring at a higher rate in mice relative to humans, and in genomes of comparable size, we considered reasons for the lack of observable clonal expansions (on the phylogenies) and absence of selection on non-synonymous mutations (using dN/dS), both of which manifest ubiquitously over time in human haematopoiesis^9^. One possibility here is that there are insufficient HSC and MPP divisions within the short lifespan of mice to facilitate detectable clonal expansions of cells with fitness-inferring mutations. Secondly, as both population size and the frequency of self-renewing cell divisions (captured in N/λ) determine the rate of random drift, and hence the drift threshold that selection must overcome^42^, the fitness (s) of newly arising mutations may also be insufficient for their carrier subclones to exceed the genetic drift threshold within a mouse lifespan (s=λ/N representing the drift threshold^42^).

In the first scenario, clones under selection (i.e., with necessary driver mutations) will still be present, but would just be too small to detect using a phylogenetic approach that only readily identifies larger clones (>5% clonal fraction). In the second scenario, the fitness landscape of any detectable clones would reflect the specific murine haematopoietic drift threshold. Therefore, we sought to address both questions to better understand the evolutionary processes shaping somatic evolution in blood.

### Positive selection during homeostatic and perturbed murine haematopoiesis

To examine murine blood for very small expanded haematopoietic clones and the presence of clonal haematopoiesis (CH), we employed targeted duplex consensus sequencing (Methods). We reasoned that clonal expansions in mice could be driven by mutations in orthologues of at least some of the same genes that drive human CH due to their evolutionary conserved biological functions. We designed a target panel covering murine orthologues of 24 genes associated with human CH (61.8 kb panel, Methods) and tested whole blood from mice aged 3 to 37 months for mutations. Median duplex consensus coverage per sample ranged from 28,000–41,000X, allowing detection of variants present at the magnitude of 1 in 10,000 cells (Methods, Supplementary Note 3).

We observed expanded CH clones that increased in prevalence with age (Fig.4A,B). Samples from young mice (3 months) displayed infrequent or absent (range 0-1) clones, while those from the oldest mice (37 months) displayed on average 3.5 clones (range 1-6) per animal across these targeted genes. Average clone size was very small at 0.017% of nucleated blood cells (range 0.0036-0.27%) – representing clonal fractions between 1 in 500 to 1 in 30,000 cells. Clonal expansions were recurrently driven by mutations in *Dnmt3a* and *Tet2*, genes frequently mutated in human CH^43^, but also *Bcor* and *Bcorl1*, observed in humans following bone marrow immune insult^44^. These data are consistent with a previous report identifying rare expanded clones in mice following transplant^25^.

**Figure 4.**
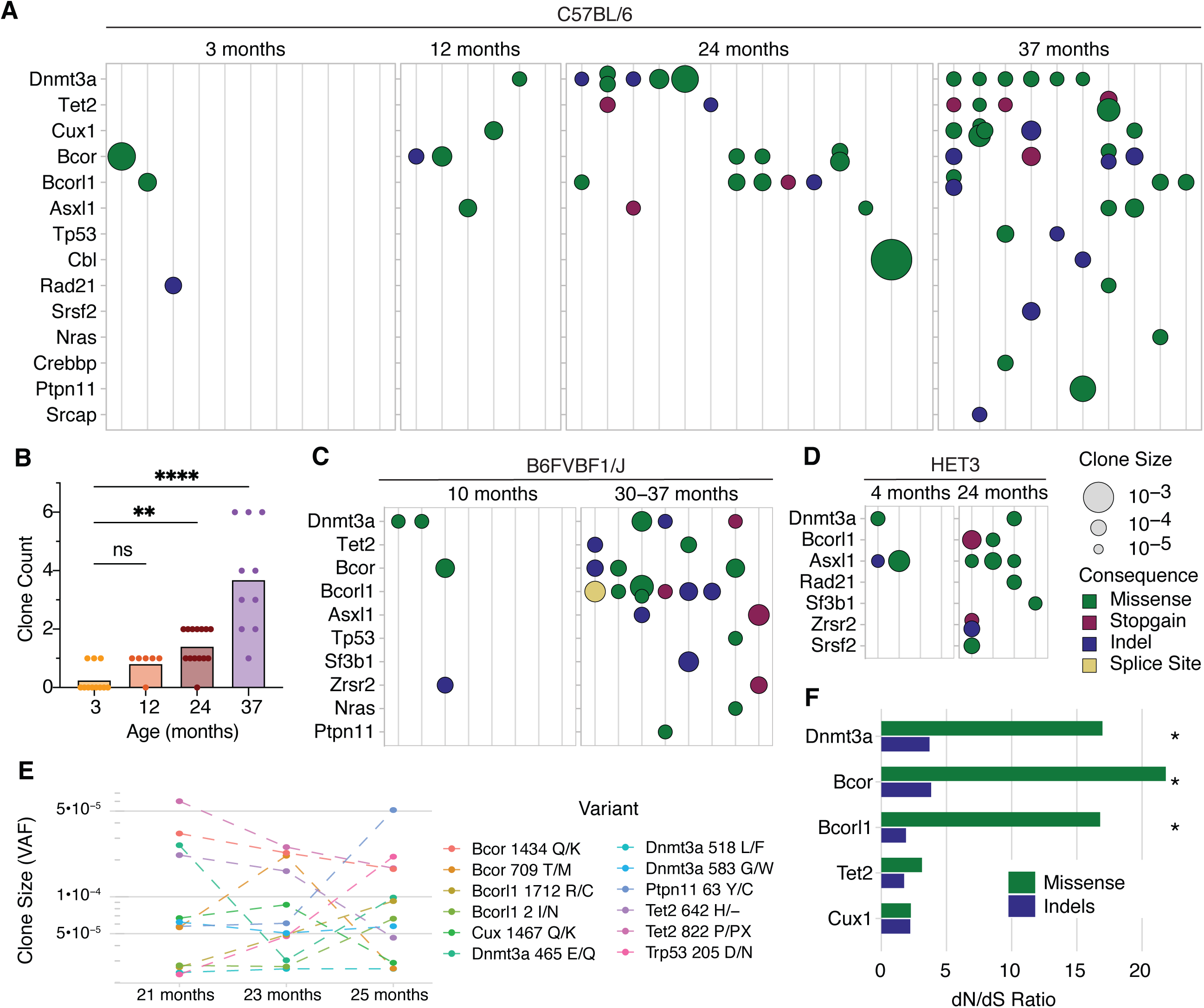
Clonal haematopoiesis during normal ageing in mouse. **A)** Dot-plot describing incidence of clonal haematopoiesis in mice at increasing age. Each vertical column represents a single mouse sample with detected clone size and consequence indicated by dot size and colour. Strain is C57BL/6J. **B)** Barplot summarising clone count per sample as illustrated in A. Differences in clone incidence were quantified by the Kruskal-Wallis test. **C-D)** Murine clonal haematopoiesis incidence in the laboratory strains **C)** B6FVBF1/J (F1 hybrid from crossing inbred C57BL/6J x FVB/NJ), and **D)** HET3 (a four-way cross between C57BL/6J, BALB/cByJ, C3H/HeJ, and DBA/2J). **E)** Clone size changes in samples collected serially over 4 months. Clones are coloured by mutation. **F)** dN/dS ratios for targeted genes mutated in murine clonal haematopoiesis. Variants from all donors in A were used to determine gene level dN/dS ratios. * represents dN/dS >1 with q-value <0.1.

Increased clonal prevalence with age was observed across different laboratory strains, including the genetically heterogeneous HET3 strain, and at similar clonal fractions (Fig.4C,D), confirming that small clones driven by known CH drivers are not specific to the C57BL/6J strain. Clones were present in biological replicates and persistent in mice sampled longitudinally over four months, though individual clonal dynamics varied (Fig.4E). Variants displayed enrichment for nonsynonymous mutations across these genes (dN/dS 2.00, CI 1.01-4.02), with per-gene positive selection evident for *Dnmt3a, Bcor*, and *Bcorl1* (dN/dS>1, q<0.1) (Fig.4F). These data confirm that these small clonal expansions in murine blood are being shaped by positive selection and are not the result of genetic drift.

Laboratory mice are maintained in exceptionally clean conditions with a controlled diet and environment, in contrast to the regular microbial exposures and systemic insults experienced by humans. We considered whether similar exposures, which may accelerate CH in humans^45,46^, could enhance selection and clonal expansion in mice. In humans, mutant *TP53* and *PPM1D* clones are positively selected for in the context of chemotherapy^47–49^, while *BCOR* mutated clones have a fitness advantage in the bone marrow environment of aplastic anaemia^44^. To examine whether the murine haematopoietic selective landscape can be similarly altered, we applied a series of infectious or myeloablative exposures.

We first subjected mice to a normalised microbial experience (NME), in which laboratory mice are infected with common mouse microbes via exposure to fomite (pet store) bedding, resulting in the transfer of bacterial, viral, and parasitic pathogens^50,51^. Such exposure has been shown to drive functional maturation of the murine immune system^50^. Aged NME-exposed mice displayed an increased burden of somatic clones, especially driven by *Trp53* (Fig.5A). As NME exposure transmits multiple types of pathogens, making it challenging to disentangle specific pathogen effects, we next performed targeted exposure to *Mycobacterium avium*, which has been shown to activate HSCs and lead to chronic inflammation^52^. Aged mice chronically infected with *M. avium* showed an increased frequency of *Dnmt3a*, *Bcor, Tet2*, and *Asxl1* mutant clones (Fig.5B), suggesting that clones harbouring these mutations experience a competitive advantage in the context of infectious exposure. Differences in driver mutation prevalence between NME and *M. avium*-infected mice may reflect infection severity or immune response differences.

**Figure 5.**
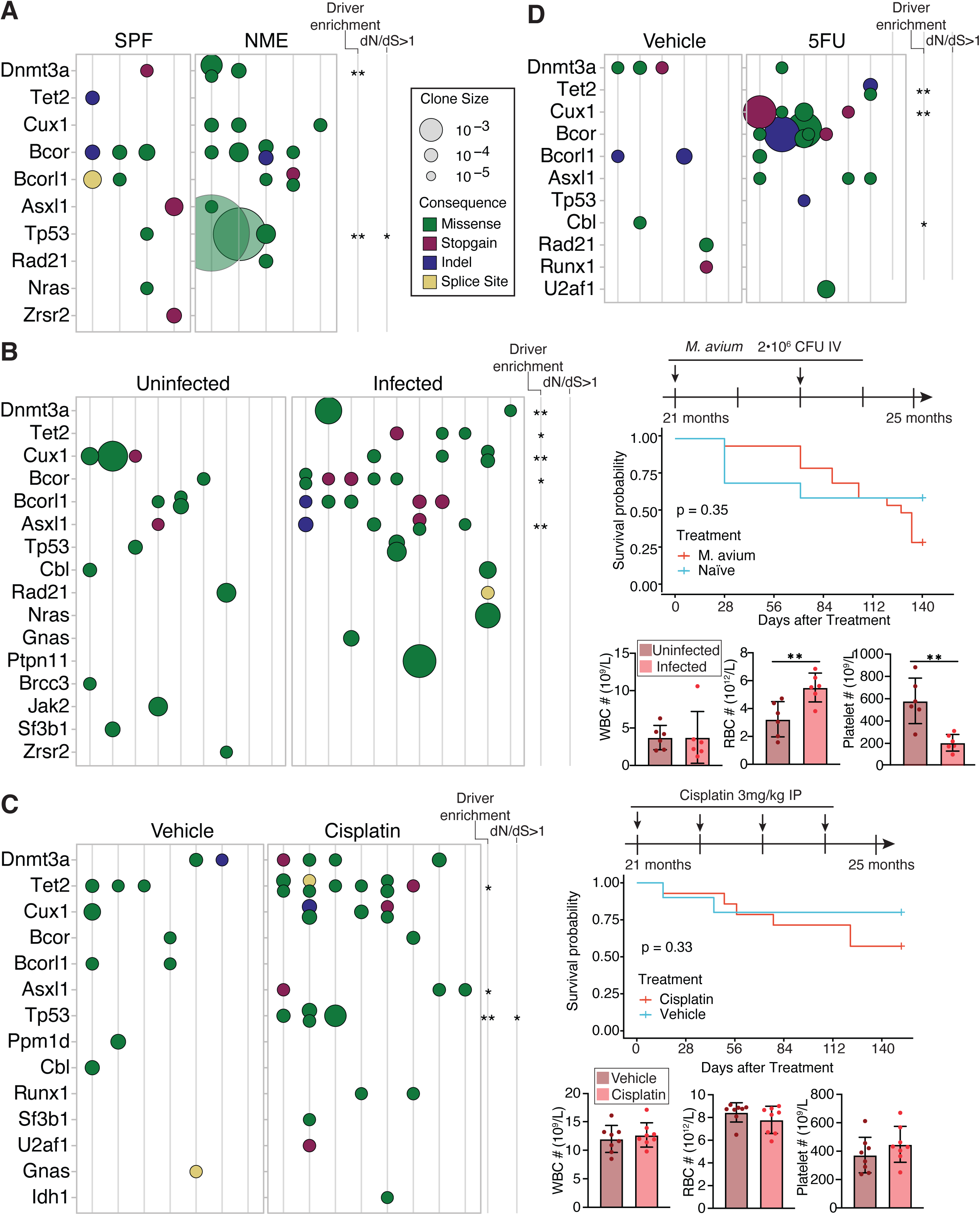
Haematopoietic perturbation modulates selection landscapes Clonal haematopoiesis prevalence in aged mice following. **A)** normalised microbial experience (NME), **B)** *M. avium* infection, **C)** cisplatin treatment, and **D)** 5-FU myeloablation. At final sampling, aged mice were 30-months-old for the NME experiments in panel A), and were 25-months-old for the perturbation experiments in panels B), C), and D). Enrichment of clonal prevalence and dN/dS ratios departing from parity following treatment are shown for each gene. Survival curves and experimental endpoint blood counts are displayed for B) and C), using log-rank and two-sided t tests, respectively. Treatment schedules are as displayed or described in Methods.

To observe the impact of myeloablation, aged mice were treated with commonly used chemotherapeutic agents 5-fluorouracil and cisplatin. When treated with monthly doses of cisplatin, we observed globally increased somatic clonal burden (Fig.5C, p= 0.027). Clones driven by *Trp53*, *Tet2*, and *Asxl1* were enriched relative to controls, and gene-level dN/dS analysis indicated that *Trp53* was under positive selection for nonsynonymous mutations (Fig.5C), analogous to human observations^47–49^. Similarly, aged mice treated recurrently with the chemotherapeutic agent 5-fluorouracil displayed clones at magnitudes-greater proportions than age-matched controls (Fig.5D). Broadly, these data illustrate that haematopoietic mutation accrual and selection are sufficient to drive native CH in mice, with modulable selection landscapes.

### Fitness landscape of clonal haematopoiesis in murine haematopoiesis

Having observed an evolutionarily conserved clonal selection landscapes in murine blood, we wished to understand why observed clone sizes were much smaller (median 0.017%) compared to human CH at equivalent times during lifespan. Therefore, we estimated the fitness landscape of these driver mutations in mouse. We evaluated the distribution of observed variant allele fractions (VAF) from the targeted duplex sequencing, using an established continuous time branching evolutionary framework for HSC dynamics^53^ (Methods). How the observed distribution of VAFs, predicted by the evolutionary framework, changes with age is then used to infer the underlying effective population size (N /λ), mutation rates (μ), and fitness effects (i.e., clonal growth percentage per year) of non-synonymous mutations. Due to increasing N /λ with mouse age (Fig.3A), only clones from mice of a similar age (here chosen to be 24-25 months) during steady-state haematopoiesis were included in the analyses.

By analysing the distribution of neutral mutation VAFs (clones at low VAF bearing synonymous or intronic mutations), we first yielded an independent orthogonal estimate of N/λ (Methods) of approximately 16,500 HSC-years (CI, 11,122-21,836, Fig.6A). This inference generated from targeted sequencing is consistent with that generated from whole genome sequencing and approximate Bayesian computation (ABC) (N/λ 7,918 HSC-years, CI 2,277-20,309). Differences in the estimates for N/λ from ABC versus the branching evolutionary framework^53^ are likely influenced by (i) the ABC method takes into account population growth inferred from the phylogenetic trees, whereas the branching evolutionary framework assumes a stable population size, and (ii) the branching evolutionary framework model relies on using the intronic/synonymous mutation rate as the background for identifying clonal expansions, which may not reflect the genome-wide mutation levels. Across the 61.8 kb panel the synonymous/intronic mutation rate was estimated at 1.8⋅10^-4^ base pairs per year (CI 1.2−2.7⋅10^-4^). We estimate a nonsynonymous mutation rate of 3.4⋅10^-4^ base pairs per year (CI: 2.9−3.9⋅10^-4^), again only considering VAFs below the maximum observed synonymous/intronic VAF (Methods), as clones larger than this could be under the influence of positive selection. The total mutation rate within our targeted panel was thus 5.2⋅10^-4^ per year, which when scaled to total genome size, corresponds to a global mutation rate of 11.77⋅10^-9^ per base pair per year (CI 9.28-14.94⋅10^-9^). Encouragingly, this is similar to the mutation rate directly observed from whole genome sequencing of single cell-derived colonies of 8.29⋅10^-9^ per base pair per year (CI 7.73-8.85⋅10^-9^). This agreement suggests that even these low-VAF clones detected from duplex consensus sequencing represent *bona fide* clonal expansions.

**Figure 6.**
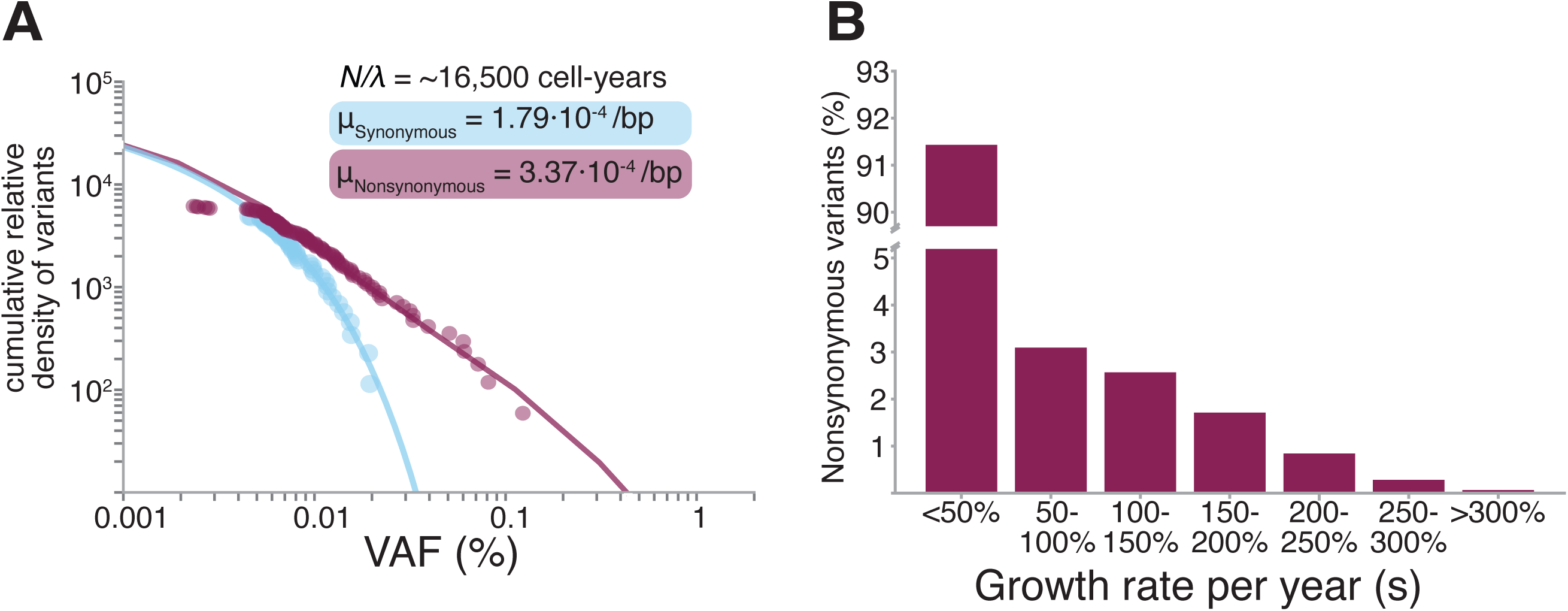
The fitness landscape of known drivers of clonal haematopoiesis. **A)** Reverse cumulative density for all synonymous (including flanking intronic regions in targeted bait set) and nonsynonymous somatic variants detected using duplex sequencing from mice aged 24-25 months, arranged by increased variant allele fraction (VAF). The relative density of synonymous (and flanking intronic) variants, which are assumed to have neutral fitness, yields an estimate for *N*/λ, the ratio of population size and symmetric cell division rate (per year). The synonymous and nonsynonymous mutation rates (μ, base pairs per year) can then be estimated using a maximum likelihood approach. **B)** Distribution of fitness effects for nonsynonymous mutations.

Having inferred N/λ and the non-synonymous mutation rate, we could estimate the distribution of fitness effects driven by non-synonymous mutations (Methods). Our analysis suggests that ∼7% (CI 5-21%) have strong fitness effects (50-200% growth per year) (Fig.6B). Considering that we infer mouse stem cells to be self-renewing roughly every 6 weeks (CI 2.3-12.5 weeks), an *annual* growth rate of 200% translates to a *per* symmetrical self-renewing division selective advantage of ∼15% (5-30%), in line with reported selection coefficients of mutated genes associated with CH in humans^9,34,53^. Indeed, in the short-lived mouse, variants with weaker fitness (<50%) might have insufficient time to enter exponential, deterministic growth within the population, given that clones are not established until *t*_*years*_ > 1/s (ref. ^54^), although any background growth in population size could circumvent this, allowing for weaker variants to fix in the population. This may also explain why some of the low-VAF clones identified by duplex consensus sequencing did not increase in clone size over time (Fig.4E).

## DISCUSSION

Here, we study the ontogeny, population dynamics and somatic evolution of haematopoietic stem cells in the most widely used mammalian model organism, the laboratory mouse. Classical models of blood production depict HSCs at the very top of the haematopoietic differentiation hierarchy, beneath which all blood cell types emanate. Recent studies suggest additional heterogeneity at the top of this haematopoietic hierarchy and nuanced self-renewing dynamics^2–4^. Our phylogenetic data suggest that MPPs (distinguished by their lack of expression of the CD150 marker^55^) do not always arise from HSCs, and that both populations are established during embryogenesis, following which they independently self-renew throughout murine life. That MPPs are noticeably generated from HSCs in a transplant setting may underscore the difference between their *potential* in an experimental setting and their steady-state *in vivo* function. Moreover, lymphoid and myeloid progenitors appear to equally derive from HSC and MPP lineages. Recapture of shared variants indicate both MPPs and HSCs contribute to differentiated peripheral blood production, with a slight bias toward production from ancestral MPP lineages. These data are aligned with lineage tracing that showed both populations are capable of making all cell types during normal life^2,32,56^, and with recent reports of MPPs derived from the embryo (eMPPs) that contribute to lifelong haematopoiesis^2,57^.

We show that HSCs and MPPs grow in lockstep over life, with indistinguishable clonal dynamics and proliferation rates, to reach a combined population of 25,000-100,000 cells, remarkably close to estimates of the human HSC pool size (20,000-200,000 stem cells)^9,10^ and reminiscent of suggestions of conservation of stem cell numbers across mammalian species^58^. Considering the log-fold difference in body mass and consequent demands on blood production, this similarity may be surprising. In both organisms, but especially the mouse, the number of stem cells far exceeds the apparent lifetime need; the stem cell compartment of a single mouse can be used to fully reconstitute the blood of ∼50 transplant recipients^59^. Perhaps a large stem cell pool confers an evolutionary advantage in the face of naturally occurring exposures to environmental pathogens^60^ and tissue injury, through both increased tolerance of stem cell losses and improved adaptation afforded by somatically acquired genomic and epigenetic diversity.

Somatic mutation rates have recently been shown to scale inversely with mammalian lifespan. In colonic epithelium, mice accumulate mutations 20 times faster than humans, aligned with the difference between their lifespans^23^. This observation raises the intriguing possibility that somatic mutation rates are visible to selection through their effects on ageing and lifespan. Our data show this pattern does not extend to blood - the murine HSC mutation rate is only two-fold higher than human^9–11^ despite a 35-fold shorter lifespan, suggesting that mutation accrual patterns across species are under tissue-specific evolutionary constraints. Indeed, somatic mutation rates in germline cells are lower in mouse than in human^61^ and under the influence of distinct factors such as effective population size and age of reproductive maturity^62^. In blood, it is plausible that a low somatic mutation rate is required to minimise the entry of detrimental disease-causing mutations, which when combined with a large stem cell pool, may also reduce the fixation probability of any such mutations. Alternatively, it is also possible that the mutation rate may not reflect haematopoietic adaptation in the mouse but rather a historical evolutionary constraint or a feature of phylogenetic legacy^63^.

Patterns of somatic evolution in humans provide one plausible mechanism by which ageing phenotypes occur. The presence of clonal expansions in elderly human blood driven by somatic mutations is associated with diseases of ageing. However, in the laboratory mouse, which also displays phenotypes of ageing including increased cancer incidence^64^, we only observe small mutation-driven clonal expansions in blood by the end of life, suggesting that any role age-associated haematopoietic oligoclonality plays in human ageing is unlikely to be shared by the laboratory mouse. The dramatically different population structures of haematopoiesis in the old mouse versus old human, together with the small clones (necessitating sensitive detection methods) are crucial factors to be considered when using murine models for future studies of natural CH or haematopoietic ageing. Alternative model organisms, such as non-human primates, display similar stem cell cycling behaviour^65,66^ to humans and larger age-related clonal expansions^67,68^, and thus may be suited to evaluate aspects of native hematopoietic dynamics across the lifespan.

Native murine clones do expand upon systemic exposures and recapitulate patterns previously observed in correlative humans studies^47,48^ and in exposures administered following murine transplant^69–72^ (reviewed in depth in refs. ^17,45,46^). We postulate that the size of clonal expansions is constrained in mouse due to infrequent HSC self-renewing divisions during homeostatic conditions. Our data fit with mouse stem cells self-renewing every six weeks (1.8-13.2 weeks), within the broad range of previous estimates from once in 4 to 24 weeks^73–75^. Whilst this is more frequent than human HSCs (estimated to divide at 1-2 times a year), for patterns of oligoclonality in humans to be recapitulated in the much shorter-lived mouse via genes conferring similar fitness advantages, stem cells would need to self-renew much more frequently. It is possible that mouse strains thought to have higher HSC turnover^76^, or maintained for longer periods in more “wild”-like microbial environments, would exhibit higher levels of native CH. Additional studies to characterise such strains and environments would be of interest.

Nevertheless, our data highlight conserved selection landscapes in mouse with detectable CH in both homeostatic haematopoiesis and under stress when using highly sensitive sequencing. With our observation of evolutionarily conserved constraints on population dynamics of blood, together these drive a distinct pattern of somatic evolution over the murine lifespan. These data provide a framework for the interpretation of future studies of haematopoietic stem cell biology and ageing using the laboratory mouse.

---

**Extended Data Figure 1.**
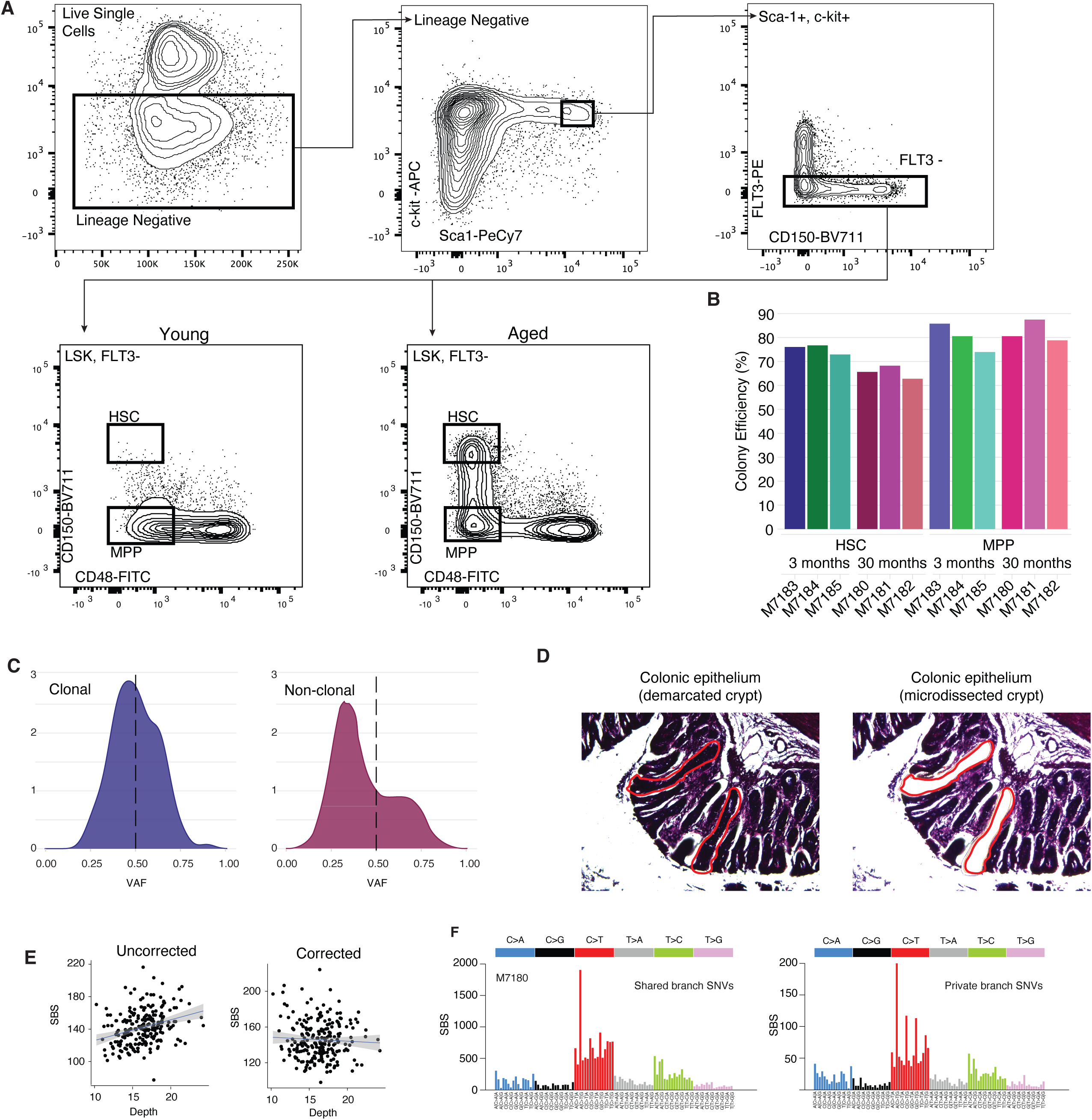
Cell isolation strategy and quality control **A)** Sorting strategy for single HSCs and MPPs from young and aged mice. Progenitor-enriched bone marrow was stained as described in the Methods, and then single cells were sorted into individual wells for *in vitro* expansion. **B)** Colony-forming efficiency of sorted HSCs and MPPs for each sample. Each bar represents the listed cell type and underlying sample ID. **C)** Variant allele fraction (VAF) distribution of all variants within a colony that pass filtering, shown for a representative clonal colony that passed sample QC (left) and a non-clonal colony that passed sample QC (right). After variant filtration, the VAF distribution of a colony’s variants is centred around 50% in clonal colonies, but less than 50% in non-clonal colonies. **D)** Representative image of two colonic crypts isolated by laser capture microdissection. **E)** Correlation between total single base substitution burden and depth, for all colonies from sample M7180, shown before (left) and after (right) sequencing depth correction. **F)** Trinucleotide spectra from aggregated somatic mutations mapped to shared (truncal) or private branches of phylogenetic trees. Signatures are highly similar, suggesting artefacts are not relatively enriched in either portion of reconstructed trees.

**Extended Data Figure 2.**
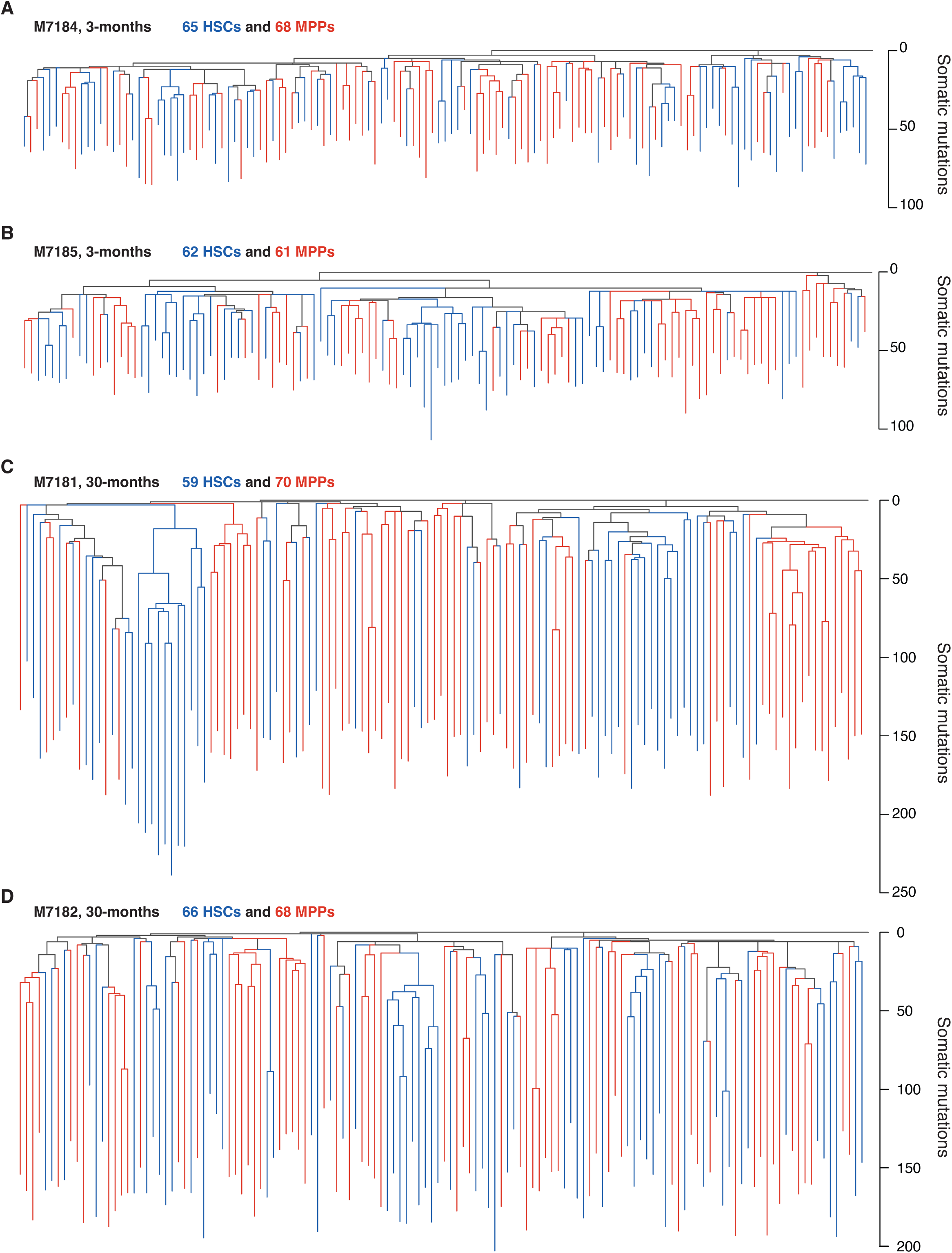
Additional phylogenetic trees from young and aged mice. Phylogenies for **A-B)** 2 additional young (3-month) mice and **C-D)** 2 additional aged mice (30-month), presented as described in Figure 2.

**Extended Data Figure 3.**
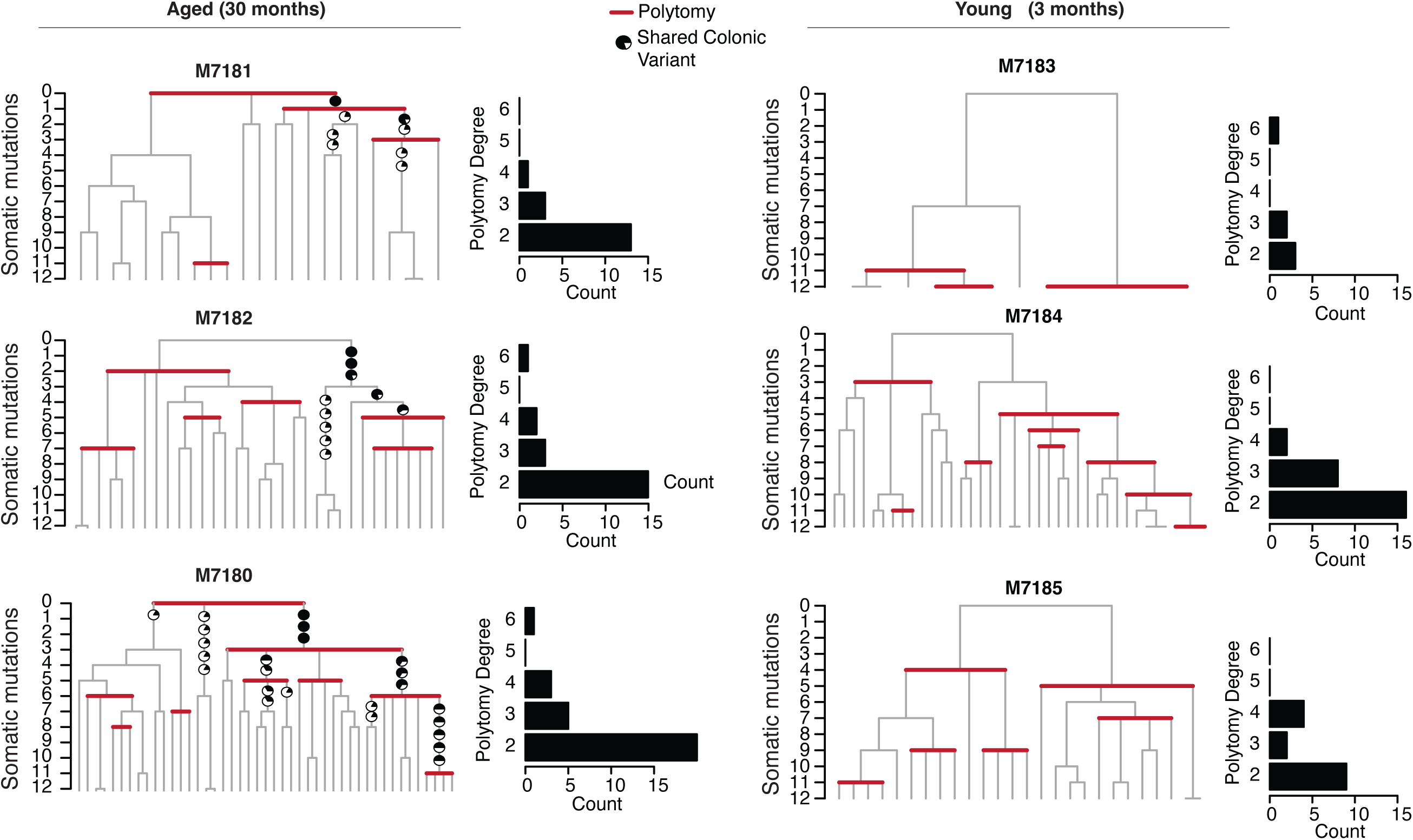
Early-in-life phylogenetic patterns and cross-tissue mutations. Phylogenies from aged (left) and young (right) HSCs zoomed into the first 12 mutations molecular time. Polytomies in the branching structure, which represent cell division without mutation acquisition, are enriched among early-in-life cell divisions at the tops of the phylogenies. Variants shared with matched colonic crypts are layered onto the trees as pie charts. Pie chart fullness represents the proportion of colonic crypts in which the mutation present on the haematopoietic phylogeny was observed. Sample M7183 lacked sufficient early life diversity (<10 unique lineages within 12 mutations molecular time) and thus was excluded.

**Extended Data Figure 4.**
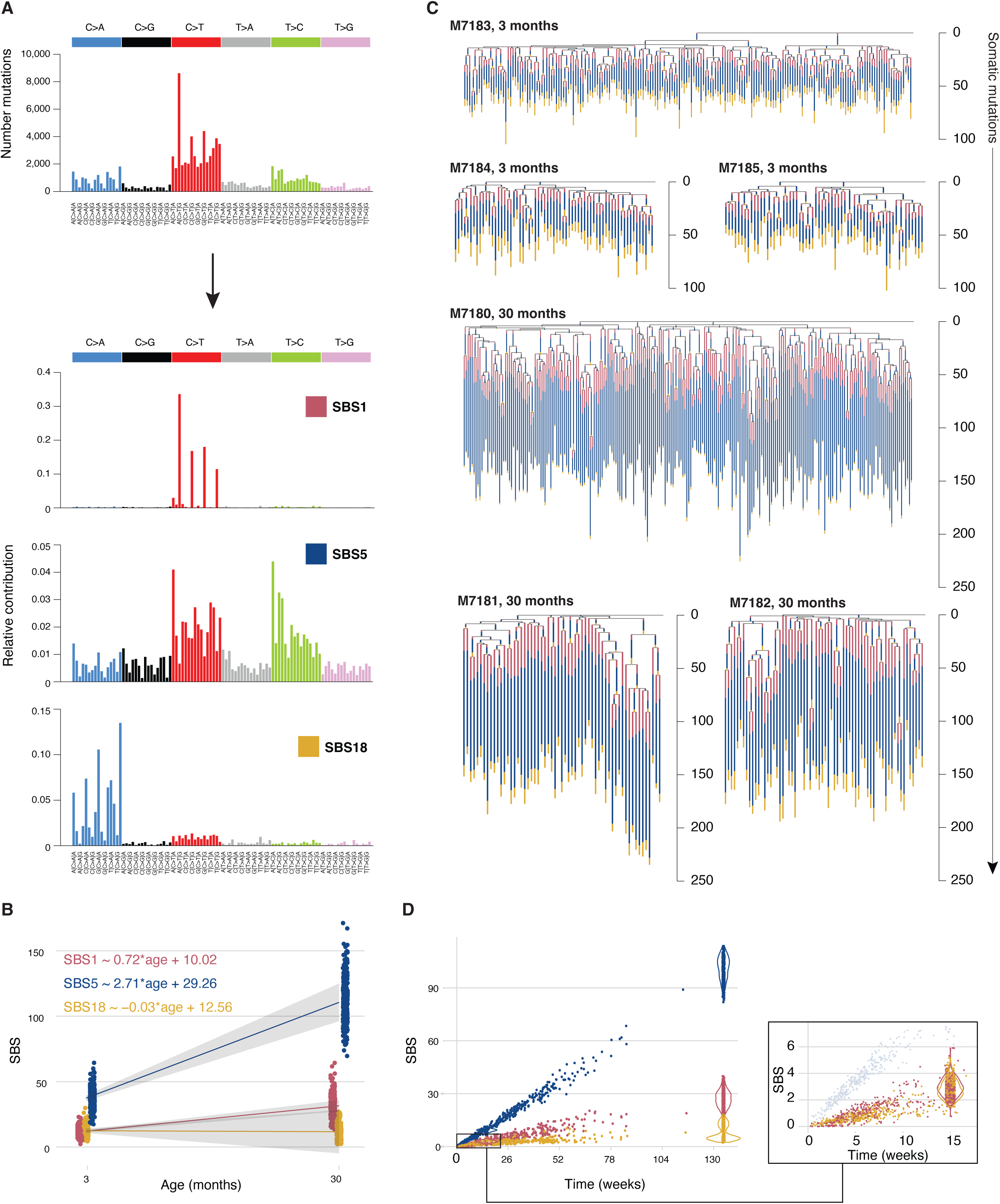
Mutational processes in murine stem cells **A)** Signature extraction overview. Trinucleotide spectra from all single-base substitutions (SBS) (top), were used for signature extraction as described in the Methods. Three signatures identified as SBS1, SBS5, and SBS18 best described the catalogue of mutations observed (cosine similarity=0.997). **B)** Linear mixed-effect regression of signature-specific mutation burdens observed in colonies. Shaded areas indicate the 95% confidence interval. **C)** Signature attribution in phylogenies. Individual branches of HSC phylogenies are overlaid with signature contribution proportions. SBSs assigned to each branch were fit to SBS1, SBS5 or SBS18. **D)** Signature-specific mutation accumulation in all branches across phylogenies. Early-life branchpoints, located at the top of a given phylogeny, and shown as an inset.

**Extended Data Figure 5.**
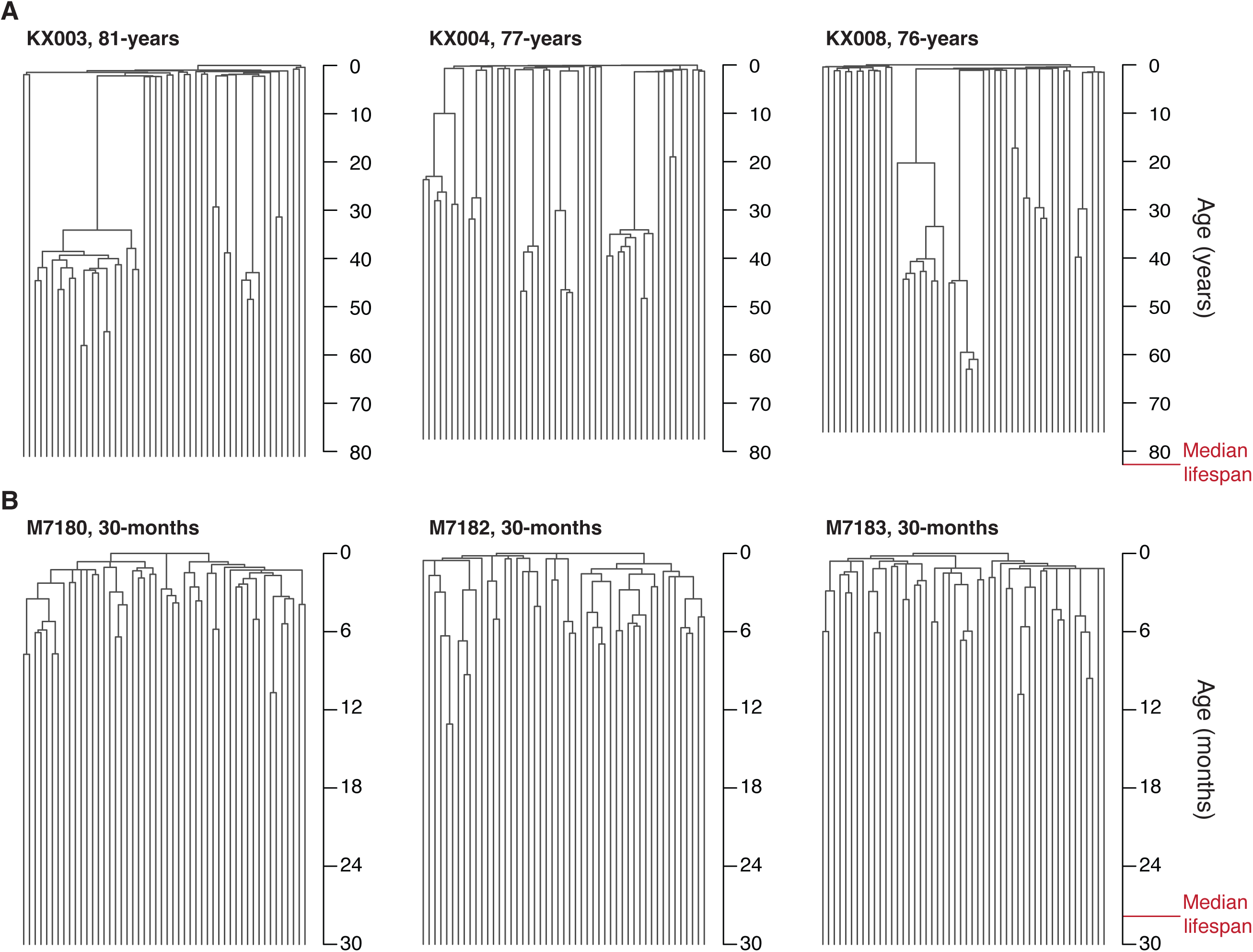
Phylogeny comparison between aged human and mouse **A)** Representative ultrametric phylogenies from the three oldest humans described in Mitchell *et al*.^9^ The published trees have been randomly downsampled to 100 colonies (tips). **B)** Aged mouse phylogenies, also downsampled to 100 colonies, to allow comparison of topological structure. The median lifespan for human and mouse species are labelled and were derived as described in Supplementary Note 1. Full murine phylogenetic trees are shown in Figure 2A-B and Extended Data Figure 2).

**Extended Data Figure 6.**
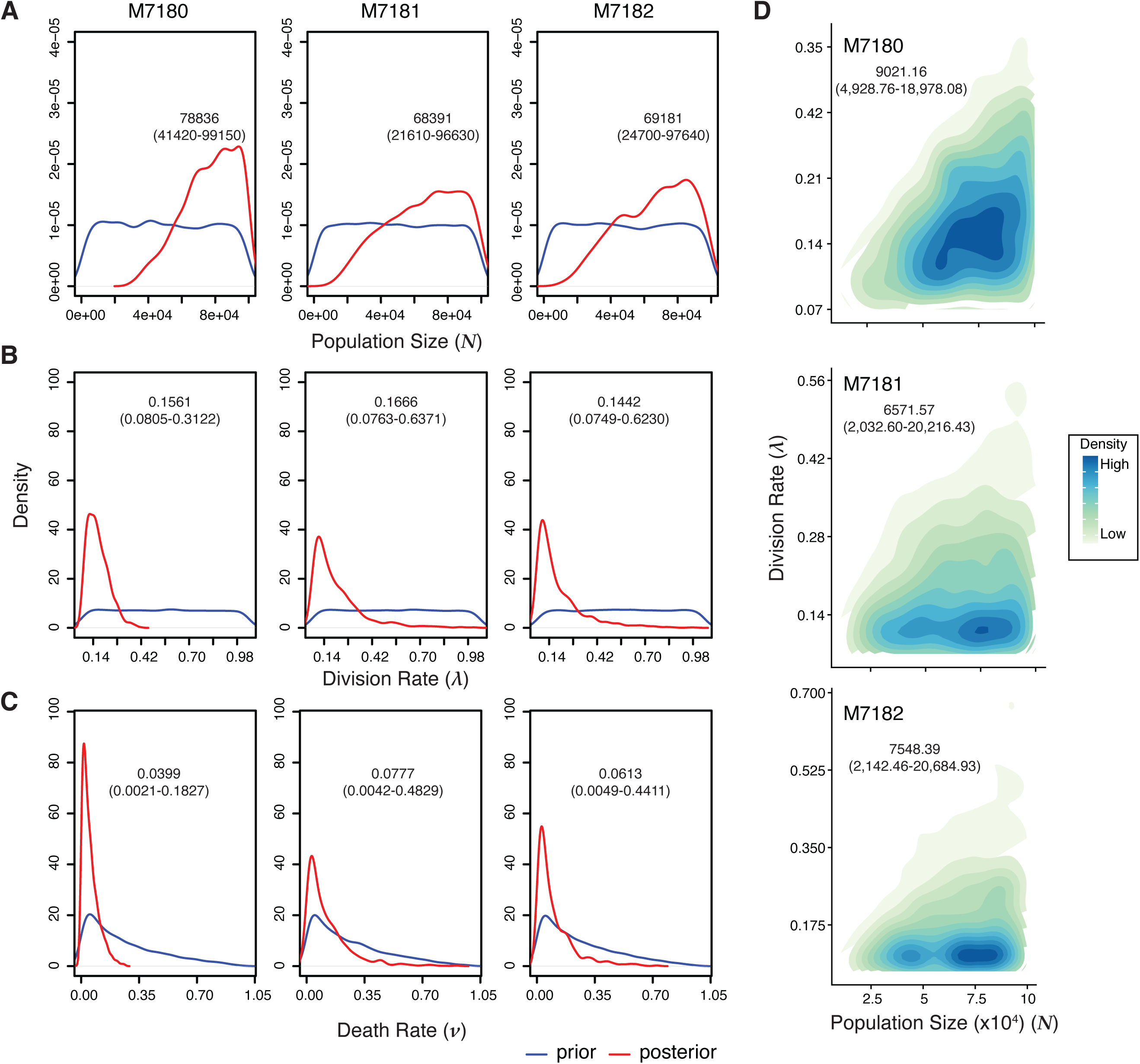
Approximate Bayesian inferences Results from approximate Bayesian computation (ABC) inference of **A)** population size (*N*), **B)** symmetric division rate per week (λ), and **C)** death rate per week (*v*) for the three 30-month-old mice. Blue lines represent the prior density of parameters; red lines represent the posterior densities. Median posterior density estimates and 95% credibility intervals are displayed for each parameter per sample. The prior density for the death rate was bounded to ensure the growth rate (λ − *v*) remained positive, as observed in *phylodyn* trajectories in Figure 3. **D)** Joint density distributions indicating optimal parameters of population size and division rates that explain observed phylogenetic trees. The estimated N/λ, in HSC-years, is shown with 95% credibility intervals. Data from the three aged mice are shown.

**Extended Data Figure 7:**
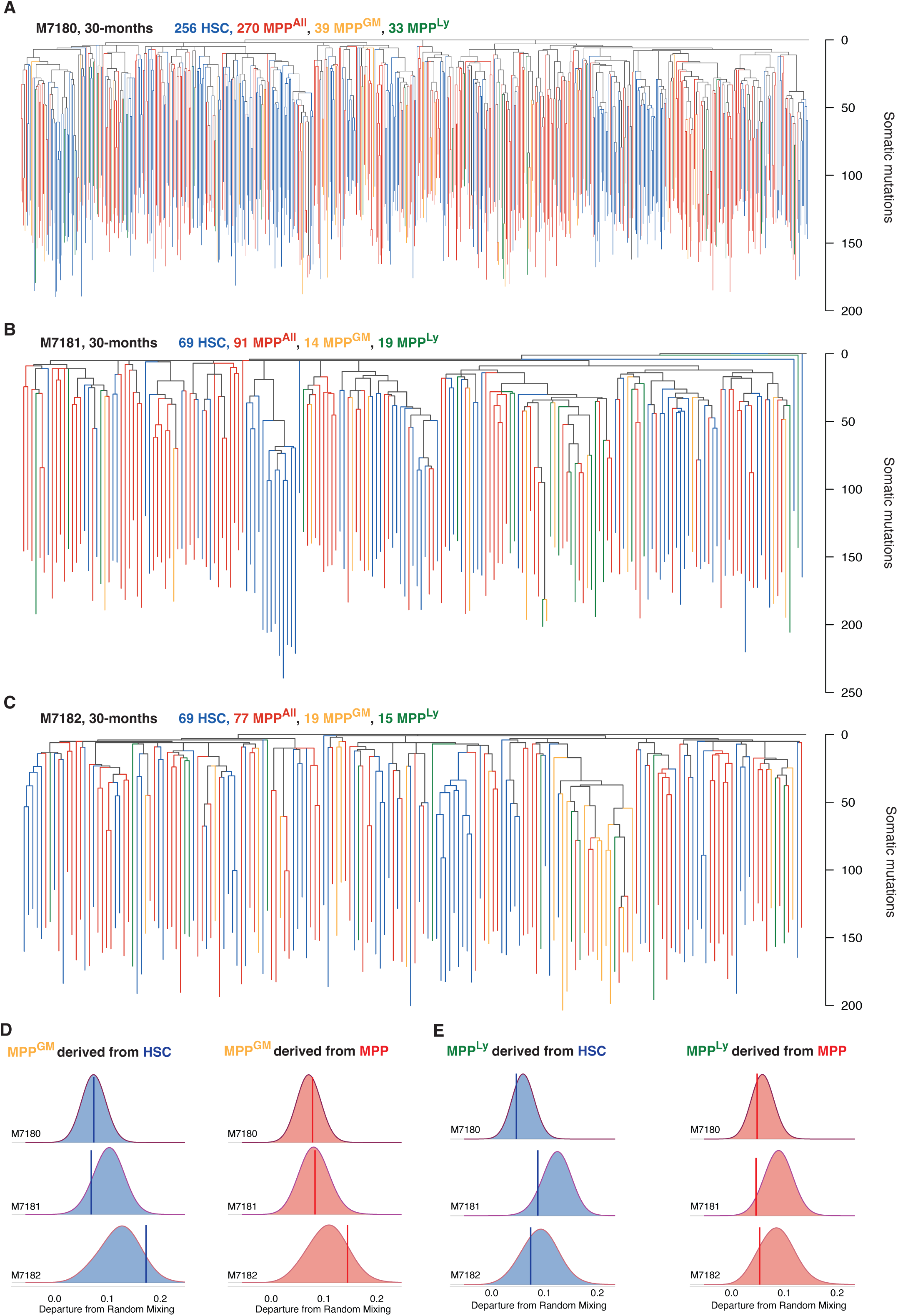
Extended phylogenetic trees (HSC, MPP, and early progenitor). **A-C)** Extended phylogenies were created for three 30-month mice using the pattern of sharing of somatic mutations among HSCs (blue), MPPs (red), and the mixed LSK (Lineage-, Sca1+, c-kit+) hematopoietic progenitor compartment. The LSK compartment contains HSCs and MPP, and additionally contains the myeloid-biased MPP^GM^ (orange) and lymphoid-biased MPP^Ly^ populations (green). LSK subcompartments were identified at time of single cell sorting using a consensus definition^55^. Each tip represents a single colony. Branch lengths represent mutation numbers. **D-E)** Clade mixing metrics for MPP^GM^ and MPP^Ly^ colonies used to evaluate interrelatedness with HSC and MPP. HSC, MPP and MPP^GM^ or MPP^Ly^ were designated as being in the same clade if they share a most recent common ancestor after 25 mutations, corresponding to early foetal development. Only clades with more than 3 colonies are considered. The vertical bar reflects the average clade mixing metric observed in the constructed phylogenies, while distributions reflect the average clade mixing metric expected random chance, estimated by reshuffling the tip states. If the observed value (vertical bar) significantly deviated from random chance (filled distribution), then there would be minimal overlap between the observed data and the random reshuffling distribution. The average clade mixing metric for MPPGM compared to HSCs (blue) and MPPs (red) is shown in **D)**. The similar measure of interrelatedness of MPPLy to HSCs and MPPs is shown in **E)**.

**Extended Data Figure 8:**
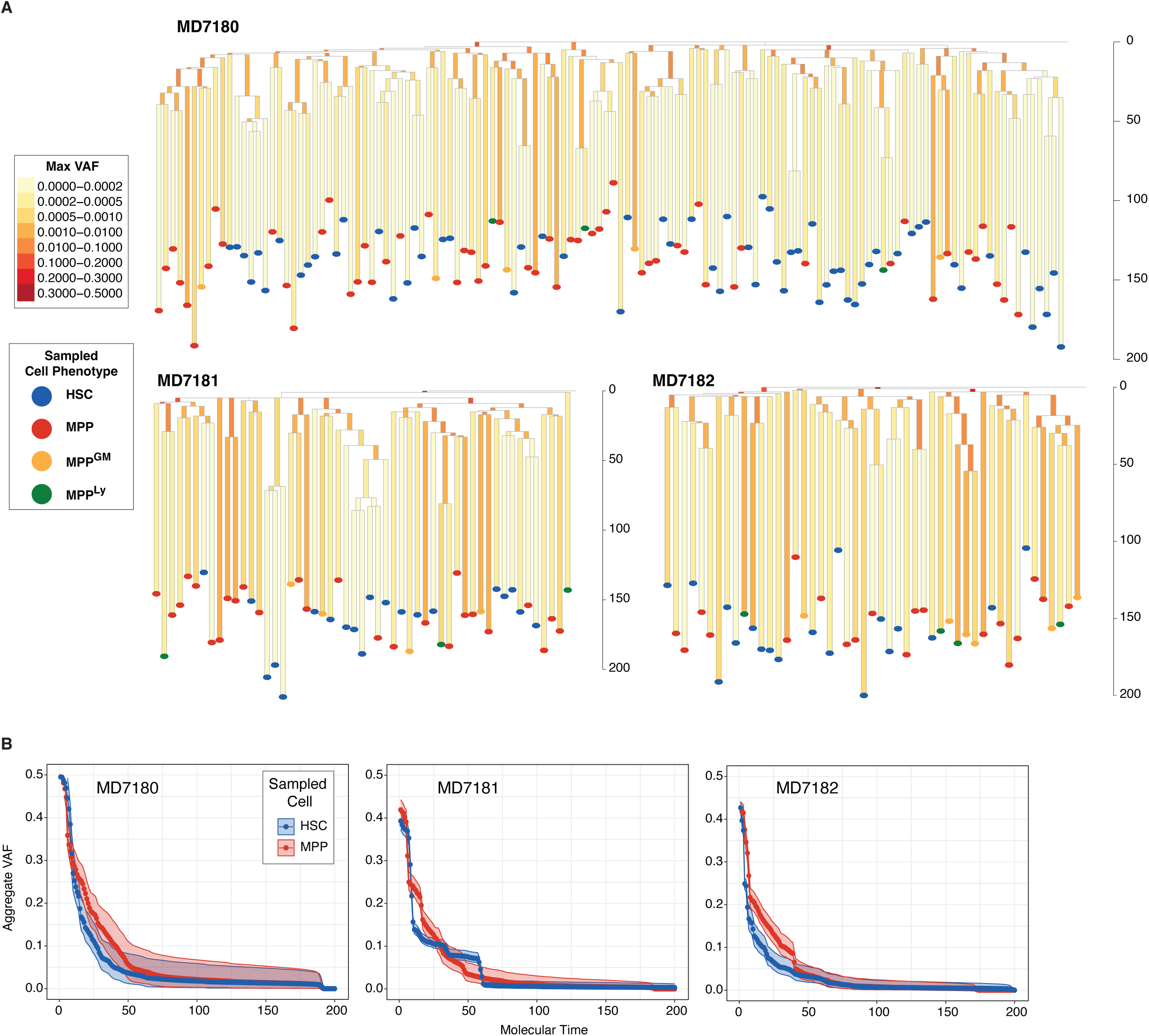
Mutation overlap between phylogenies and peripheral blood. **A)** Phylogenies for three aged mice (as described in Extended Data Figure 7A-C) constructed to only include private branches targeted with the peripheral blood baitset. Branch shading indicates the maximum VAF among branch-specific variants captured in peripheral blood. The sampled cell immunophenotype is indicated by dot colour at the bottom of each private branch. **B)** VAF trajectories of HSC and MPP variants shared in peripheral blood. The aggregate VAF across molecular time is calculated using Gibbs sampling (Methods). Earlier molecular time corresponds to further in the ancestral past. Shaded regions denote 95% confidence intervals of the VAF estimates.

**Extended Data Figure 9:**
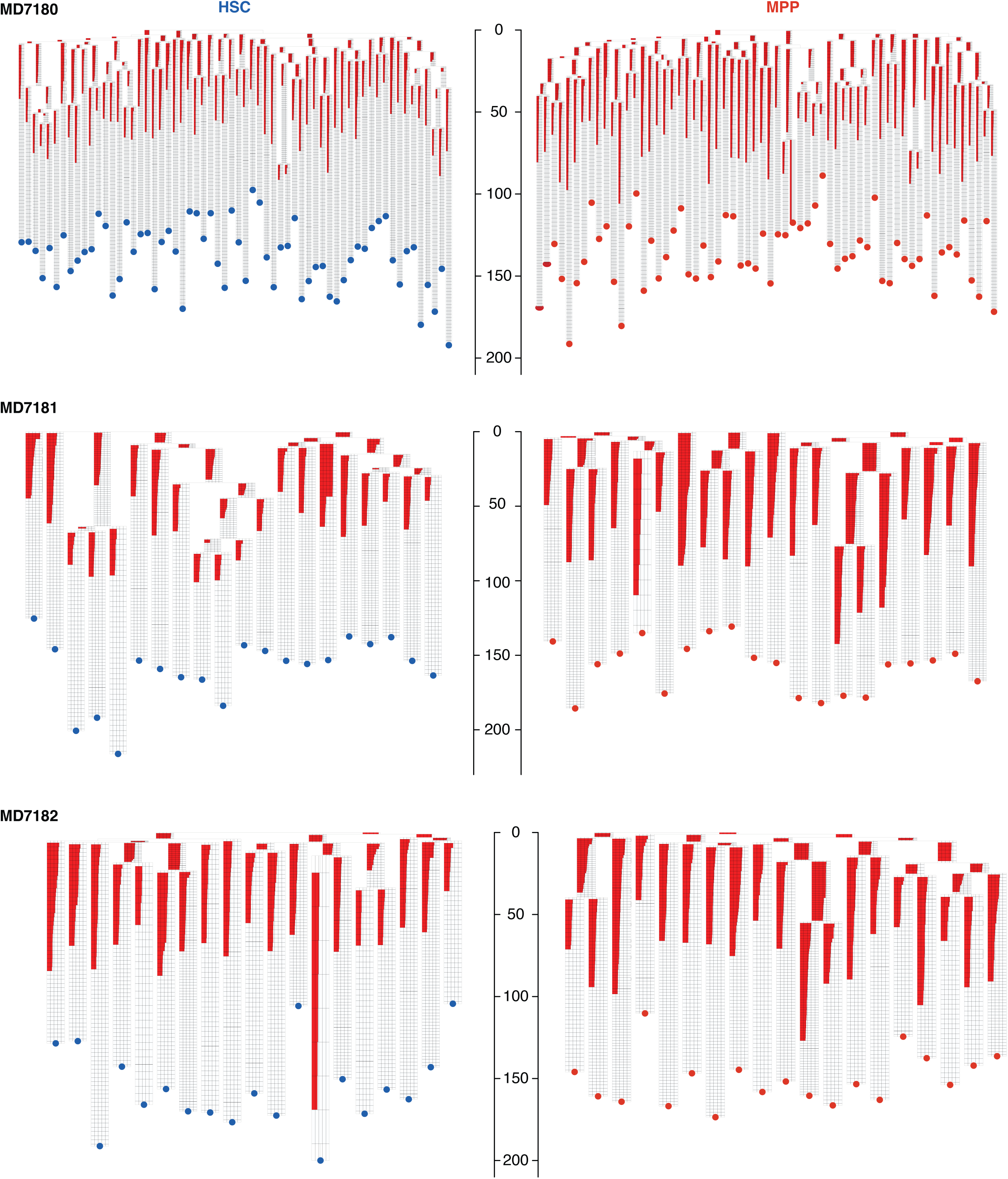
Peripheral blood VAF of variants shared with HSCs and MPPs. Baitset mutation-specific HSC and MPP phylogenies are shown for each 30-month mouse. Each branch shows mutations that were detected in peripheral blood in descending VAF order. On each branch, a row denotes a single variant mapped to that specific branch. Red fill denotes the peripheral blood VAF for the variant. VAF is denoted on a log scale from 10-5 to 1; internal divisions are marked from left to right at VAF 0.0001, 0.001, 0.01, and 0.1. HSC trees are shown on the left with blue dots at terminal branches; MPP trees are shown on the right with red dots. Trees are downsampled to allow equivalent comparison between HSC and MPP branches. Only variants seen in peripheral blood with a depth > 100X are shown.

## Acknowledgements

C.D.K. was supported by F30DK131638. The Goodell lab is supported by grants from the National Institutes of Health, including AG036695, CA183252, CA237291, DK092883, 1P01CA265748, F30HD111129 (SW), and the Milky Way Research Foundation. J.N. is supported by a Cancer Research UK Advanced Clinical Fellowship and work in the Nangalia lab is supported by Wellcome, Cancer Research UK, Alborada Trust, Rosetrees Trust and the MPN Research Foundation. LJN is supported by NIH AG063543 and AG056278. The Niedernhofer lab is supported by AG063543 and AG063543-02S1.The Harrison lab is supported by 5U01AG022308. DL, MAF, and KYK were supported by R35 HL155672 (KYK), R01 AI141716 (KYK), F31 HL154661 (DL), F31 HL156500 (MF), and a minority graduate fellowship from the American Society of Hematology (MF). KN is supported by the Wellcome Trust and CRUK. We are grateful for the assistance of Ryan D. O’Kelly and Mark Pierson in conducting NME experiments. We thank Elisa Laurenti, Stephen Loughran and Tony Green for constructive discussions and J. Thomas Gebert and Hilda L. Chan for the critical feedback.

## Author contributions

CDK, JN and MAG designed the experiments. JN and MAG supervised the project. CDK performed cell sorting and *in vitro* culture with support from SW, RA, AM, AG, EM. CDK performed genomic, phylogenetic, signature, and population dynamics analyses with support from NW, KJD, DL, JF, EM, PJC, PG, JN. NK developed hidden Markov modelling and KJD performed population dynamic inferences. AC and KN prepared colonic crypt microdissections. CDK performed mouse experiments with advice and assistance from MJY, SW, KN, AC, DL, MAF, RA, AM, AG, DH, KYK, LJN. CJW and JRB developed the population genetic analyses of clone sizes and parameter inferences in Figure 6. CDK, JN and MAG wrote and edited the manuscript. All authors reviewed and edited the manuscript.

## Database accession

Whole genome sequencing data will be deposited at the European Nucleotide Archive at accession numbers ERP138320 and ERP144323. Targeted duplex sequencing data will be deposited at NCBI BioProject PRJNA1033340.

## METHODS

### Cohort

Wild-type C57BL/6 mice were bred at Baylor College of Medicine or received from the Aged Rodent Colony at the National Institute of Aging (Baltimore, MD). C57BL/6J:FVB/NJ F_1_ hybrid mice were bred in the Niedernhofer laboratory at the University of Minnesota as previously described^77^. HET3 mice were bred at the Jackson Laboratories as described^78^. C57BL/6 were housed at the AALAC-approved Center for Comparative Medicine in BSL-2 suites. Experimental procedures were approved by the Baylor College of Medicine or University of Minnesota Institutional Animal Care and Use Committees and performed following the Office of Laboratory Animal Welfare guidelines and PHS Policy on Use of Laboratory Animals.

### Hematopoietic progenitor purification

Whole bone marrow (WBM) cells were isolated from murine hindlimbs and enriched for c-Kit+ hematopoietic progenitors prior to fluorescence-activated cell sorting (FACS) using a BD Aria II. WBM was incubated with anti-CD117 microbeads (Miltenyi Biotec) for 30 minutes at 4C following my magnetic column enrichment (LS Columns, Miltenyi Biotec). Progenitor-enriched cells were stained with an antibody cocktail to identify specific progenitor populations using a recent consensus definition^55^. LSKs, containing a mixture of stem and progenitor cells, were defined as Lineage^-^ckit^+^Sca-1^+^ (Lineage^-^ refers to being negative for expression of a set of lineage-defining markers indicated below). HSCs were defined as LSK^+^FLT-3^-^CD48^-^CD150^+^; MPPs were defined as LSK^+^FLT-3^-^CD48^-^CD150^-^. MPP^GM^ was defined as LSK^+^FLT-3^-^CD48^+^CD150^-^ and MPP^Ly^ was defined as LSK^+^FLT-3^+^CD150^-^. The gating strategy is illustrated in Extended Data Fig.1A. This immunophenotypic HSC population includes long-term stem cells with serial repopulating ability, while the MPP population is limited to short-term repopulation, as demonstrated in transplantation assays^26–31^. For sorting HSCs from newborn pups, the lineage marker Mac1 was excluded because it is known to be highly expressed on foetal HSCs^79^. Antibodies were c-kit/APC, Sca1/Pe-Cy7, Flt-3/PE, CD48/FITC, CD150/BV711, Lineage (CD4, CD8, Gr1, Mac1, Ter119)/Pacific Blue and purchased from BD Biosciences or eBioscience.

### Single-cell haematopoietic colony expansion *in vitro*

Cell sorting was performed on a BD Arial II in two stages. First, HSCs and MPPs were sorted into separate tubes containing ice-cold FBS using the “yield” sort purity setting to maximise positive cells. Second, the cell populations from stage one were single-cell index-sorted into individual wells of a 96-well flat bottom tissue culture plates containing 100uL of Methocult M3434 medium (Stem Cell) supplemented with 1% penicillin-streptomycin (ThermoFisher). No cytokine supplements were added to the base methylcellulose medium. Cells were incubated at 37⁰C and 5% CO_2_ for 14±2 days, followed by manual assessment of colony growth. Colonies (>200 cells) were transferred to a fresh 96-well plate, washed once with ice-cold PBS, then centrifuged at 800xg for 10 minutes. Supernatant was removed to 10-15 µL prior to DNA extraction on the fresh pellet. The Arcturus Picopure DNA Extraction kit (ThermoFisher) was used to purify DNA from individual colonies according to the manufacturer’s instructions. 62-88% HSCs and MPPs produced colonies, indicating we are sampling from representative populations within each individual compartment (Extended Data Fig.1B). Extracted DNA from each colony was topped off with 50ul Buffer RLT (Qiagen) and stored at −80⁰C.

### Laser capture microdissection

Matched colonic tissue from the three 30-month-old mice used in this study was dissected and snap frozen at the time of bone marrow harvest. Colon tissue sectioning and laser capture microdissection (LCM) was performed as previously described^80^. Briefly, previously snap frozen colon tissue was fixed in PAXgene FIX (Qiagen) at room temperature for 24 hours and subsequently transferred into PAXgene Stabilizer for storage until further processing at −20⁰C. The fixed tissue was then paraffin-embedded, cut into 10 μm sections, and mounted on PEN-membrane slides. Staining of histology sections was done using haematoxylin and eosin as previously described^23^, with scans of each section captured thereafter. Individual colonic crypts were identified, demarcated, and isolated by LCM using a Leica Microsystems LMD 7000 microscope (Extended Data Fig.1D) followed by lysis using the Arcturus Picopure DNA Extraction kit (ThermoFisher).

### Whole genome sequencing

For low DNA input whole genome sequencing of haematopoietic colonies (from young and aged mice) and colonic crypts (from aged mice), enzymatic fragmentation-based library preparation was performed on 1-10 ng of colony DNA, as previously described^80^. Whole genome sequencing (2×150 bp) was performed at a median sequencing depth of 14X for haematopoietic colonies and 17X for colonic crypts on the NovaSeq platform. Reads were aligned to the GRCm38 mouse reference genome using bwa-mem. For whole genome single molecule (nanorate) sequencing, we used matched whole blood genomic DNA collected from the three aged mice during tissue harvest. Nanorate sequencing library preparation was performed as previously described^12^, followed by sequencing to 146-153X coverage on the Illumina Novaseq platform.

### Somatic mutation identification and quality control in haematopoietic colonies

Single nucleotide variants (SNVs) in each colony were identified using CaVEMan^81^, including an unmatched normal mouse control sample that had previously undergone whole genome sequencing (MDGRCm38is). Insertions and deletions were identified using cpgPindel^82^. Filters specific to low-input sequencing artefacts were applied^80^. As variant calling utilised an unmatched control, both somatic and germline variants were initially called. Germline variants and recurrent sequencing artefacts were then identified using pooled information across mouse-specific colonies and filters as follows: i) *Homopolymer run filter.* To reduce artefacts due to mapping errors or introduced by polymerase slippage, SNVs and indels adjacent to a single nucleotide repeat of length 5 or more were excluded. ii) *Strand bias filter.* Variants supported by reads only in positive or negative directions are likely artefacts. For SNVs, a two-sided binomial test was used to assess if the proportion of forward reads among mutant allele-supporting reads differed from 0.5. Any variant with significantly uneven mutant read support (cutoff of p<0.001) and with over 80% of unidirectional mutant reads were excluded. For each indel, if the Pindel call in the originally supporting colony lacked bidirectional support, the indel was excluded. *iii) Beta binomial filter.* Variants were filtered based on a beta-binomial distribution across all colonies, as previously described^23^. The beta-binomial distribution assesses the variance in mutant read support at all colonies for a given mutation. True somatic variants are expected to be present at high VAF (∼0.50) in some colonies and absent in others, yielding a high beta-binomial overdispersion parameter (⍴). In contrast, artefactual calls are likely to be present at low VAF across many colonies, which corresponds to low overdispersion. The maximum likelihood estimate of the overdispersion parameter ⍴ was calculated for each loci. For samples with greater than 25 colonies, SNVs with ⍴ < 0.1 and indels with ⍴ < 0.15 were discarded. For samples with fewer than 25 colonies, SNVs and indels with ⍴ < 0.20 were discarded. iv) *VAF filters.* Variants with VAF significantly lower than the expected VAF for clonal samples across all mutant genotyped colonies, as assessed with a binomial test with p threshold <0.001, were discarded. Additionally, variants with VAF less than half the median VAF of variants that pass the beta-binomial filter were discarded. v) *Germline filter.* All sites at which the aggregate VAF is not significantly less than 0.45 are assumed to be germline and discarded. The aggregate VAF is derived from the mutant read count across all colonies for a sample. The binomial test with a confidence threshold <0.001 was used to assess departure from germline VAF. vi) *Indel proximity filter.* SNVs were discarded if they occurred within 10 base pairs (bp) of a neighbouring indel. vii) *Missing site filter.* Loci at which genotype information is unavailable due to poor sequencing coverage will interfere with accurate phylogeny construction. Variants which have no genotype or coverage less than 6X in over one-third of samples were discarded. viii) *Clustered site filter.* SNVs and indels within 10 bp of a neighbouring SNV or indel, respectively, were filtered. ix) *Non-variable site filter.* Sites genotyped as mutant or wildtype in all colonies do not inform phylogeny relationships and are likely recurrent artefacts or germline variants, thus were discarded.

Some colonies were excluded based on low coverage or evidence of non-clonality or contamination. Visual inspection of filtered variant VAF distributions per colony was used to identify colonies with mean variant allele fraction (VAF)<0.4 or with evidence of non-clonality (Extended Data Fig.1C).

### Mutation burden estimation

Total SNV burden from WGS of individual colonies was corrected for differing depths of sequencing using a per-sample asymptomatic regression fit^80^ (Extended Data Fig.1E). A linear mixed effect model was used to estimate the rate of mutation acquisition with age, taking into account individual animals as a random effect as follows: *burden∼age + (0 + age|SampleID)*.

We filtered nanorate sequencing calls as previously described^12^, with the following modifications: we excluded variants (i) mapped to the mitochondrial genome, (ii) located within 15 bp of sequencing read ends, or (iii) observed in all duplex consensus reads as these are likely germline events. Matched colony whole genome sequencing data was used as a normal control. Mutation burdens were normalised to diploid genome size to determine the global SNV burdens.

### Phylogeny construction and quality control

Phylogenetic trees were constructed based on shared mutations between colonies for each mouse, as extensively described previously^9,34^. The steps, in brief, were as follows: (i) *Create genotype matrix.* Every colony has high sequencing coverage (median 14X) distributed evenly across the genome, allowing the determination of a genotype for nearly every mutated site observed across colonies. Each locus was annotated as Present, Absent, or Unknown in a read depth-specific manner. The number of unattributable sites was low, allowing precise inferences of colony interrelatedness. (ii) *Infer phylogenetic tree from genotype matrix.* We applied the maximum parsimony algorithm MPBoot to construct phylogenetic trees from the genotype matrix. Only SNVs were used to infer tree topology, but both SNVs and indels (if any) were assigned to inferred branches using *treemut*. Loci with unknown genotypes in at least one-third of colonies were annotated as missing sites and not used in phylogeny inference. (iii) *Normalise branch lengths for differing sequencing depth and sensitivity.* Branch lengths at this stage are defined by the number of mutations supporting each branch (molecular time). However, each colony has slightly different sequencing coverage, which correlates with differences in mutation detection sensitivity. Thus, we normalised branch lengths based on genome coverage to correct for sensitivity differences across colonies with varying depth, as described in ref. ^34^ (Extended Data Fig.1E). (iv) *Annotate trees with phenotype and genotype information.* Each terminal branch (tip) of a tree represents a specific colony. Thus, we annotated each branch of the tree with the sampled cell phenotype (HSC versus MPP).

Tree-level checks were used to identify any discordant branch assignments and assess the validity of tree topology. Any branches supported by variants with mean VAF <0.4 likely contained contamination by non-clonal variants and suggested the filtering strategy (see above) was insufficient. Similarly, the branch-level VAF distributions of every colony (tip) in the tree were manually inspected to confirm supporting variants were not present in unrelated portions of the tree (topology discordance). Finally, the trinucleotide spectra of individual somatic mutations were compared between those mutations located on shared branches (that is, mutations supported by >2 colonies) and mutations only observed once, and thus present on terminal branches. Mutation spectra were highly similar, indicating that mutations not shared by more than one colony were not populated by a relative excess of artefacts (Extended Data Fig.1F).

### Population size trajectories

We use the *phylodyn* package, which uses the density of coalescent events (bifurcations) in a phylogenetic tree to estimate the trajectory of *N*(*t*)/λ(*t*) over time^9,10^. Ultrametric lifespan-scaled trees were used to infer chronological timing. Under a neutral model of population dynamics, the phylogeny of a sample is a realisation of the coalescent process. In the coalescent process, the rate of coalescent events at time *t* is proportional to the ratio of population size, N(*t*), to the birth rate, λ(*t*) (which in the context of stem cell dynamics is the symmetric cell division rate). The sequence of inter-coalescent intervals across any time interval [*t*_1_, *t*_2_] is informative about the value of the parameter ratio *N*(*t*)/λ(*t*) across the same time interval. We note that only with a constant cell division rate λ over time can the trajectory parameter be interpreted as a scalar multiple of the trajectory of population size *N*(*t*). *Phylodyn* assumes isochronous sampling and a neutrally evolving population. We overlaid separate population size trajectories for HSCs and MPPs in Figure 3A.

### Approximate Bayesian computation

We used inference from phylodynamic trajectories to inform the development of an HSC population dynamics model. Population size trajectories from *phylodyn* indicated two successive ‘epochs’ of exponential growth, with some variation in growth rate between epochs, and a steady increase in population size over time (Fig.3A). Given the constraint of tissue volumes, it may be implausible that the HSC population grows constantly. We reconcile this discrepancy by noting that there are very few late-in-life coalescences in our phylogenies, and, as a consequence, the estimated *phylodyn* trajectory in late adulthood is associated with very wide credible intervals. We employed a population growth model based on a linear birth-death process^83^ (in which a population tends to grow exponentially, subject to stochastic fluctuations), together with a fixed upper bound N on population size. The model assumed a constant birth rate λ and constant death rate *v*, with the population trajectory growing at a rate λ-*v* within an epoch. The shape of the trajectory of N(*t*)/λ(*t*) depends on the cell division rate parameter λ, not only through the denominator in the ratio N(*t*)/λ(*t*), but also on λ-*v*, through the tendency of the population size N(*t*) to grow exponentially at a rate λ-*v*. In particular, if we increase the fixed upper limit N, and at the same time increase the cell division rate λ, so that their ratio remains constant, the shape of the trajectory of N(*t*)/λ(*t*) will change as a consequence of the changes in the value of the parameter λ. This suggests that the parameters λ, *v*, (in each epoch), and N, are all identifiable, and so can be estimated separately. The identifiability of λ, *v*, and N are expanded upon in Supplementary Note 4.

We applied Bayesian inference procedures^41^ to estimate the parameters (λ, *v*, and N) of the bounded birth-death process. We used Approximate Bayesian Computation (ABC). This method first generates simulations of population trajectories and (sample) phylogenetic trees across a lifespan. Each population simulation is run with specific values for the population dynamic parameters drawn from a prior distribution over biologically plausible ranges of parameter values. The ABC method includes a rejection step that retains only those parameter values which generated simulated phylogenies resembling the observed phylogeny (as measured by an appropriate Euclidean distance). The accepted simulations constitute a sample from the (approximate) posterior distribution. Population trajectories and sample phylogenies were simulated using the *rsimpop* R package. Approximate posterior distributions were computed using the R package, *abc.* We specified uniform joint prior densities for λ, *v*, and N which encompassed published estimates for N (population size) and λ (symmetric division rate)^31,73–75,84^: N ranged from 10^2^ to 10^5^ cells, λ ranged from 0.01 to 0.15 cell division per day, and *v* ranged from 0 to λ, such that the growth rate (λ-*v*) is always positive (as observed in the *phylodyn* trajectories).

Our population dynamics model was a birth-death process incorporating two separate growth epochs. The first (early) epoch lasted until 10 weeks post-conception, and the second (later) epoch lasted from 10 weeks onwards and corresponded to murine adulthood. Inferences were weak for the early epoch; thus, the later epoch was used for parameter inferences. Posterior densities from the three older mice were computed using the ‘rejection’ method (Extended Data Fig.6) and pooled to yield parameter estimates and credible intervals.

### Early life polytomy analysis

The polytomies were used to estimate lower and upper bounds for the mutation rate per symmetric division during embryogenesis. The method detailed in Lee-Six *et al.* ^10^ was used, whereby the number of edges with zero mutation counts at the top of the tree (up to the first 12 mutations) is inferred from the number and degree of polytomies assuming an underlying tree with binary bifurcations. The mutations per division are assumed to be Poisson distributed. A maximum likelihood range is then calculated in two steps: first, using the 95% confidence interval of the proportion of zero length edges, with this next leading to a maximum likelihood estimate for the Poisson rate. Sample M7183 lacked sufficient early life diversity (<10 unique lineages within 12 mutations molecular time) and thus was excluded.

### Shared variants between blood and colonic crypts

Mutation genotype matrices (described above) were generated for colonic crypt samples at loci observed in truncal (shared) branches in the matched HSC tree. Every variant was annotated as present or absent for each colonic crypt. We applied two stages to crypt annotation. First, a crypt sample was marked positive if the given variant exceeded a per-sample minimum VAF threshold. The minimum VAF threshold was defined as half the median VAF for all pass-filter colonic crypt variants (as described above). Next, for each variant represented in at least one crypt, any remaining crypt with >2 mutant allele read support was marked positive. This tiered definition allowed for shared variant capture despite differences in coverage among crypt samples. The proportion of a shared variant present among crypt samples was illustrated as a pie chart and annotated to the respective branch of the matched HSC tree (Extended Data Fig.3).

### Mutational signature analysis

We used the Hierarchical Dirichlet Process (HDP) algorithm to extract mutation signatures across aged and young HSC and MPP colony samples, following the process detailed in ref ^85^. Prior work in humans has applied mutation signature extraction to SNVs found only on terminal branches of phylogenetic trees – such terminal branches displayed mutation burdens in excess of 1000 mutations, depending on the organ. Given the low mutation burden in mouse hematopoietic colonies (terminal branch lengths spanning 30-150 mutations), and thus reduced mutational information, we utilised all branches with length ≥30 mutations as input. To circumvent any bias against shared variants, branches with less than 30 SNVs were collapsed to a single ‘shared branch’ sample. We generated mutation count matrices for each branch, using the 96 possible trinucleotide mutational contexts as input to the R package *hdp*. HDP was run i) without priors (de novo), ii) with the reference catalogue of all 79 signatures derived from the PanCancer Analysis of Whole Genomes study (COSMIC version 3.3.1) as priors, or iii) with the signatures previously defined as active in mouse colon^23^, SBS1, SBS5, SBS18, as priors. Trinucleotide signature definitions were adjusted to mouse genome mutation opportunities before usage as priors, and all prior signatures were weighted equally. Signature extraction parameters i) and ii) produced profiles that did not resemble any existing signatures (cosine similarity < 0.9), likely due to relatively limited SNV burden in mouse colony data. Usage of mouse colon signatures as prior information (iii) yielded four signature components. Two signature components demonstrated high similarity to SBS1 and SBS5 (cosine >0.9). The remaining two unknown components were deconvoluted to reattribute their composition to known signatures using the *fit_signatures* function from *sigfit*. This yielded three components with a reconstruction cosine similarity metric exceeding 0.99 for similarity to SBS1, SBS5, and SBS18, indicating these three signatures explain the majority of our data (Extended Data Fig.4A). We surmise the final reattribution step was necessary because of the log-fold lower SNV burdens in mouse blood colonies (30-200 mutations) relative to other tissues examined in previous work (>1000 mutations).

### Branch signatures assignment and analyses

For each mouse, we pooled the assigned SNVs into a “private” or “shared” category depending on whether the variant maps to a shared branch or not. Signature attribution to signatures SBS1, SBS5, and SBS18 was then carried out for each of these per mouse category using *sigfit*::*fit_to_signature* with the default “multinomial” model. The per-branch attributions were then carried out by 1) assigning a per-mutation signature membership probability and then 2) summing these signature membership probabilities over all SNVs assigned to a branch to obtain a branch - level signature attribution proportion. The per-mutation signature probability was calculated using:

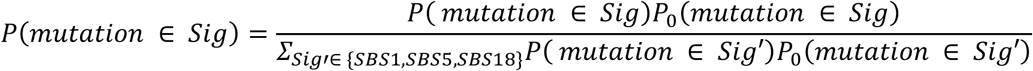

Where the prior probability,*P*_0_(*mutation* ∈ *Sig*), is given by the mean Sigfit attribution probability of the specified signature, *Sig*, for the category that the mutation belongs to.

A linear mixed effect model was used to assess the relationship between age and the signature - specific substitution burden for each colony while accounting for repeated measures. The signature-specific burdens per colony were estimated using a linear mixed model (R package *lme4*) with age as a random effect and mouse ID as grouping variable:

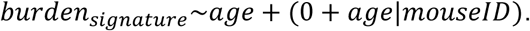

### Hidden Markov tree approach

#### Modelling the ancestral unobserved MPP and HSC states with a hidden Markov tree

We defined three unobservable (“hidden”) ancestral states, embryonic precursor cell (EMB), HSC and MPP, and used the observed outcomes (HSC or MPP tip states) to infer the transition probabilities between these identities and the most likely sequence of cell identity transitions during life. The transitions between states are modelled by a discrete time Markov chain with one step in time representing one mutation in molecular time. We require the root of the tree, presumably the zygote, to start in the “EMB” state and to stay in that state until 10 mutations in molecular time. After 10 mutations the cell then has a non-zero probability of transitioning to another state given by the transition transition matrix *M*:

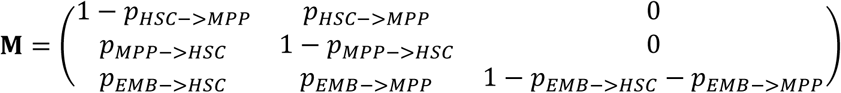

This then implies the following transition probabilities for branch *u*, having length *l*(*u*) (excluding any overlap with molecular time less than 10 mutations), starting in state *i* and ending in state *k*:

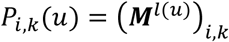

Now for a node that is in a specified state, the probability of descendent states is independent of the rest of the tree. This conditional independence property facilitates recursive calculation of a best path (“Viterbi path”), the likelihood of the Viterbi path, and the full likelihood of the observed phenotypes given the model. The approach is essentially an inhomogeneous special case of the approach previously described^86^.

Upward algorithm for determining likelihood of the observed states given *M* and a prior probability of root state π: The probability of the observed data descendant from a node *u* whose end of branch state is *i* is given by:

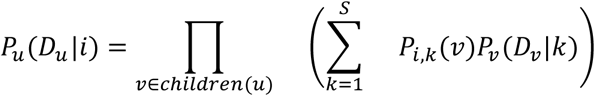

where *S* is the number of hidden states (*S* = 3 in our usage), and *D*_*u*_ denotes the observed data descendant of *u*, that is, the observed tip phenotypes of the clade defined by *u*.

#### Initialisation of terminal branches

The probability of observing a matching phenotype is assumed to be:

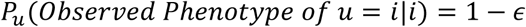

The probability of observing a mismatching phenotype, *j* ≠ *i*, is:

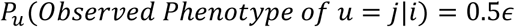

The root probability *P*_*roott*_(*D*_*roott*_|*i*) is calculated recursively from the above and the model likelihood is given by:

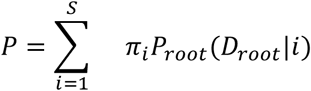

Given the two-stage cell sorting approach described above, we assume nearly error-free phenotyping and set є = 10^-12^.

#### Determining the most likely sequence of hidden end-of-branch states

This Viterbi-like algorithm can be run in conjunction with the upward algorithm. Here, instead of summing over all possible states, we keep track of the most likely descendant states for each possible state of the current node *u*.

The quantity δ_*u*_(*i*) is the probability of the most likely sequence of descendant states given that node *u* ends in state *i*:

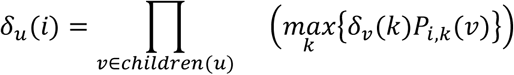

Additionally, for each node we store the most probable child states given that *u* is in state *i*:

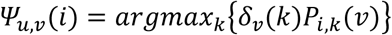

The tip deltas are initialised using the emission probabilities:

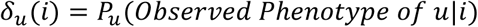

The above provides a recipe for recursively finding δ_*roott*_(*i*) and is combined with prior root probability π to give the most likely root state, *max*_*k*_{δ_*roott*_(*i*)}, in our case we set the prior probability of “EMB” to unity - so EMB is the starting state. The child node states are then directly populated using ψ.

### Targeted duplex-consensus sequencing

Genomic DNA from freshly collected peripheral blood was purified using the Zymo Quick-DNA Miniprep Plus kit according to the manufacturer’s instructions. 1650 ng of high-molecular-weight DNA was ultrasonically sheared to an average 300 bp fragment size using a Covaris M220 and ligated to duplex identifier sequencing adapters^87^ using the Twinstrand Biosciences DuplexSeq library prep kit. A large input of gDNA was used to ensure that the number of input genomic equivalents (about 275,000-330,000 genomes) did not limit the achievable duplex sensitivity. A custom baitset of biotinylated probes was used to enrich sequences targeting mouse orthologues of common human CH driver genes over two overnight hybridisation reactions. Our target panel spanned 61.8 kb and captured homologous regions from the entire coding region of the following genes: *Dnmt3a*, *Tet2*, *Asxl1*, *Trp53*, *Rad21*, *Cux1*, *Runx1*, Bcor, and Bcorl1, and specific exons with hotspot mutations (as observed in COSMIC) for the following genes: *Ppm1d*, *Sf3b1*, *Srsf2*, *U2af1*, *Zrsr2*, *Idh1*, *Idh2*, *Gnas*, *Gnb1*, *Cbl*, *Jak2*, *Ptpn11*, *Brcc3*, *Nras*, and *Kras*. Targeted loci encompass >95% of human CH events^43^ and are described in Supplementary File 2. Libraries were sequenced on the NovaSeq platform to a raw depth between 1-3 million reads, corresponding to duplex-consensus depths between 30,000-50,000X that vary across targeted exons (Supplementary Note 3). Quality control of duplex sequencing is discussed in Supplementary Note 3.

### Variant identification in targeted gene duplex-consensus sequencing

Duplex-consensus and single-strand consensus reads were generated using the *fgbio* suite of tools according the fgbio Best Practices FASTQ to Consensus Pipeline Guidelines (https://github.com/fulcrumgenomics/fgbio/blob/main/docs/best-practice-consensus-pipeline.md). To build a duplex-consensus read, we required at least 3 reads in each supporting read family (i.e., at least 3 sequenced PCR duplicates of matched top and bottom strands from an original DNA molecule). The ‘DuplexSeq Fastq to VCF’ (version 3.19.1) workflow hosted on DNANexus was also used to generate duplex-consensus reads. Next, VarDict^88^ was used to identify all putative variants, followed by functional annotation using Ensembl Variant Effect Predictor^89^. Finally, numerous post-processing filters were applied to remove false positives and artefactual variants: (i) *Quality flag filter.* VarDict annotates all variants using a series of quality flags that assess mapping and read-level fidelity^88^. Any variant with a quality flag other than “PASS” was discarded. (ii) *Read support filter.* Duplex sequencing enables detection of somatic variants even from a single read^87^; however, variants supported by a consensus read (singletons) were found to be highly enriched for spurious calls. Thus, any variant supported by a single read was discarded. (iii) *Mismatches per read filter.* Variants were excluded if the mean number of mismatches per supporting read exceeded 3.0. (iv) *End Repair & A-tailing artefact filter.* Library preparation enzymatic steps may introduce false positive SNVs near read ends due to misincorporation of adenine bases during A-tailing or mistemplating during blunting of fragmented 3’ ends. The fgbio FilterSomaticVcf tool was used to assess the probability that any variant within 20 bp of read ends was due to such enzymatic errors; probable end-repair artefacts were discarded. (v) *Read position filter.* Variants in positions ≤ 15 bp from the 5’ or 3’ end of a consensus read were observed to be enriched for spurious variants based on trinucleotide signature and were discarded. (vi) *Oxidative damage filter.* Mechanical fragmentation (prior to duplex adapter attachment) creates oxidative DNA damage, often in the form of 8-oxoguanine^90,91^, which mis-pairs with thymine and is fixed after PCR amplification. Any variant fitting the previously described oxidative artefact signature (SBS45) were discarded. (vii) *Sequencing coverage filter.* Variants at loci with duplex depth of ≤ 20,000X were considered under-sequenced and discarded. (viii) *Strand bias filter.* We employed a Fisher’s exact test to assess for forward or reverse strand bias between wildtype and mutant reads. Any variant enriched for unidirectional read support was discarded. (ix) *Recurrent variant filter.* Variants present in ≥5% of samples per duplex-sequencing batch or in ≥5 independent samples were discarded. (x) *Indel length filter.* Long insertions or deletions could be attributed to poor mapping, erroneous fragment ligation, or false positive calls by VarDict. Any indels ≥15 bp were excluded. (xi) *High VAF filter.* Germline variants display a VAF of 0.5 or 1.0. Any variants with VAF ≥0.4 were excluded as putative germline variants. (xii) *Impact filter.* CH is driven by functional coding sequence changes in driver genes. Thus, synonymous mutations were excluded during generation of the dot-plots in Figures 4-5. This filter was not utilised for analyses that require synonymous variant information (dN/dS, fitness effect estimation). (xiii) *Homologous position filter.* Residues conserved with humans are likely to be functional in mice. Variants at loci without a matching reference allele at homologous position in humans were discarded. This filter primarily eliminated intronic variants and was not utilised for analyses incorporating synonymous variant information. Variants identified are detailed in Supplementary File 2.

### Murine perturbation experiments

Perturbation experiments were initiated in aged (21-month) male and female mice unless otherwise described. Mice were randomly allocated to control or experimental groups. Investigators were not blinded to the group assignment during experiments. For *Mycobacterium avium* infection, mice were infected with 2 x 10^6^ colony-forming units of *M. avium* delivered intravenously as previously described^92^. Mice were infected once every 8 weeks (twice in total) to ensure chronic infection. For cisplatin exposure, mice were exposed to 3 mg/kg cisplatin delivered intraperitoneally every four weeks, as indicated. Dose spacing was selected to allow for sufficient recovery following myeloablation and blood counts were not altered in cisplatin-treated mice (Fig.4C), indicating recovery of haematopoiesis. For 5-Fluorouracil exposure, 150 mg/kg 5-FU was delivered intraperitoneally every four weeks two times; this 5-FU dose has previously been shown to drive temporary activation of HSCs in mice^93,94^. Exposure to a normalised microbial experience (NME) of murine transmissible pathogens was performed as previously described^51^. Briefly, immune-experienced “pet store” mice were purchased from pet stores around Minneapolis, MN. Aged (24-month) C57BL/6J:FVB/NJ laboratory mice were either directly cohoused with pet store mice or on soiled (fomite) bedding collected from cages of pet store mice. Mice were exposed to continuous fomite bedding for 1 month, followed by 5 months recovery on SPF bedding before tissue collection. All NME work was performed in the Dirty Mouse Colony Core Facility at the University of Minnesota, a BSL-3 facility. Age-matched C57BL/6J:FVB/NJ F1 laboratory mice maintained in specific pathogen free (SPF) conditions were used as controls. For monitoring, peripheral blood (∼50uL) was collected in EDTA-coated tubes and analysed on an OX-360 automated hemocytometer (Balio Diagnostics). For all aforementioned mouse cohorts, peripheral blood genomic DNA was purified and converted to duplex sequencing libraries as described above.

Differences in clone burden between control and treated cohorts was quantified using a Mann-Whitney test on cumulative VAFs per sample. Gene-level enrichment was measured using a Fisher’s exact test on the number and mutant and wildtype reads, normalised for coverage differences between samples. Gene-level dN/dS estimates were generated as described below.

### dN/dS analysis

The ratio of nonsynonymous to synonymous mutation rates (dN/dS) can be used to assess for selection within somatic mutations by comparing the observed dN/dS to that expected under neutral selection. We use the R package dNdScv^95^ to estimate dN/dS ratios of somatic mutations derived from whole genome and targeted gene duplex-consensus sequencing. To incorporate mouse-specific differences in trinucleotide context composition and background mutation rates, we generated a murine reference CDS dataset using the *buildref* function and genome annotations in Ensembl (version 102). For the phylogenetic trees, we input all tree variants to the *dndscv* function. dN/dS output and all coding variants detected in trees are listed in Supplementary File 1. To examine dN/dS in targeted duplex-consensus sequencing data, we pooled all variants observed in cross-sectionally sampled mice across ages (Fig.3A) and ran *dndscv* limited to exons only included on our targeted panel (Supplementary File 2).

### Targeted capture of tree variants

We designed a custom targeted DNA baitset (Agilent SureSelect) targeting mutations on the phylogenetic trees of the aged mice, and then queried genomic DNA purified from matched peripheral blood for tree-specific mutations using high-depth targeted sequencing. The baitset was designed to capture mutations on the phylogenetic trees of all 3 aged mice (MD7180, MD7181, and MD7182), and to cover mutations found in HSCs, MPPs and LSKs. The baitset was designed as follows: (i) All variants on shared branches that pass the SureDesign tool’s “moderately stringent filters”. (ii) All variants on a random subset of private branches that pass SureDesign’s “most stringent filters”. Approximately 25% of the private branches of each mouse were randomly selected. (iii) The exons and 3’ and 5’ UTRs for all CH driver genes used in our duplex sequencing panel (listed above). Target-enriched libraries were generated according to the manufacturer’s protocol and sequenced using the Illumina Novaseq platform. Baits were sequenced to median depths of 2616X, 2549X and 2628X for MD1780, MD7181 and MD7182 respectively.

To quantify the degree of HSC and MPP contribution to peripheral blood, we estimated the posterior distribution of true VAF for every mutation captured with our targeted baitset. This was done using the Gibbs sampling method previously developed^96^. Then, for each molecular time *t*, and for each branch that overlaps *t*, we estimate the VAF of a hypothetical mutation at time *t*. This is done by arranging our baitset variants in descending estimated VAF order at equally spaced intervals down the branch and then linearly interpolating the VAF at time *t* based on the estimated VAF of the neighbouring mutations. The aggregate VAF at time *t* for a tree or lineage is then calculated as the sum of the estimated VAFs of the overlapping branches at time *t*.

### Maximum likelihood estimates of fitness effects

#### Evolutionary framework

To generate estimates of fitness effects, mutation rates, and population size, we applied an evolutionary framework based on continuous time branching for HSCs, as previously reported^53^. The framework is based on a stochastic branching model of HSC dynamics, where variants with a variant-specific fitness effect, *s*, are acquired stochastically at a constant rate μ. Synonymous and nonsynonymous mutations detected with duplex sequencing in untreated 24-25-month-old mice were used in the analysis. Synonymous and nonsynonymous mutations were considered independently. Synonymous mutations are assumed to have no fitness effect and reflect behaviour under neutral drift, while non-synonymous mutations were hypothesised to reflect behaviour under a positive selective advantage. The density of variants declined at VAF 5•10^-5^, so to only include VAF ranges supported by informative variants, only variants above this threshold were included in maximum likelihood estimations described below.

How the distribution of VAFs, predicted by our evolutionary framework, changes with age (*t*), the variant’s fitness effect (*s*), the variant’s mutation rate (μ), the population size of HSCs (*N*) and the time in years between successive symmetric cell differentiation divisions (τ) is given by the following expression for the probability density as a function of *l* = *log*(*VAF*):

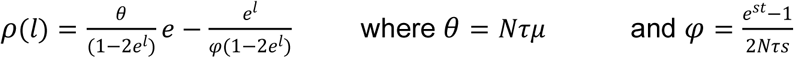

The value of 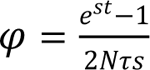 is the typical maximum VAF a variant can reach and this increases with fitness effect (*s*) and age (*t*). To reach VAFs > Φ requires a variant to both occur early in life and stochastically drift to high frequencies, which is unlikely. Therefore, the density of variants falls off exponentially for VAFs > Φ. For neutral mutations (*s* = 0),

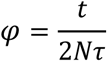

Because the mouse age *t* is known and the neutral Φ is measurable from the data, the ratio Φ/*t* allows us to infer *N*τ from the distribution of neutral mutation VAFs. Because the neutral θ is measurable from the data, and θ = *N*τμ, we can also infer the neutral mutation rate (μ).

Probability density histograms, as a function of log-transformed VAFs, were generated using Doane’s method for log(VAF) bin size calculation. Densities were normalised by the product of bin sample size and width. Estimates for *N*τ and μ were inferred using a maximum likelihood approach, minimising the L2 norm between the cumulative log densities and the predicted densities. For synonymous mutations, maximum likelihood estimates were optimised for *N*τ and μ. For nonsynonymous mutations, variants with VAFs below the observed maximum synonymous VAF (1.99•10^−4^) were used – these variants are within the “neutral” range – and estimates were optimised for with the *N*τ estimated from synonymous mutations.

#### Differential fitness effects

We estimated the distribution of fitness effects across nonsynonymous variants using our derived estimates of *N*τ and nonsynonymous μ. We parameterised the distribution of fitness effects using an exponential power distribution, which captures a strongly decreasing prevalence of mutations with high fitness:

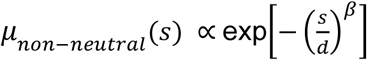

The shape of the distribution was fixed to β = 3^97^. Using the VAF density histograms from nonsynonymous variants, we estimated the scale of the distribution and non-neutral mutation rate: *∫^∞^_s=0_ μ_non-neutral_(s)ds*. The maximum likelihood fit predicted a scale of about *d*=2 and the proportion of non-neutral nonsynonymous mutations to be about 12% (Fig.6B).

### Code and data availability

SNVs and indels were detected using CaVEMan (version 1.14.0, https://github.com/cancerit/CaVEMan), cgpPindel (version 3.9.0, https://github.com/cancerit/cgpPindel), and VarDict (version 1.8.3, https://github.com/AstraZeneca-NGS/VarDictJava). Variants were annotated using VAGrENT (version 3.7.0, https://github.com/cancerit/VAGrENT), and Ensemble VEP (release 107-110.0, https://github.com/Ensembl/ensembl-vep). Phylogenies were constructed using MPBoot (version 1.1.0, https://github.com/diepthihoang/mpboot). Variants were assigned to phylogenies using Rtreemut (https://github.com/nangalialab/treemut). Population trajectories were inferred using *phylodyn* (https://github.com/mdkarcher/phylodyn). Bayesian inferences utilized the packages *rsimpop* (https://github.com/nangalialab/rsimpop) for simulations and *abc* (version 2.2.1, https://CRAN.R-project.org/package=abc) for approximate Baysesian Computation. Mutation signatures were inferred using the hdp (https://github.com/nicolaroberts/hdp) and sigfit (version 2.2.0, https://github.com/kgori/sigfit). Duplex consensus reads were generated using the fgbio suite of tools (version 1.5.1-2.1.0, http://fulcrumgenomics.github.io/fgbio/). dN/dS ratios were calculated using dNdScv (version 0.1.0, https://github.com/im3sanger/dndscv). Population genetic analyses of clone sizes and parameter inferences were based on code available at https://github.com/blundelllab/ClonalHematopoiesis/. Other analyses were carried out using custom R scripts and will be available at https://github.com/CDKapadia/somatic-mouse.

## Supplementary Note 1: Age equivalents between mouse and human

We used mouse and human survival data to estimate age equivalency between species. The median lifespan of C57BL/6J laboratory mice is 28-months^24^ (published data reproduced in Supplementary Fig.S1). We retrieved 2017 life-table data from the USA and the UK compiled at the Human Mortality Database (mortality.org). Only female data was included to match the makeup of our aged mouse dataset. We took the average of the median lifespans in the UK (82.5) and USA (80.7) to estimate the female mean lifespan as 81.6 years. Lastly, we normalised mouse age by median lifespan to determine an estimated equivalent human age. The above was only performed for the aged samples. Mice reach sexual maturity earlier in lifespan relative to humans, so age-equivalency was determined by onset of reproductive maturity between species, as previously described^98^.

**Supplementary Figure S1.**
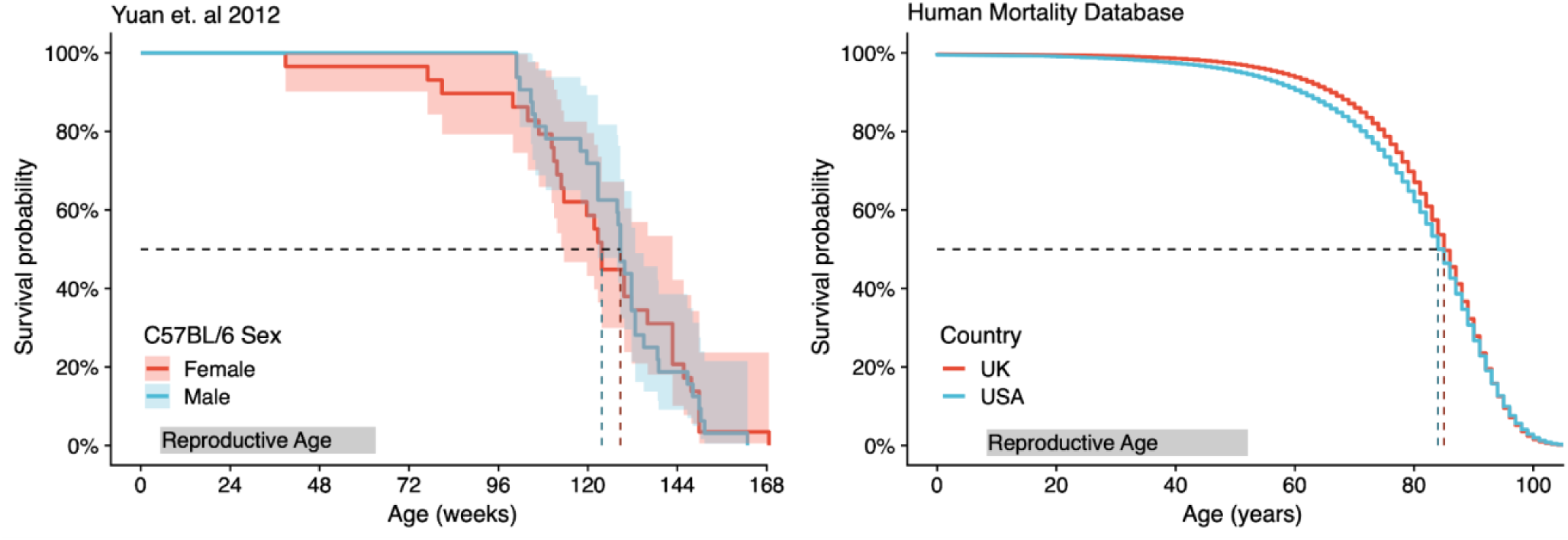
Mouse (C57BL/6J strain) survival data (left graph) by age for males (blue) and females (red). Human survival data (right graph) by age for females in the UK (red) and USA (blue). Dashed black lines mark the age of 50% survival probability for both species. Reproductive age is highlighted by the grey box.

## Supplementary Note 2: Ancestral cell identity inference

In our phylogenies, coalescences represent cell divisions of ancestral cells whose progeny have been captured as observable cells (tips on the tree). Comparison of the observed cell identity between closely related tips allows inferences of the identity of their most recent common ancestor (MRCA) and the nature of the ancestral cell division captured as a coalescence on the phylogenetic tree.

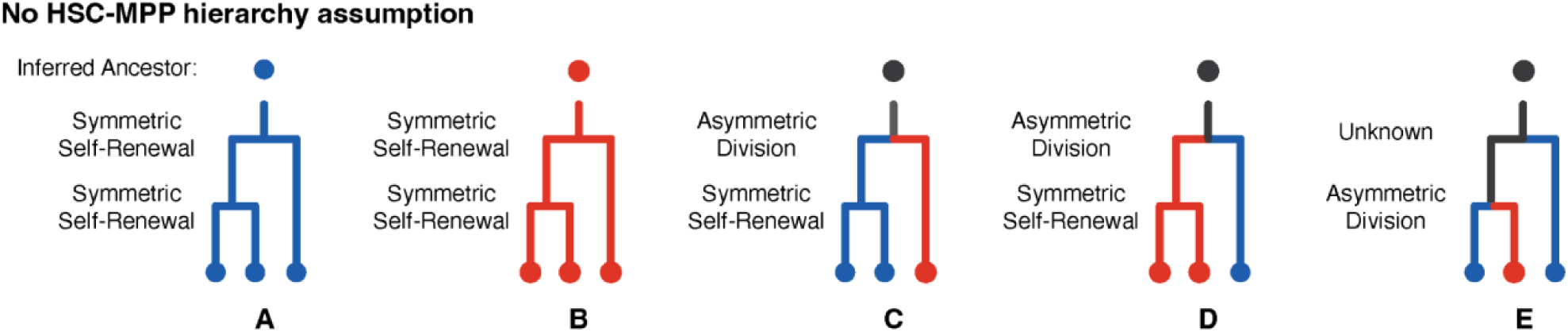

As an illustrative example above, if two closely related observed (‘tip’) cells are HSCs (scenarios A and C), then it is inferred that their most recent common ancestor was also an HSC. This HSC must have symmetrically divided to create two daughter HSCs, with both lines of descent also generating HSCs that were eventually sampled as the observed cells. From this inferred cell identity of their most recent common ancestor, if the cell state of the next closest relative is also an HSC, then their most recent common ancestor is similarly inferred to be an HSC (scenario A). HSCs coalescences are in blue, while MPPs are in red (scenarios B and D). Neighbouring tip states that differ in cell type (*e.g.,* 1 HSC and 1 MPP as in scenario E) can arise in two ways. First, there may have been an ancestral asymmetrical cell division generating one HSC and one MPP initially, with subsequent progeny along both lines of descent retaining these identities until sampling. Alternatively, the same tip states could also occur via a symmetrical self-renewing division of either MPP or HSC, followed by a later cell type change (*e.g.,* via asymmetric cell division or direct change) of one of the daughter cells. Either way, one cell type change from HSC to MPP (or MPP to HSC) is required to explain these tip states; therefore we mark their ancestral coalescence as blue/red. In these scenarios, because we cannot infer the cell identity of the MRCA, the upstream lineage is subsequently labelled in black. These principles can be applied to all coalescences in the observed phylogenetic trees (Fig.2A, Extended Data Fig.2, scenarios A-E above). This intuition does not rely on any assumptions of ontogeny, such as the hierarchy of HSCs over MPPs.

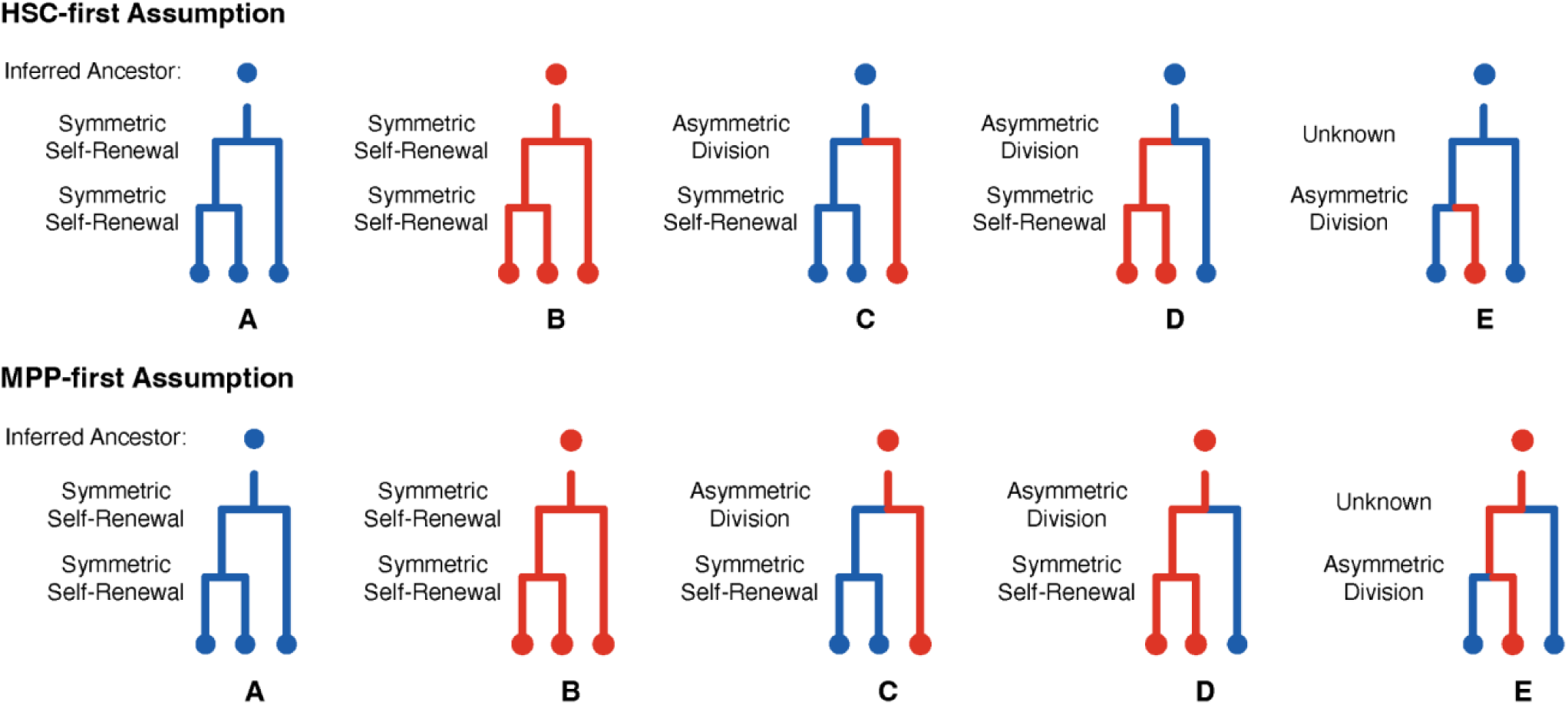

However, current models of the hematopoietic hierarchy dictate that HSCs give rise to MPPs (HSC>MPP). With this assumption made (top row “HSC-first assumption), one can label more of the branches and coalescences assigned as ‘black’ in the logic detailed above. For example, assuming an ‘HSC-first’ hierarchy, the common ancestor for scenarios C-E is now inferred to be an HSC, and the unobservable ancestral division in scenario E is inferred to be an HSC self-renewal. In the ‘MPP-first’ assumption (lower row), the inverse is inferred. These heuristics are applied to all coalescences in the observed phylogenetic trees.

We then asked how many cell state transitions are required to explain the tip states given an HSC-first or an MPP-first model. To perform this comparison, for each tree, we subsampled the largest category of HSC and MPPs so that there were equal numbers of MPP and HSC tips. To reduce the risk of the downsampling being unrepresentative the subsampling was conducted 10,000 times for each tree, and the average number of required transitions under the two unidirectional models was calculated. We then counted the total number of transitions required to result in the observed cell type tips. It was observed that the number of transitions required was similar for HSC-first and MPP-first and that there was no consistent pattern of one being higher than the other (Fig.2D). It was then natural to ask whether our cell type information was at all informative and so we randomly permuted the tip cell types and then resolved the tree in an HSC-first fashion. This sub-sampling and permutation was carried out 10,000 times and, as expected, the number of changes required by either the MPP-first or HSC-first models were generally far fewer than is consistent with the null model that all balanced cell type categorisations require the same number of tree-based transitions to explain the tip phenotype under an unidirectional model. In summary, both HSC-first and MPP-first models are less parsimonious, i.e., requiring more cell state changes, than the model first presented in which no assumptions are made about a hierarchy between HSCs and MPPs. The most parsimonious model would be that of HSC and MPP lineages being derived in parallel during similar development periods from non-overlapping common ancestors.

### A simple 3 state model for Murine Progenitor Ontogeny

To formalise the above ideas in the context of a simple model of HSC and MPP ontogeny, we considered the state of all cells prior to 10 mutations in molecular time as being in an embryonic precursor state (EMB), given that haematopoietic and colonic lineages remain uncommitted until at least this time (Extended Data Fig.3). We then assumed that in each unit molecular time there is a fixed probability of transitioning out of this embryonic state into either an HSC state, *p*_EMB → HSC_, or an MPP state, *p*_EMB→MPP_. Furthermore, there is a fixed probability of transitioning from an HSC to an MPP, *p*_HSC→MPP_, and from an MPP to an HSC, *p*_MPP→HSC_. Thus, the evolution of the cells down the tree is governed by a discrete time Markov chain process. The likelihood of the observed tip cell types is calculated using a hidden Markov tree approach (Methods). Maximum likelihood estimates of the model parameters are obtained by maximising the sum of the log-likelihoods across mouse-specific phylogenetic trees. Finally, for each mouse, the most likely sequence of unobserved states for the nodes of the phylogenetic tree is calculated using the fitted model parameters.

We performed the maximum likelihood estimation using the R package “bbmle”. The maximisation was performed on logit transformed quantities: 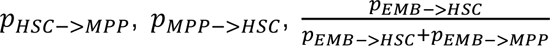 and intervals did not work in all cases. So, to obtain more robust estimates in the CIs of the model we implemented a Stan-based Bayesian version of the model using the directly calculated likelihood as described above. Uniform priors on the unit interval were assumed for 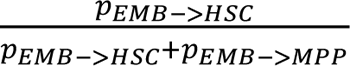 and *p_EMB→HSC_ + p_EMB→MPP_* uniform priors on the interval (0-0.5) were assumed for both *p*_HSC→MPP_ and *p*_MPP→HSC_. The model was run with four chains, each for 10,000 iterations.

### Separate young and old mouse cohorts provide optimal model fit

We compared fitting the model with a per-mouse, per-age, and pan cohort strata. A likelihood ratio test analysis revealed that the best model is an age-specific model where parameters are estimated separately in old and young mice (Supplementary Table S1).

**Supplementary Table S1.** Log likelihood values and Akaike information Criterion (AIC) assessing model fit.

| Model | Degrees of freedom | Log Likelihood | AIC | Likelihood Ratio Test |
| --- | --- | --- | --- | --- |
| Pan Cohort | 4 | -840.2 | 1,688.4 |  |
| Age-Specific | 8 | -800.2 | 1,616.5 | vs. Pan Cohort: P=1.78e-16 |
| Mouse-Specific | 24 | -796.2 | 1,640.3 | vs. Age Specific: P=0.945 |

### Young and Old mice exhibit differing patterns of differentiation

Applying our hidden Markov tree approach, we fitted an HSC-first model where,

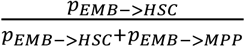

is fixed at unity. That is, all EMB must transition to an HSC before any emergence of MPPs (HSC-first). In the context of this simple model, we can reject the HSC-first model across the combined age group model (p=1.11e-18) and also for the old group (p=1.38e-19). However, we were unable to reject the HSC-first model for the young animal group (p=0.397).

Examining the types of cell-state transitions in the trees, we observed that aged animals exhibit several independent transitions from the embryonic precursor state followed by relatively few transitions between HSC to MPP or vice versa (Supplementary Fig.S2). In contrast, young animals exhibit a tendency towards HSC-first followed by a relative abundance of HSC->MPP transitions (Supplementary Fig.S2). Both HSC-specification and MPP-specification occur within the first 50 mutations molecular time (Supplementary Fig.S3). The cell identity transition rates, per unit molecular time, are listed below and were used to generate Fig.2E.

| | $p_{EMB \rightarrow HSC}$ | $p_{EMB \rightarrow MPP}$ | $p_{HSC \rightarrow MPP}$ | $p_{MPP \rightarrow HSC}$ |
| --- | --- | --- | --- | --- |
| Young Donors | 0.158 | 0.037 | 0.0164 | 0.0070 |
| Aged Donors | 0.036 | 0.034 | 0.0013 | 0.0006 |

The above result are fairly consistent with the Stan based results for which we show the medians of the marginal posterior distribution followed by the 95% credibility intervals:

| | $p_{EMB \rightarrow HSC}$ | $p_{EMB \rightarrow MPP}$ | $p_{HSC \rightarrow MPP}$ | $p_{MPP \rightarrow HSC}$ |
| --- | --- | --- | --- | --- |
| Young | 0.43(0.11 - 0.9) | 0.063(0.0048 - 0.27) | 0.017(0.014 - 0.022) | 0.0078(0.0029 - 0.016) |
| Aged | 0.04(0.025 - 0.068) | 0.038(0.025 - 0.064) | 0.0014(0.00079 - 0.0022) | 0.00071(0.00022 - 0.0015) |

Of note, the mode of the marginal posterior distribution of *p*_EMB→HSC_ peaks at 0.19, which is reassuringly close to the maximum likelihood estimate of 0.158.

**Supplemental Figure S2:**
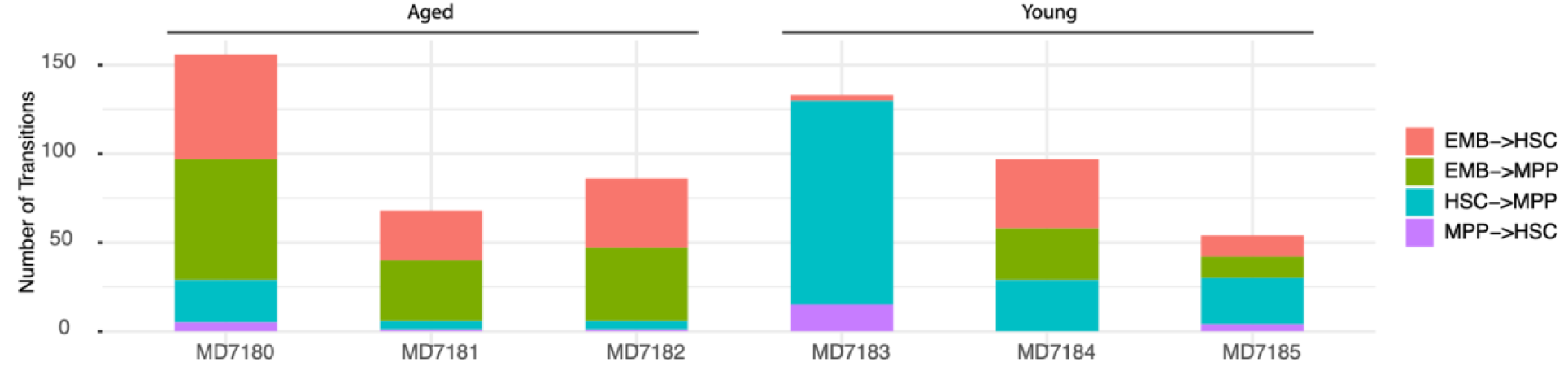
Transition Type Counts. The old mice exhibit an abundance of approximately equally prevalent EMB->HSC and EMB->MPP transitions followed by relatively few transitions to the eventual observed cell types. The young mice exhibit relatively fewer EMB->HSC and EMB->MPP and then a relative abundance of HSC->MPP transitions.

**Supplementary Figure S3:**
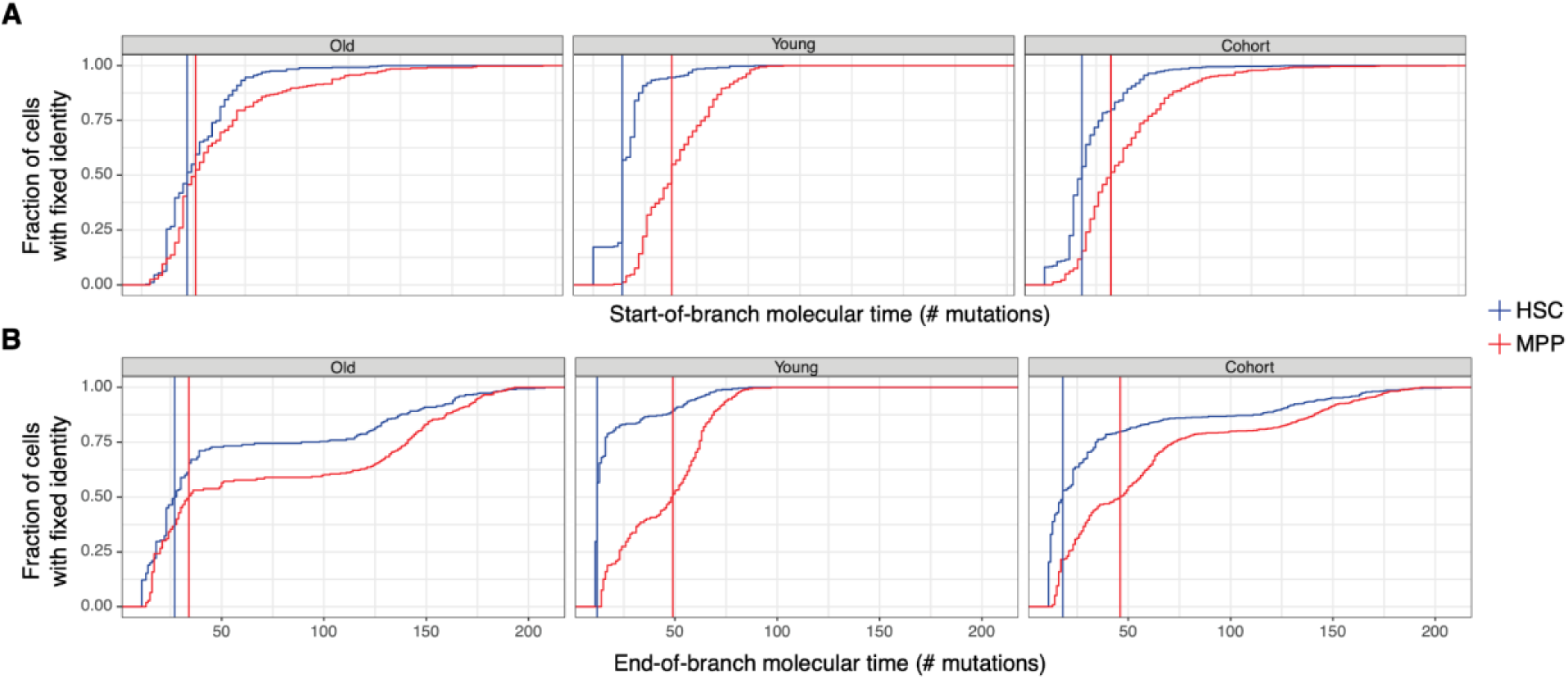
Cumulative Distribution of Specification Timing. For each colony we use the molecular time of the **A)** start of the branch, or **B)** end of the branch on which the ancestral lineage first transitions to the observed cell type as an upper bound for the timing of its transition to its final observed state. The panels show the cumulative distribution of these upper bounds calculated form the most likely sequence of transitions inferred using the age-specific model and the pan cohort model. Vertical lines indicate the time at which 50% of the sampled cells have specified identity.

## Supplementary Note 3: Quality control of targeted duplex-sequencing

We expected that somatic clones in mice might be rare events at small clone sizes, thus would require a sensitive detection assay. High-depth sequencing can be used for detection of subclonal variants, but with increasing coverage, the error-rate intrinsic to short-read sequencing can obscure true low variant allele fraction (VAF) variants. To circumvent this sensitivity limit, read-level error-correction approaches are necessary. Thus, we applied duplex-consensus sequencing, which offers among the highest sensitivity for subclonal variant detection. In duplex-sequencing, each initial dsDNA molecule is uniquely barcoded such that reads derived from complementary 5’ and 3’ strands are linked, but also distinguishable. Detected variants must be present on both uniquely barcoded strands of the initial dsDNA fragment to pass bioinformatic filtration (Supplementary Fig.S4). By enforcing that variants are present in reads derived from both of the matched complementary strands of DNA, one can eliminate the majority of sequencer-induced artefacts that usually hamper sensitivity. To apply this technology to murine clonal haematopoiesis (CH), we developed a target panel of the mouse homologs of genes most frequently mutated in human CH (Methods, Supplemental File 2).

**Supplementary Figure S4:**
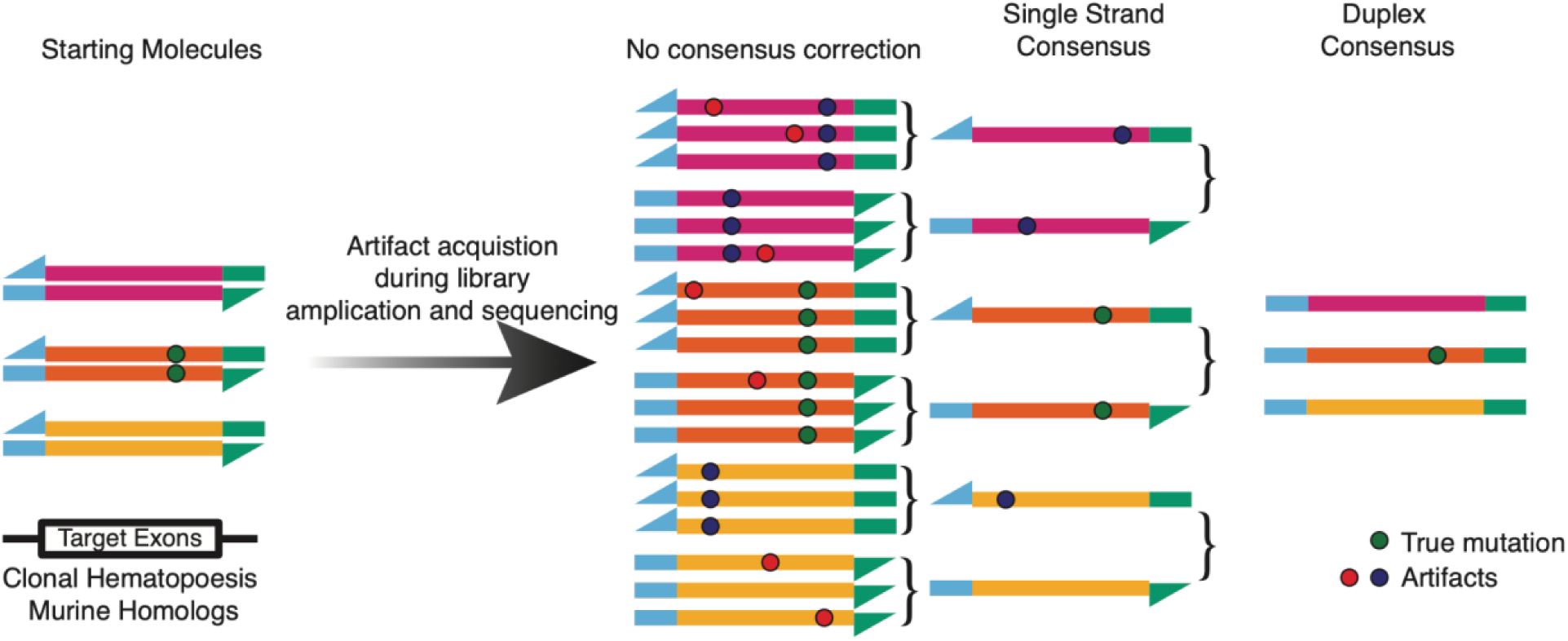
Error correction strategy in targeted duplex sequencing.

PCR during library preparation and sequencing introduce low-frequency artefacts. Duplex barcodes allow grouping of PCR duplex reads from a single DNA library molecule (read families) and single-strand read consensus generation. Next, single strand consensus reads from complementary strands on initial dsDNA are matched to generate a duplex consensus. To build a duplex-consensus read, we required at least 3 reads in each supporting read family (i.e., at least 3 sequenced PCR duplicates of matched top and bottom strands from an original dsDNA molecule).

### Coverage requirements to generate a duplex consensus

To generate a duplex-consensus read, an initial DNA molecule must be sequenced multiple times with reads from matched 5’ and 3’ strands sufficiently represented. To ensure that clone detection sensitivity would not be limited by input genomic DNA (*i.e.*, the libraries contained sufficient genomic complexity), we input at least 100,000 genomic equivalents (or at least 1650 ng of genomic DNA) into our library preparations. High library complexity decreases the probability of matched 5’ and 3’ reads being sequenced by chance; thus, even with a target panel enrichment, extremely high sequencing depth is required to capture library complexity in duplex consensus reads. Median raw, non-deduplicated coverage spanned 1,000,000X to 3,000,000X at targeted loci per sample. This correlated with a single-strand consensus coverage spanning 60,000X-120,000X, which, after 5’ and 3’ linkage, further collapsed to duplex consensus coverage spanning 30,000X-40,000X (Supplementary Fig.S5). Duplex coverage at specific exons within targeted genes was variable between samples (Supplementary Fig.S6)

**Supplementary Figure S5:**
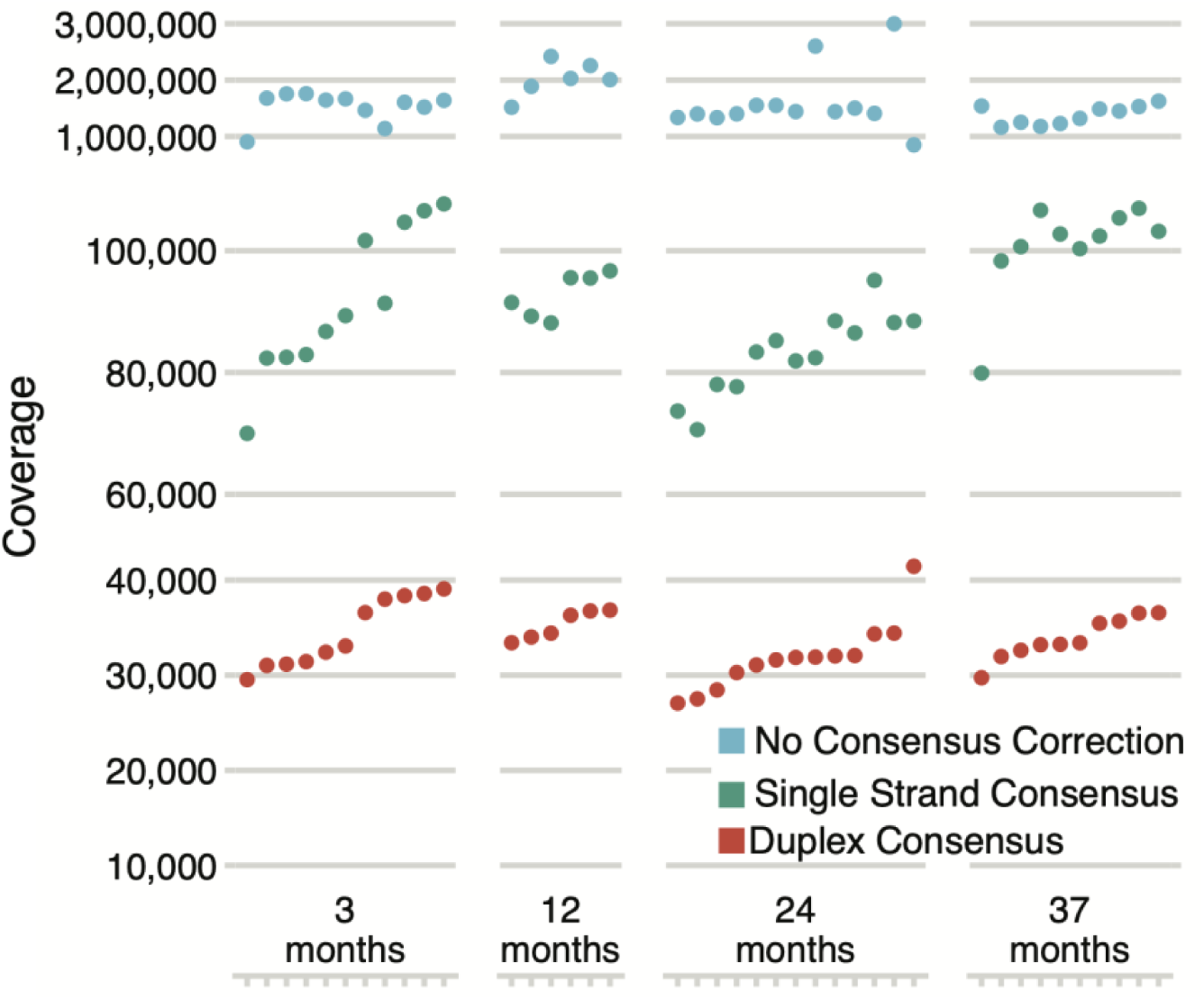
Sequencing coverage at targeted loci for all samples in Fig.4A. The relationship between raw (not deduplicated), single strand consensus, and duplex consensus coverage is shown.

**Supplementary Figure S6:**
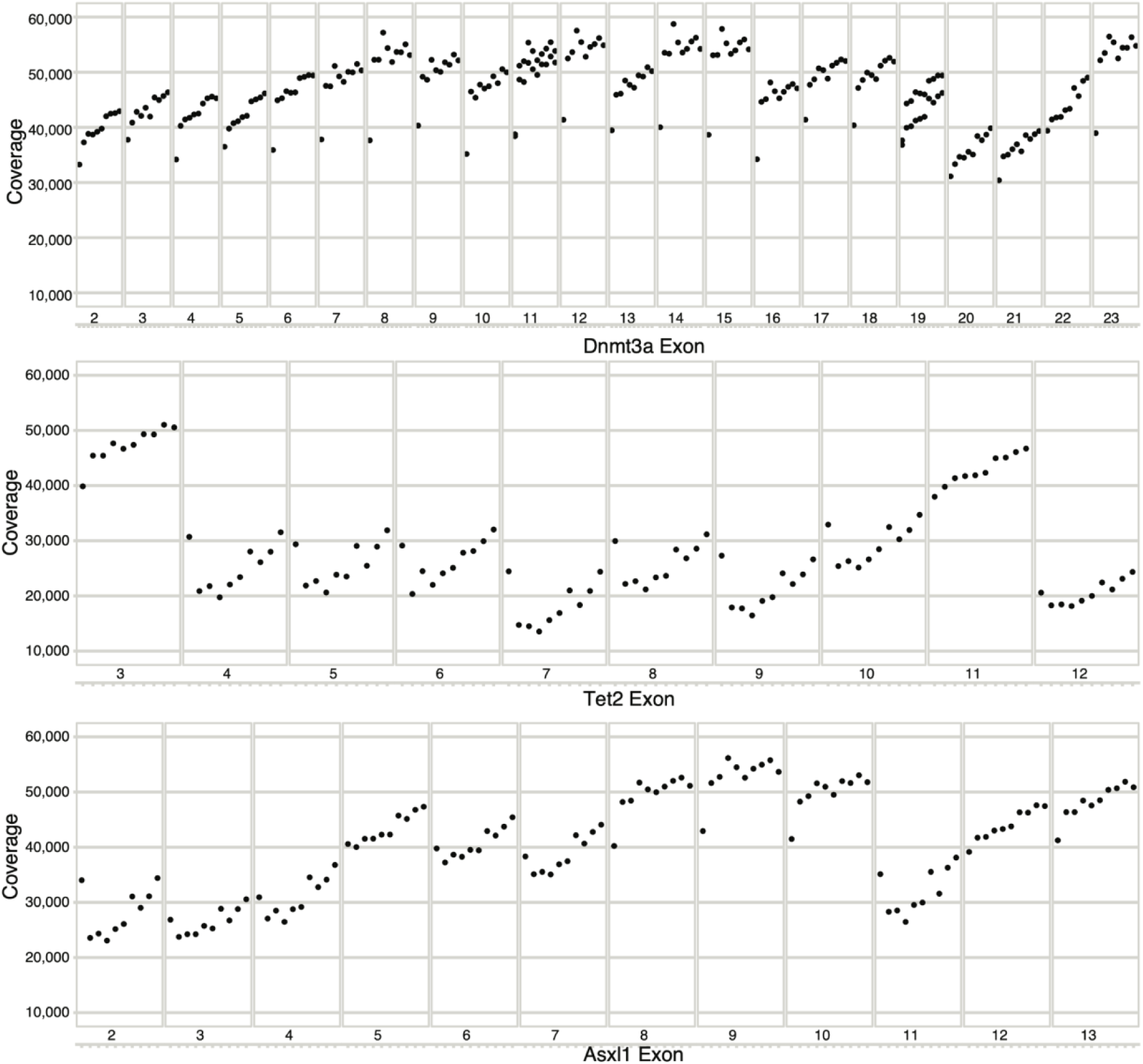
Duplex coverage at coding exons in *Dnmt3a*, *Tet2*, and *Asxl1* for a series of aged samples.

### Mutation filtering

To supplement the sensitivity afforded by duplex sequencing, stringent read- and variant-level filters were applied to reduce the presence of false positive mutations or spurious calls. Without filtration, we observed an enrichment of C>A mutations (Supplementary Fig.S7), reminiscent of mutation signature SBS45,which is likely attributable to oxidative damage during sequencing^90,91^. Such oxidative damage mutations likely arose after duplex barcode attachment, were enriched at read ends, and likely caused mutations within the duplex barcode sequence. Due to mutations in duplex barcodes, a read family derived from a single initial dsDNA molecule (a singleton) would erroneously appear as derived from an additional read family (a doublet). This observation led us to apply a stringent series of filters (Methods), after which the trinucleotide spectra of variants detected in duplex sequencing more resembled that seen with blood (Supplementary Fig.S7).

**Supplementary Figure S7:**
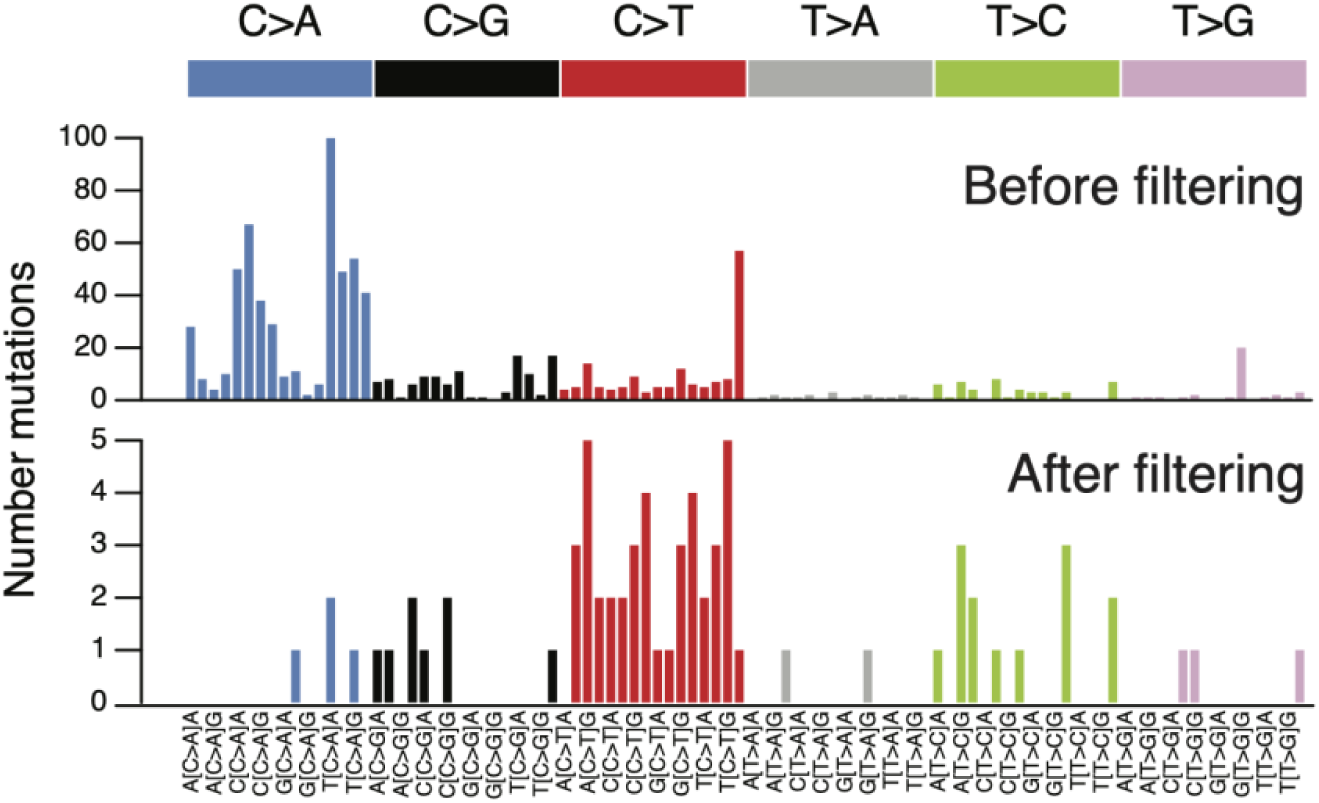
Trinucleotide spectra of duplex-sequencing variants before and after post-processing filters. See Methods for filtering strategy details.

### Clone size calculation

Given the differences in coverage between loci, we normalised the variant read counts to allow accurate clone size comparisons between samples. In general, clone size for a given variant is defined as:

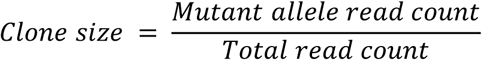

For very small clones, there is a degree of stochasticity affecting if sufficient mutant read alleles will be converted to duplex consensus reads to allow detection. Duplex clones supported by very few mutant allele reads would have a low numerator, thus clone size estimations may be skewed. Given more single-strand consensus reads are generated than duplex consensus reads (Supplementary Fig.S6), we reasoned that mutant allele reads would be relatively more abundant within single-strand consensus reads – that is, mutant allele reads would be present among the reads ‘discarded’ due to insufficient evidence to generate a duplex consensus. To normalise clone size, especially in low-magnitude clones, we used de-duplicated single-strand consensus reads, as follows:

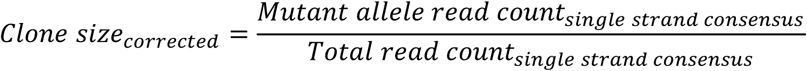

By using the de-duplicated single-strand consensus reads for clone size calculation, the numerator (variant allele count) and denominator (coverage) both increase, reducing any skewing that may be present in clone size calculations from duplex consensus reads. All clone sizes depicted on dot plots are calculated in this manner.

### Biological replicates

To validate reproducibility within the targeted duplex-sequencing library preparation and variant calling pipelines, we assessed clone prevalence in biological replicate samples. For each replicate, peripheral blood was separately collected (in different tubes) and underwent genomic DNA extraction independently. Thus, the genomic DNA “pools”, while derived from the same sample mouse, were purified in separate reactions. Replicate DNA samples underwent duplex library preparation and variant calling as described in Methods. As shown in Supplementary Figure S8, clone detection is concordant between paired replicates. Clones unique to a single replicate were at the limit of detection for the specific locus, and thus it is likely in the paired replicate that insufficient variant reads were sequenced to generate duplex consensus read support. Such borderline detectable clones will likely be detectable within single-strand consensus reads, which carry nearly double greater read depth, though at the expense of duplex sensitivity. We examined single-strand consensus reads from the biological duplicate samples and were able to “rescue” missing variants from the paired replicate sample, in about half of cases(Supplementary Figure S8). This confirms that much of the missing replicate clones were lost during duplex consensus building, for example when a clone has insufficient top or bottom strand support to create a duplex read.

**Supplementary Figure S8.**
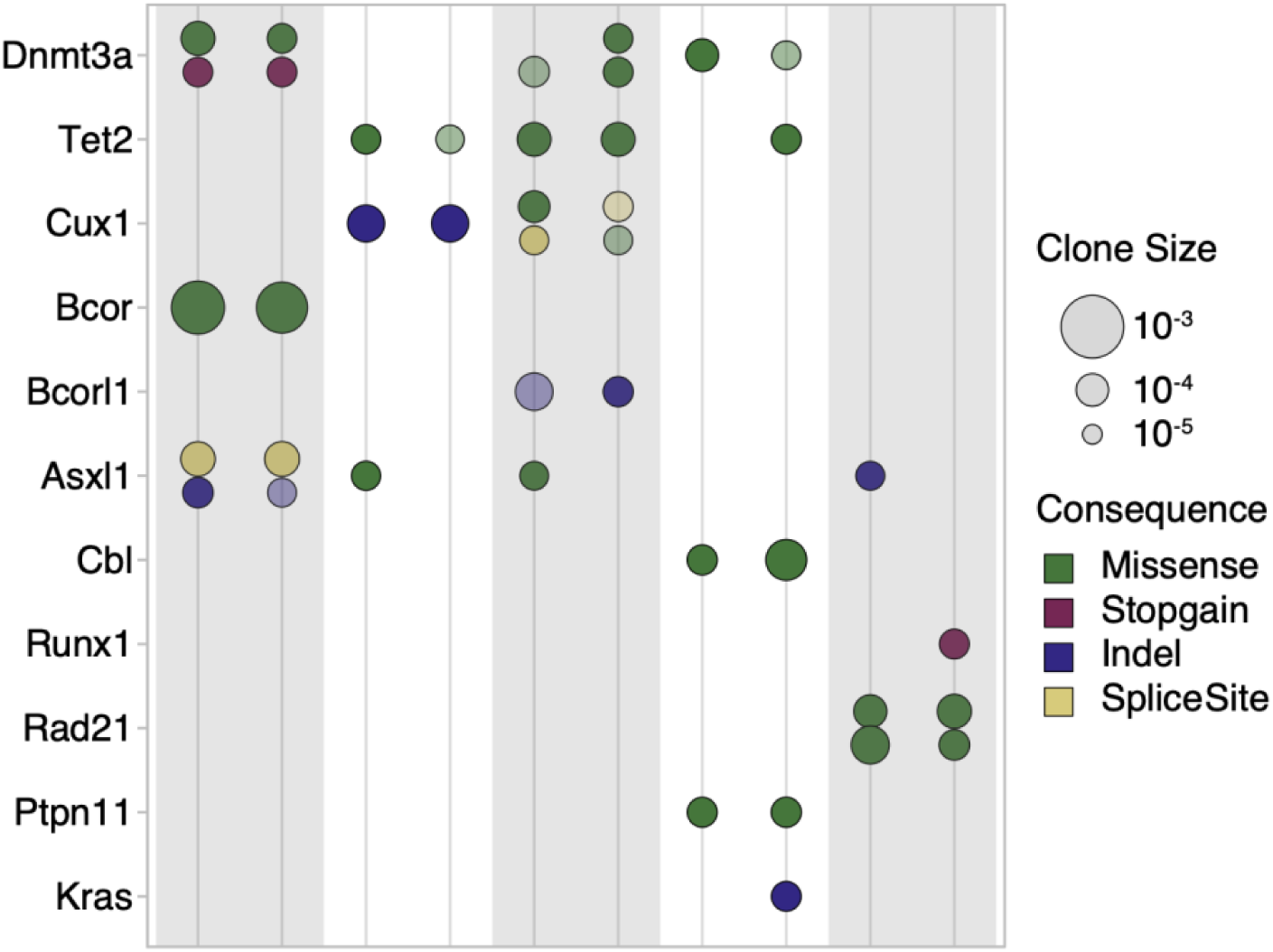
Native CH in biological replicate samples. Shaded and unshaded pairs represent duplex libraries separately prepared from an identical initial blood sample. Clones are presented as described in Fig.4A. Transparency indicates a clone that was only detectable within single strand consensus reads but not duplex consensus reads.

### In silico estimation of the sensitivity and specificity of duplex-sequencing results

We next sought to understand if the degree of clone concordance between biological replicate samples was consistent with the sensitivity of our assay. We consider a simple model for SNVs of conditional base calling probabilities for the reference base (R), a mutant base (A) and the two other bases (B,C). For an individual read (or read family/bundle) the probability of observing the bases is modeled in the following manner:

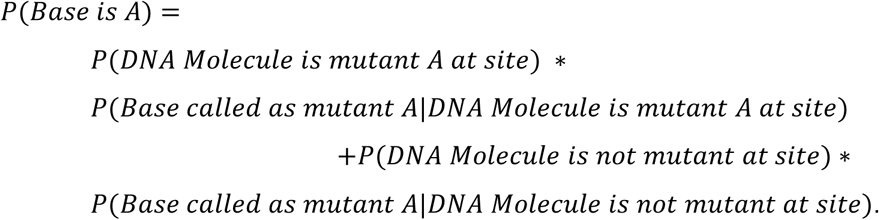

Now 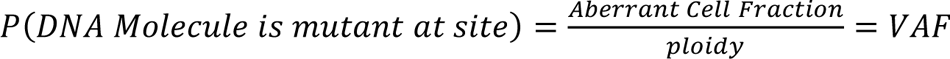 where for economy we now use the term (true) VAF to characterise the clone. Moreover we assume there is a base calling error rate (“epsilon”) є. It is assumed that this results in the one of the 3 incorrect bases to be called with equal probability of є/*3*:

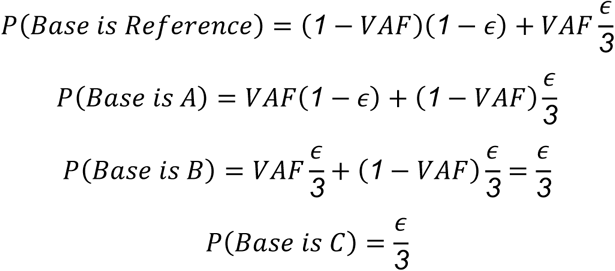

For a given bait set wide depth of sequencing, *depth*, a given site has depth that is Poisson distributed with mean *depth*. For a clone to be detected it is only required that at least 2 mutant reads are observed. We assume we have a known clone, with VAF=1^-4^ or VAF=1^-3^, and with mutant allele A. The A clone is discovered if there are 2 or more mutant “A” reads, and no other mutant reads (“B” or “C”). With these criteria, we can plot the sensitivity for a given error rate, є, shown below in Supplementary Figure S9.

**Supplementary Figure S9:**
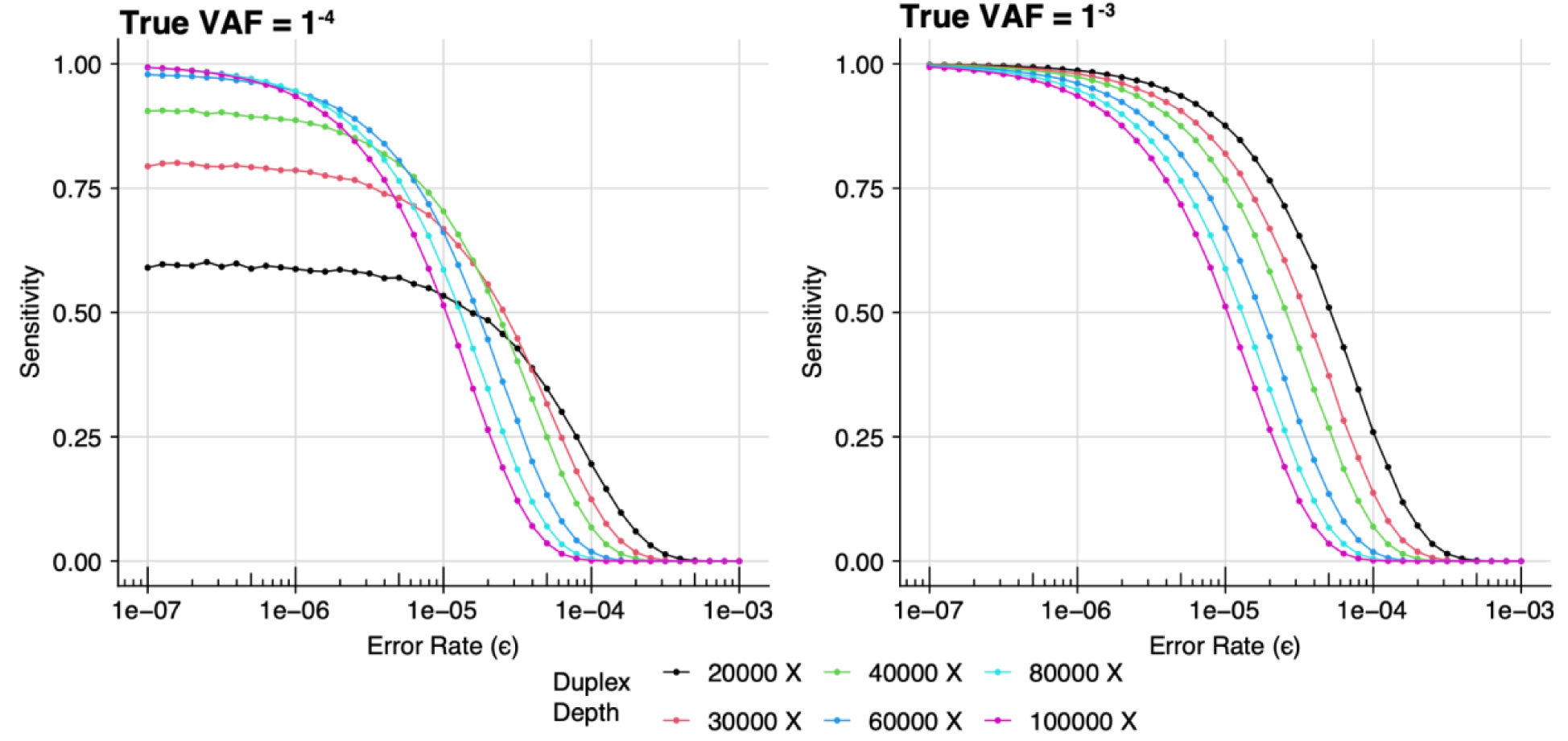
True clone discovery across error rates using multinomial modeling. Estimated sensitivity of detecting a variant at a given site with true VAF 1^-3^ **(left)** or 1^-4^ **(right)** across increasing error rates. A range of duplex depth at variant sites are shown.

The above plots show that using single strand consensus sequencing with error rate of ∼3 ^-5^ at depth 60,000x provides a sensitivity of 30% for clone sizes of VAF=1^-3^ or less. However, using duplex depth of 30,000x with an error rate of 1^-6^ to 1^-7^ (as described in Kennedy *et al*.^87^) provides a sensitivity of >75%.

If we assume the extreme (and implausible) case of error-free sequencing, then the clone detection sensitivity is purely governed by the binomial distribution with a probability of True VAF. Importantly, even if the sequencing was error-free, we would not expect there to be concordance of clone detectability in different samples. In the Supplementary Figure S10 below we can see that for error-free sequencing at depth 20,000X, we would actually have a concordance of around 60%. This aligns with the observed duplex clone concordance seen in the biological replicate samples shown in Supplementary Figure S8.

**Supplementary Figure S10:**
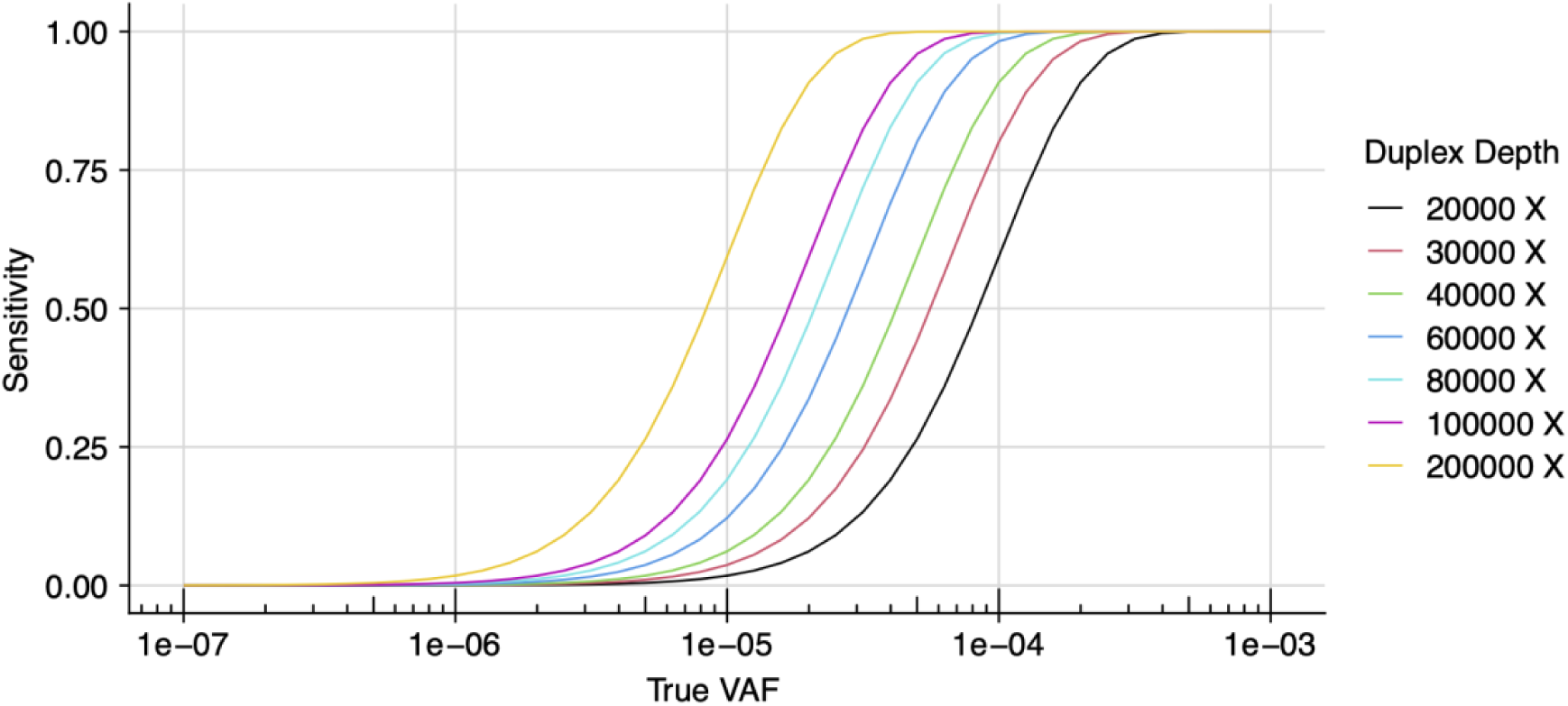
True clone discovery with error-free sequencing. The estimated sensitivity for clone detection at increasing VAFs in the scenario of error-free variant detection. In the absence of an error rate, detection sensitivity can be described with a binomial distribution. A range of duplex depth at variant sites are shown.

Finally, we can estimate the probability of false positive clone detection at a given error rate є. As shown below, when querying a range of feasible duplex-sequencing sensitivities and duplex-corrected sequencing depths, a false positive clone is far less likely than a false negative clone (missing a true event). As an illustrative example, for duplex depths 20,000X to 30,000X and the duplex error rate of <8e-04 (estimated error rate of <1e-06), the false positive rate is <0.01. (Supplementary Figure S11).

**Supplementary Figure S11:**
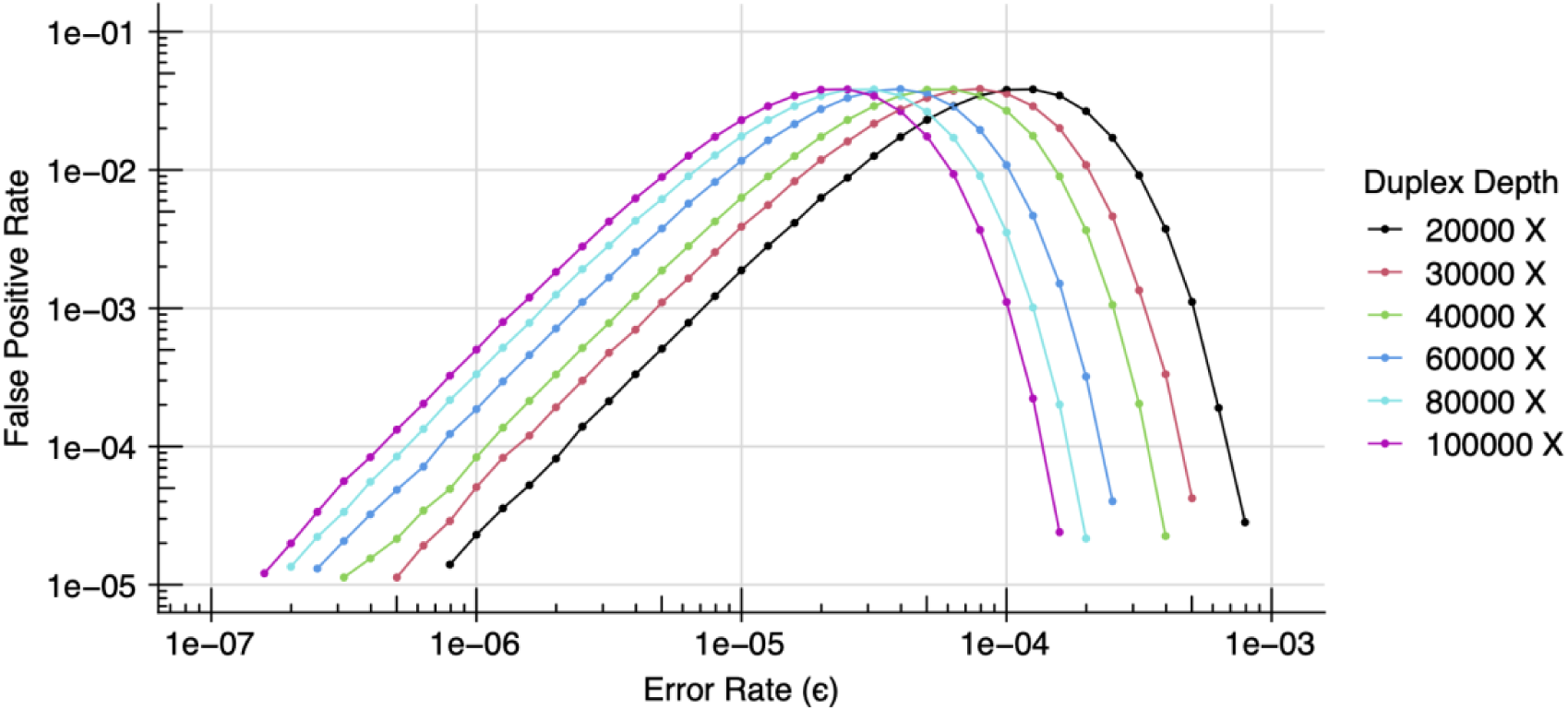
False positive variant detection. The estimated incidence of incorrectly detecting a variant at a given site is shown, using multinomial modeling of detection error rate and site-specific duplex depth.

### Concordance of duplex sequencing data explored through mixing mutant and wildtype reads

The in-silico analyses described above suggest that a true variant clone may not be observed due to insufficient duplex read support, and sensitivity increases with additional duplex depth. In this case, an *expected* variant would likely be detectable in single-strand consensus reads (Supplementary Figure S4), which require reduced read support to build a consensus read, and harbour far higher coverage (Supplementary Figure 5), though at the expense of sensitivity.

We performed a mixing analysis using our duplex data, with the aim to evaluate 1) the concordance of calling serially lower VAF clones in different sample, and 2) the degree missing-but-expected clones can be found in single-strand consensus data.

We selected clones with a large detectable clone size, then generated serial dilutions of input mutant file reads with wild-type file reads to simulate diminishing read support and the subsequent detection of an expected variant in duplex reads. Mutant file reads were diluted by the following percentages: 50%, 20%, 10%, 5%, 2%. Five replicates of each random subsample dilution were used as technical replicates. Read dilution was done with raw, unmodified reads; that is, before any mapping or consensus building steps. Mutant reads were mixed with wildtype reads to the same overall read count as the original data, then analysed using the duplex consensus building and variant calling pipeline described herein. In cases where the expected variant was not detected in duplex consensus reads (either due to lack of read support, or failing to pass stringent filters), we examined matched single strand consensus reads for the variant, and often were able to detect the expected clone.

**Supplementary Figure S12:**
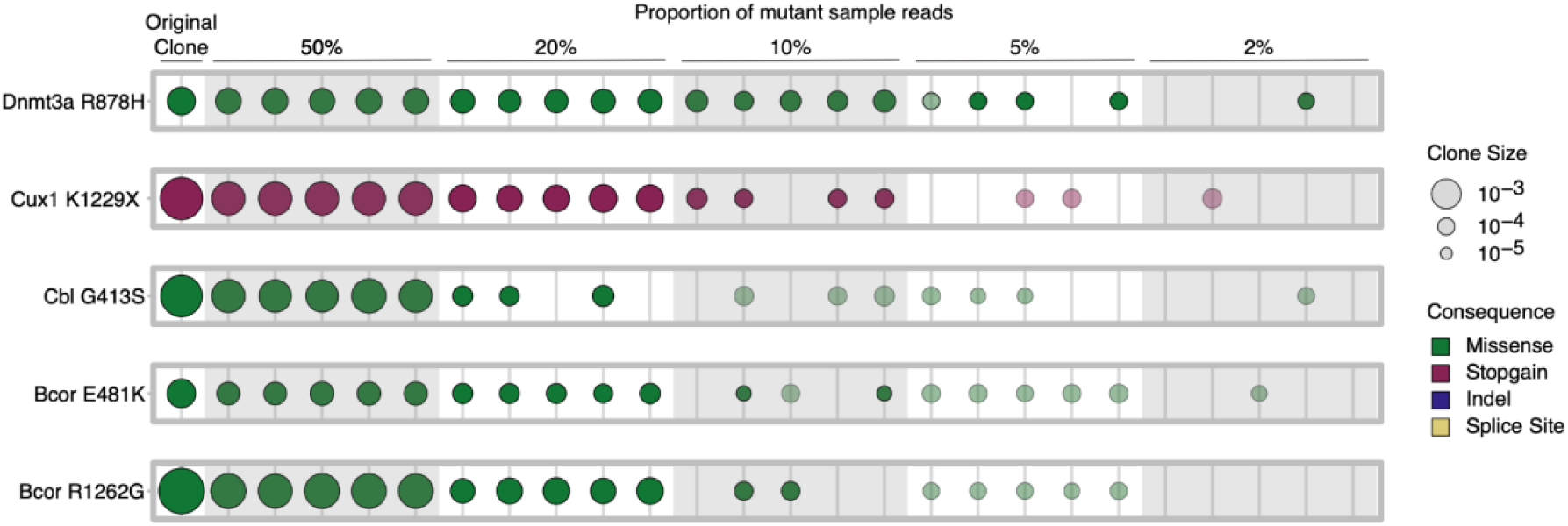
Dilution of mutant reads and subsequent variant call concordance. For five initially large clones, reads from the original input file (ie supporting the observed clone) were diluted at the indicated proportion with wildtype reads. Five replicates for each dilution factor are grouped. The original clone observed in these unmixed data are shown at the far left column. Clones are presented as described in Fig.4A. Transparency indicates a clone that was not detected with standard duplex filtering, but was detectable within single-strand consensus reads.

As shown below in Supplementary Figure S12, we observe concordance among technical replicates when the mutant clone is relatively less diluted from the original data, with reduced variant detection in duplex reads as the mutant read support is diminished. The missing variants can be rescued when examining single-strand consensus data. With increasing dilution, the variant eventually lacks sufficient read support to build both duplex or single-strand consensus reads, and is not detectable.

## Supplementary Note 4: Inferring population size and division rates from cell phylogenies Introduction

The aim of this exercise is to understand the apparent identifiability of the model parameters, seen in Bayesian inferences about the dynamics of HSC populations in mice (and other systems with similar population dynamics), when the phylogeny of a sample of descendent cells is the only available data. These Bayesian inferences were performed using ABC (approximate Bayesian computation) methods. Here the term approximate Bayesian computation refers to a class of Monte Carlo methods for generating samples from posterior distribution, which avoid computation of the likelihood function, by relying on simulation of the model. The more descriptive term *likelihood-free* Bayesian computation is also used. These methods include rejection sampling (pioneered by Pritchard *et al.*^99^), and various regression methods (introduced by Beaumont *et al.* ^41^).

The ABC results reported here were obtained by using the *rsimpop* package^34^ to perform simulations of models of cell population dynamics, and then using the *abc* package^100^ to compute (approximate) marginal posterior densities for modal parameters. The *rsimpop* package allows us to specify a wide range of stochastic growth models based on an underlying birth-death process.

In the case of neutral deterministic growth models, we have exact formulas for the likelihood function, where the model parameter is a sequence of *effective population sizes* (or a sequence of *drift intensities*). For these models, efficient Monte Carlo methods^101^ are available for sampling from the exact posterior distribution of the model parameters. In the case of stochastic growth models which can be approximated by a neutral deterministic growth model, we can obtain an approximate formulas for the likelihood function in which sequence of effective population sizes is replaced by a parameter vector which includes birth rates and death rates as model parameters. We will use approximate likelihood functions obtained in this way to address the issue of identifiability of parameters for various models.

### Likelihood functions for neutral models given phylogeny data

When we have genome sequences from a sample of single cells taken from an individual donor, we can construct a phylogeny for the sample, with the mutations assigned to branches. From this phylogeny, we can obtain an ultrametric tree, in which the relative lengths of the branches can be estimated (taking account of the number of mutations assigned to each branch). We also know the age *t*_*S*_ of the donor at the time point at which the sample was taken. (Here age is measured from the moment of conception.)

From the phylogeny on a sample of *n* cells, together with the estimated absolute branch lengths, we can label the internal nodes (coalescent event) with integers 2, 3, …, *n*, where *n* is the label on the most recent node (closest to the time of sample collection), and where 2 is the label on the earliest node on the phylogeny (the root node). We let *S* be the sequence of node heights (*S*(*n*), *S*(*n* − 1), …, *S*(2)), where *S*(*r*) is the height (time in days or years, measured backwards from sample collection) of internal node *r*. These node heights are determined by the branch lengths. The same information is contained in the sequence *T* of inter-coalescent interval durations (*T*(*n*), *T*(*n* − 1), …, *T*(2)). The inter-coalescent interval duration *T*(*r*) is the duration (in days or years) of the time interval during which exactly *r* lines of descent remain.

We begin by allowing the neutral model to take a very general form, which can be viewed as a generalisation of the neutral Moran model^102^. For now, we measure time *t* forward from conception (*t* = 0) when the population of cells contains a single founder cell (*N*_0_ = 1), which is the zygote. This time *t* coincides with age (measured from conception). The sequence of distinct time points (ages) at which the population size changes, together with the age *t*_*c*_ at the time of sample collection, is recorded as *t* = (*t*_1_, *t*_2_, …, *t*_*c*_), where 0 < *t*_1_ < *t*_2_ < ⋯ < *t*_*c*_. So we have a sequence of *c*-1 population events which occur before sample collection. We assume that at each population event, these changes in population size occur instantaneously, so that we can define a population size *N*_*k*_ which persists throughout the time interval [*t*_*k*_, *t*_*k*+1_⟩ from event *k* to the moment immediately preceding the next event.

At each of these events (*k* = 1, 2, …, *c*-1) at which the population size changes, we allow the number of *births b*_*k*_ (cell division) to be either 0 or 1, and the number of *deaths d*_*k*_ (cells which leave the stem cell population, either via cell deaths, or via cell differentiation events) to any integer value from 0 up to *N*_*k*−1_ (the size of the population when it enters event *k*).

Note that in a birth-death process it is more usual to assume that each event is either a birth event (where *b*_*k*_ = 1, and *d*_*k*_ = 0) or a death event (where *b*_*k*_ = 0, and *d*_*k*_ = 1). However, it turns out that while the analysis outlined below is greatly complicated if we allow *b*_*k*_ to exceed 1, when we relax the constraints on *d*_*k*_ we encounter very little additional difficulty. We have the following recursion for the population size

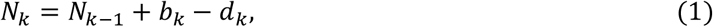

for *k* = 1, 2, …, *c*-1, subject to the constraints that either *b*_*k*_ = 0 or *b*_*k*_ = 1, and 0 ≤ *d*_*k*_ ≤ *N*_*k*−1_. There is one more sequence which it is useful for us to define here. This is the sequence of *drift intensities*, ξ = (ξ_1_, ξ_2_, …, ξ_*c*−1_), where

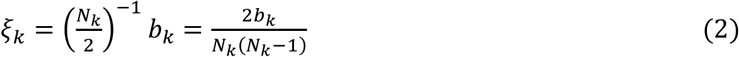

for *k* = 1, 2, …, *c* − 1. Recall that if there was no birth (cell division) at event *k*, then *b*_*k*_ = 0, and therefore ξ_k_ = 0. Notice that here we are using the conventional notation ^*n*^, for binomial coefficients. In particular we have

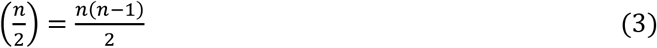

We can define the function

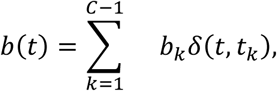

which represents the intensity of birth events. We can also define the function

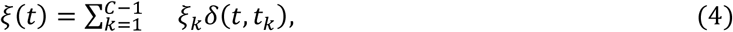

which represents the intensity of random drift.

We can express the drift intensity function as

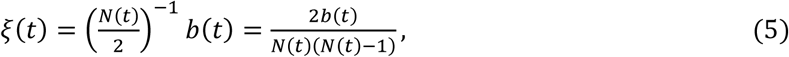

which is in agreement with the earlier definition (Equation 4). The trajectory of the intensity of random drift, as specified by the drift intensity function ξ(*t*) (Equations 4 and 5), takes us a step closer to our goal of deriving an expression for the likelihood function for the sample phylogeny data. However, in order to express the likelihood function in its most familiar and convenient form, we need to express the trajectory ξ(*t*) (and the related trajectories *N*(*t*), and so on) as functions of time *s* measured backwards from the time point at which the sample was collected (*s* = 0). The relationship between the forward time *t* (age from conception) and the backwards time *s*, is given by

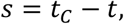

and hence *t* = *t*_*c*_ − *s*.

So we can represent the backwards time trajectory for population size as the function

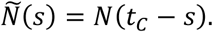

Similarly, we can represent the backwards time trajectories for other quantities of interest as follows

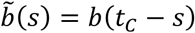

and

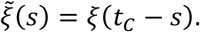

Now that we have this definition of the (reverse time) population size function *N*∽(*s*), we can express the (reverse time) drift intensity function as

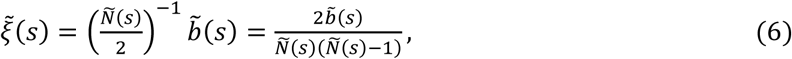

which is simply the reverse time version of Equation 5.

Recall that we defined the sequence *t* of distinct (forward) times (ages) at which the population size changes, together with the age *t*_*c*_ at the time of sample collection, *t* = (*t*_1_, *t*_2_, …, *t*_*c*_), where 0 < *t*_1_ < *t*_2_ < ⋯ < *t*_*c*_. The same sequence of time points, representing population events, which we have labelled with forward times (ages) *t*_*k*_, can also be labelled with reverse times *s*_*k*_ = *t*_*c*_ − *t*_*k*_, for *k* = 1, 2, …, *c*-1. We now define the sequence *s* of distinct reverse times at which the population size changes, together with the time *s*_0_ (= *t*_*c*_) at which conception occurred, *s* = (*s*_0_, *s*_1_, *s*_2_, …, *s*_*c*−1_), where *s*_*k*_ = *t*_*c*_ − *t*_*k*_, for each event *k*. Therefore, we have *s*_0_ > *s*_1_ > *s*_2_ > ⋯ > *s*_*c*−1_ > 0. The function ξ∽(*s*) (and the function ξ(*t*)) is completely determined by the sequence pair (*t*, ξ), and also by the (equivalent) sequence pair (*s*, ξ).

When the phylogeny, with (estimated) absolute branch lengths, is the only available data, the likelihood function of the model parameter given the data, is (up to a constant factor) equal to the joint probability density

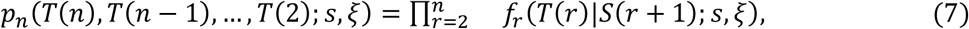

where

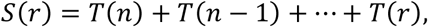

and where each factor

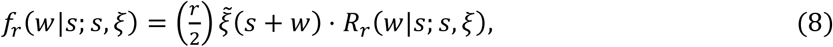

is the (marginal) probability density of the waiting time to the next coalescent event, starting from time point *s*, when *r* lines of descent remain (each of which can be traced back from the sample). The function ξ∽(*s*) is the drift intensity at time *s* (measured backwards from the time of sample collection). The function

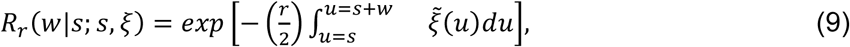

gives the probability that the waiting time to the next coalescent event (starting from time point *s*, when *r* lines of descent remain) is exceeds *w*. We could describe *R*_*r*_(*w*|*s*; *s*, ξ) as the *reliability* function (or survival function), and interpret *T*(*r*) as a kind of *failure time* (at which one line of descent fails to persist).

Strictly speaking, Equations 8 and 9 represent an approximation which is valid whenever the entire sample phylogeny lies within a time interval throughout which the intensity of random drift ξ∽(*s*) remains small (the effective population size remains large). See refs. ^103,104^ for derivation of the properties of the (reverse-time) genealogical process.

We want to draw attention to a feature of the likelihood function represented by Equations 7, 8, and 9. From the likelihood function (Equations 7 and 8) it is evident that, while segments (spanning certain time intervals) of the trajectory for the drift intensity (represented variously as a sequence pair *s*, ξ, or as a function of time), may constitute an identifiable parameter (when we have a phylogeny on a large enough sample), the trajectory for the population size, and the trajectory for the intensity of birth events, in the absence of additional constraints, are non-identifiable parameters. This is because the population sizes and the counts of birth events do not appear separately in the likelihood function, but only in the particular combination represented by the trajectory for the drift intensity.

In Section 3 below, parameter identifiability is defined more carefully, with some pointers to the literature. We also discuss in more detail the implications of non-identifiability for parameter estimation in our current model. In particular we will discuss how additional constraints on the population trajectory can restore identifiability of the population size, and the intensity of birth events.

### Parameter estimation and identifiability

We usually make some further assumptions about the possible trajectories which the population is allowed to follow through time. In the case of a deterministic growth model, we assume that the sequence pair (*s*, ξ) of event times and drift intensities belongs to a family of trajectories, in which the individual trajectory is completely determined by a parameter vector φ. (Typically this parameter vector is of low dimension.) We say that the family of trajectories is parametrised by φ. Here we have in mind models of deterministic exponential growth, where the model parameters include rates of cell division and rates of cell death.

In the case of a stochastic growth model, we assume that the sequence pair (*s*, ξ) is drawn from a distribution which belongs to some family of distributions. Within this family of distributions, the specific distribution is completely determined by a parameter vector φ. We say that the family of distributions is parametised by φ. Here we have in mind models based on a birth death process, where the model parameters again include rates of cell division and rates of cell death.

In order to emphasise the dependence on the parameter vector φ, it is convenient to use the notation

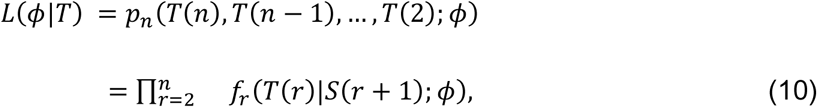

for the likelihood function specified by Equations 7 and 8.

We say that a parameter vector φ is *non-identifiable* whenever there is a mapping ν (to a vector of lower dimension) for which the likelihood function *L*(φ|*T*) depends on the parameter vector φ only through θ = ν(φ). In other words ν(φ_1_) = ν(φ_2_) implies that *L*(φ_1_|*T*) = *L*(φ_2_|*T*). If there is no such mapping ν, then we say that the parameter vector φ is *identifiable*. When the parameter vector φ is *identifiable*, we may also refer to the components of this vector as *identifiable* parameters. See ref. ^105^ (*non-identifiability* is introduced in Section 3.15, on page 70, and discussed further on pages 72 and 74).

If such a mapping ν (to a vector of lower dimension) exists (so that φ is non-identifiable), then this means (loosely speaking) that from the fixed data *T*, we can not learn anything about the unobserved parameter vector φ, beyond what we can learn about the (lower dimensional) parameter vector θ. We can state this more precisely. First, we can always (leaving aside technical issues and pathological cases) express the prior density π(φ) for the parameter vector φ, in the form

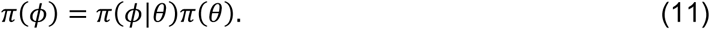

If φ is non-identifiable, and θ = ν(φ) is identifiable, then the posterior density π(φ|*T*) of the parameter vector φ is of the form

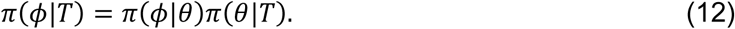

As a consequence, we also have

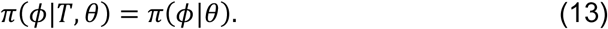

This means that if we knew the (lower dimensional) parameter vector θ, then the observed data *T* would tell us nothing more about the (higher dimensional) parameter vector φ.

First we consider a family of models where the population trajectory includes prolonged epochs during which birth events and death events occur equally often, so that the population size remains stable. Then we consider neutral models where the trajectory includes epochs of (deterministic) exponential population growth (Section 5). Finally, we consider birth-death processes, without an upper boundary (Section 6), and with an upper boundary (Section 7) on the population size, and how these stochastic growth models can be approximated by deterministic growth models.

### Epochs of stable effective population size

First we consider a family of models where the population trajectory includes prolonged epochs during which the population size remains stable. Suppose that across the time interval [*a*, *b*], the population size remains constant at *N*_*A*_. In order to maintain a constant population size, the birth rate β_*A*_ must be balanced by an equal death rate.

The observed inter-coalescent interval durations *T*(*r*), which fall within the time interval [*a*, *b*], contribute factors to the likelihood function which are of the form

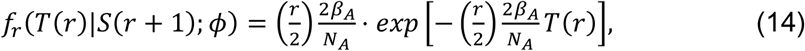

where φ = (*N*_*A*_, β_*A*_) is the parameter vector of the model.

From the expression on the right hand side of Equation 14, it appears that the only identifiable parameter is the ratio β_*A*_/*N*_*A*_.

### Epochs of exponential population growth

Now we turn to neutral models where the trajectory includes epochs of exponential population growth. Suppose that the (forward time) estimated trajectory ξ^(*t*) of the drift intensity appears to fit an exponential growth path across the time interval [*t*_*A*_, *t*_*c*_], where *t*_*c*_ is the time (age) at which the sample of *n* genome-sequenced cells was collected. The estimated trajectory ξ^(*t*) at time *t* can be interpreted as a kind of average drift intensity over some interval centred on the time point *t*. The (forward time) estimated trajectory is

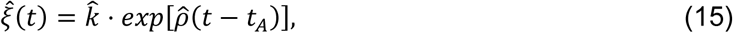

which is based on point estimates *p*^ (for the growth rate) and *k*^ (for the initial drift intensity). Notice that when *p*^ is positive, the drift intensity declines exponentially, with increasing age *t*.

If we measure time backwards from sample collection, then the (reverse time) estimated trajectory ξ∽^(*s*) of the drift intensity appears to fit an exponential growth path across the time interval [0, *s*_*A*_]. The (reverse time) estimated trajectory is

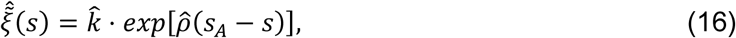

where *s*_*A*_ = *t*_*c*_ − *t*_*A*_ is the time measured backwards from sample collection to the time point at which the epoch of exponential growth began. Notice that when *p*^ is positive, the drift intensity increases exponentially, with increasing time *s*.

There is this one very simple model of population growth, in which births occur at a constant rate λ, and deaths occur at a constant rate *v*, which results in an exponential trajectory. This is an exceptionally parsimonious explanation for the observed exponential trajectory. If we can accept this parsimonious explanation, then we can set aside the general problem of making inferences about an arbitrary trajectory ξ(*t*) for the intensity of random drift (the reciprocal of the effective population size), and restrict our attention to the very specific problem of making inferences about the parameters of the deterministic exponential growth model, or the parameters of the birth death process.

Having observed an (approximately) exponential trajectory for the drift intensity (and its reciprocal, the effective population size), from age *t*_*A*_, up to the point of sample collection (at age *t*_*c*_), we have arrived at a parsimonious explanation which we now examine in more detail. The population size has been growing at a constant growth rate *ρ*, while the birth rate has remained constant at a value λ, and the death rate has remained constant at a value *v*, which yields the constant growth rate *ρ* = λ − *v*. Now we can express the trajectory for the population size *N*(*t*), forward in time across the epoch of exponential growth (from age *t*_*A*_ to age *t*_*c*_) as

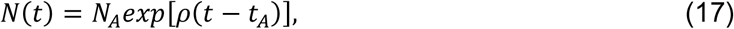

while the forward time trajectory for the drift intensity is

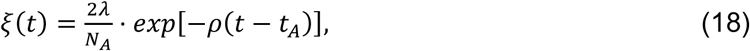

where *N*_*A*_ is the size of the ancestral population at age *t*_*A*_ (when the epoch of exponential growth begins).

We now return to time measured backwards from sample collection. The reverse time trajectory for the population size is

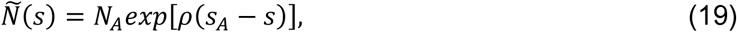

where *s*_*A*_ = *t*_*c*_ − *t*_*A*_ is the time measured backwards from sample collection to the time point at which the epoch of exponential growth began. The reverse time trajectory for the drift intensity is

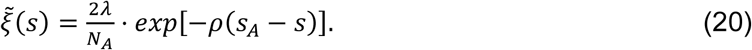

The (marginal) probability density *f*_*r*_(*w*|*s*; φ) of the waiting time to the next coalescent event (starting from time point *s*, when *r* lines of descent remain), is in this case

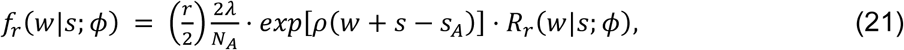

where φ = (λ, *v*, *N*_*A*_) is the parameter vector of this model, and where

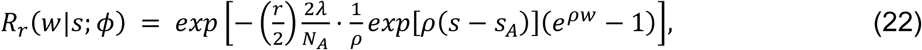

is the reliability function.

The observed inter-coalescent interval durations *T*(*r*), which fall within the time interval [0, *s*_*A*_] (the epoch of exponential growth), contribute factors to the likelihood function which are of the form

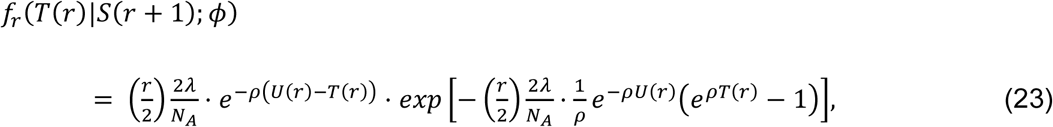

where *T*(*r*) = *s*_*A*_ − *S*(*r* + 1).

The parameter vector of this model is φ = (λ, *v*, *N*_*A*_), where λ is the birth rate, *v* is the death rate, and *N*_*A*_ is the size of the ancestral population at the start of the epoch of exponential growth. (This occurs at age *t*_*A*_, which precedes sample collection by time interval of duration *s*_0_ = *t*_*c*_ − *t*_*A*_.) From the formula for this factor of the likelihood function, it appears that the parameter vector φ is *non-identifiable*, while the parameter vector θ = (*N*_*A*_/λ, *ρ*) is *identifiable*. The components of the parameter vector θ are the ratio *N*_*A*_/λ, and the difference *ρ* = λ − *v* (the population growth rate).

In the special case where the epoch of exponential growth (at constant growth rate *ρ*) extends all the way back to the founding individual (zygote cell), we know *N*_*A*_ = 1, and we know that (reverse) time *s*_*A*_ = *s*_*c*_ (age *t*_*A*_ = 0) corresponds to the moment of conception. In this special case, the unobserved parameters λ and *v*, are identifiable. More generally, if the population size at the beginning of the epoch of exponential growth *N*_*A*_ is known with certainty, then the parameter vector θ = (λ, *v*) is *identifiable*.

In the case of a sample of single cell genome sequences obtained from blood-derived colonies, from a mouse (or any species with similar HSC dynamics), the parameter *N*_*A*_ is the size of the ancestral population of HSCs at age *t*_*A*_ (when the epoch of exponential growth begins); or if the time *t*_*A*_ is even earlier, then *N*_*A*_ is the size of the population of embryonic cells existing at this time which are ancestral to the HSCs. Unfortunately we do not have direct observations of the ancestral HSC population size *N*_*A*_ (at the age *t*_*A*_ when the epoch of exponential growth begins).

However, we can place some bounds on the value of *N*_*A*_. First of all there is an upper bound *M*_*A*_, on *N*_*A*_, which can be obtained from embryological observations. We know the approximate number of cells in the embryo at age *t*_*A*_. If some differentiation has already occurred, we may be able to exclude some cell types as HSC ancestors, and thus perhaps obtain an upper bound *M*_*A*_ which is somewhat lower than the average total number of cells in a mouse embryo at age *t*_*A*_. Secondly, we have a lower bound on *N*_*A*_, which we can obtain directly from the phylogeny. This the number of lines of descent *n*_*A*_ present on the tree at time *t*_*A*_.

### The linear birth-death process

A linear birth-death process is a simple stochastic growth model in which birth events and death events occur at constant rates (birth rate λ and death rate *v*) per individual (cell) per unit of time (day or year). Therefore the total rate of birth (respectively death) events in the population at each time point is proportional to the total number of individuals in the population at that time point (hence a *linear* birth-death process. The total size *N*(*t*) of the population at each time point is determined by the (stochastic) sequence of events (births and deaths) up to that time point. For the properties of the linear birth-death process, see ref. ^83^, and ref ^106^, pages 174–177.

Whenever the population size is not too small, and the growth rate is not too close to zero, the linear birth-death process behaves much like deterministic exponential growth. The trajectory for the population size *N*(*t*) is well approximated by Equation 17, with growth rate *ρ* = λ − *v*, provided that the birth rate λ exceeds the death rate *v*, so that *ρ* is positive.

In the case of an epoch of stochastic growth (under a linear birth-death process) it is important to bear in mind that the formula for the factors of the likelihood function in Equation 21, is an approximation, which can break-down. A conclusive argument about the identifiability of the model parameters should be based on an exact formula for the likelihood function for the linear birth-death process, when the phylogeny is the only available data.

### The birth-death process with an upper boundary on population size

If a mouse lives long enough, we would expect that the propensity of the mouse HSC population to grow exponentially will eventually be checked by the physical constraints on the space available to accommodate the HSC cells within the bone marrow.

In the case of a model where the population undergoes deterministic exponential growth until an upper boundary *N*_*B*_ on population size is reached, the phylogeny may contain additional information about the time *T*_*B*_ at which the population first hits the upper boundary *N*_*B*_. Such information can be present only if the sample of cells has been collected from the population at a time point after the time *T*_*B*_. In this case, the hitting time parameter *T*_*B*_ occurs in the likelihood function.

In the case of a model where the population undergoes deterministic exponential growth until an upper boundary *N*_*B*_ population size is reached. The hitting time *T*_*B*_ is determined by model parameters (*N*_*A*_/*N*_*B*_ and *p* = λ − *v*). Using Equation 17, we can obtain

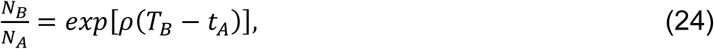

and therefore

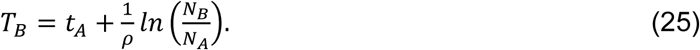

When the population reaches the upper boundary on population size, the marginal birth rate and the marginal death rate must be equal (δ_*B*_ = β_*B*_). The parameter vector of the model is now φ = (λ, *v*, *N*_*A*_, *N*_*B*_, β_*B*_).

As usual we inspect the formula for the likelihood function in order to discover which parameters may be identifiable, and which are clearly non-identifiable. The factors of the likelihood function representing the epoch of exponential growth are of the form given in Equation 21, in which the parameter combinations λ/*N*_*A*_ and *ρ* appear. The factors of the likelihood function representing the epoch of stable population size are of the form given in Equation 14, in which the parameter combination β_*B*_/*N*_*B*_ appears. We have also seen from Equation 24 that the ratio *N*_*B*_ /*N*_*A*_ is determined by the parameter *p* and the the hitting time *T*_*B*_. The hitting time *T*_*B*_ is a change point, which also appears in the likelihood function. Therefore, from the formulas for the factors of the likelihood function, it appears that the parameter vector θ = (*ρ*, λ/*N*_*A*_, β_*B*_/*N*_*B*_, *N*_*B*_ /*N*_*A*_) is identifiable. Notice also that by combining the last three components of θ, we obtain

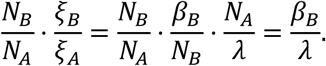

So the ratio β_*B*_/λ is also identifiable.

In the special case where *N*_*A*_ is known for certain, the parameter vector θ = (λ, *v*, *N*_*B*_, β_*B*_) is identifiable. As already discussed in Section 5, when the epoch of exponential growth (at constant growth rate *ρ*) extends all the way back to the founding individual (zygote cell), we know *N*_*A*_ = 1. So, in this case, the parameters λ, *v*, *N*_*B*_, and β_*B*_, are all identifiable, and amenable to estimation from the phylogeny of a sample.

## REFERENCES

1. Sender, R. & Milo, R. The distribution of cellular turnover in the human body. Nat Med 27, 45–48 (2021).

2. Patel, S. H. et al. Lifelong multilineage contribution by embryonic-born blood progenitors. Nature 606, 747–753 (2022).

3. Sun, J. et al. Clonal dynamics of native haematopoiesis. Nature 514, 322–327 (2014).

4. Kucinski, I. et al. A time- and single-cell-resolved model of murine bone marrow hematopoiesis. Cell Stem Cell 31, 244–259.e10 (2024).

5. Takizawa, H., Regoes, R. R., Boddupalli, C. S., Bonhoeffer, S. & Manz, M. G. Dynamic variation in cycling of hematopoietic stem cells in steady state and inflammation. J Exp Med 208, 273– 284 (2011).

6. Munz, C. M. et al. Regeneration after blood loss and acute inflammation proceeds without contribution of primitive HSCs. Blood 141, 2483–2492 (2023).

7. Fanti, A.-K. et al. Flt3- and Tie2-Cre tracing identifies regeneration in sepsis from multipotent progenitors but not hematopoietic stem cells. Cell Stem Cell 30, 207–218.e7 (2023).

8. Trumpp, A., Essers, M. & Wilson, A. Awakening dormant haematopoietic stem cells. Nature Reviews Immunology 10, 201–209 (2010).

9. Mitchell, E. et al. Clonal dynamics of haematopoiesis across the human lifespan. Nature 606, 343–350 (2022).

10. Lee-Six, H. et al. Population dynamics of normal human blood inferred from somatic mutations. Nature 561, 473–478 (2018).

11. Osorio, F. G. et al. Somatic Mutations Reveal Lineage Relationships and Age-Related Mutagenesis in Human Hematopoiesis. Cell Reports 25, 2308–2316.e4 (2018).

12. Abascal, F. et al. Somatic mutation landscapes at single-molecule resolution. Nature 593, 405–410 (2021).

13. Spencer Chapman, M., et al. Lineage tracing of human development through somatic mutations. Nature 595, 85–90 (2021).

14. Jaiswal, S. Clonal hematopoiesis and nonhematologic disorders. Blood 136, 1606–1614 (2020).

15. Xie, M. et al. Age-related mutations associated with clonal hematopoietic expansion and malignancies. Nature Medicine 20, 1472–1478 (2014).

16. Jaiswal, S. & Ebert, B. L. Clonal hematopoiesis in human aging and disease. Science 366, (2019).

17. Kapadia, C. D. & Goodell, M. A. Tissue mosaicism following stem cell aging: blood as an exemplar. Nat Aging 4, 295–308 (2024).

18. Moore, L. et al. The mutational landscape of normal human endometrial epithelium. Nature 580, 640–646 (2020).

19. Martincorena, I. et al. High burden and pervasive positive selection of somatic mutations in normal human skin. Science 348, 880–886 (2015).

20. Lawson, A. R. J. et al. Extensive heterogeneity in somatic mutation and selection in the human bladder. Science 370, 75–82 (2020).

21. Martincorena, I. et al. Somatic mutant clones colonize the human esophagus with age. Science 362, 911–917 (2018).

22. Ng, S. W. K. et al. Convergent somatic mutations in metabolism genes in chronic liver disease. Nature 598, 473–478 (2021).

23. Cagan, A. et al. Somatic mutation rates scale with lifespan across mammals. Nature 604, 517–524 (2022).

24. Yuan, R. et al. Genetic coregulation of age of female sexual maturation and lifespan through circulating IGF1 among inbred mouse strains. Proc Natl Acad Sci U S A 109, 8224– 8229 (2012).

25. Chin, D. W. L. et al. Aged healthy mice acquire clonal hematopoiesis mutations. Blood 139, 629–634 (2022).

26. Osawa, M., Hanada, K., Hamada, H. & Nakauchi, H. Long-Term Lymphohematopoietic Reconstitution by a Single CD34-Low/Negative Hematopoietic Stem Cell. Science 273, 242– 245 (1996).

27. Adolfsson, J. et al. Upregulation of Flt3 Expression within the Bone Marrow Lin−Sca1+c-kit+ Stem Cell Compartment Is Accompanied by Loss of Self-Renewal Capacity. Immunity 15, 659–669 (2001).

28. Kiel, M. J. et al. SLAM family receptors distinguish hematopoietic stem and progenitor cells and reveal endothelial niches for stem cells. Cell 121, 1109–1121 (2005).

29. Challen, G. A., Boles, N. C., Chambers, S. M. & Goodell, M. A. Distinct Hematopoietic Stem Cell Subtypes Are Differentially Regulated by TGFβ1. Cell Stem Cell 6, 265–278 (2010).

30. Cabezas-Wallscheid, N. et al. Identification of regulatory networks in HSCs and their immediate progeny via integrated proteome, transcriptome, and DNA methylome analysis. Cell Stem Cell 15, 507–522 (2014).

31. Pietras, E. M. et al. Functionally Distinct Subsets of Lineage-Biased Multipotent Progenitors Control Blood Production in Normal and Regenerative Conditions. Cell Stem Cell 17, 35–46 (2015).

32. Sun, J. et al. Clonal dynamics of native haematopoiesis. Nature 514, 322–327 (2014).

33. Busch, K. et al. Fundamental properties of unperturbed haematopoiesis from stem cells in vivo. Nature 518, 542–546 (2015).

34. Williams, N. et al. Life histories of myeloproliferative neoplasms inferred from phylogenies. Nature 602, 162–168 (2022).

35. Machado, H. E. et al. Diverse mutational landscapes in human lymphocytes. Nature 608, 724–732 (2022).

36. Coorens, T. H. H. et al. Inherent mosaicism and extensive mutation of human placentas. Nature 592, 80–85 (2021).

37. Bryder, D., Rossi, D. J. & Weissman, I. L. Hematopoietic stem cells: the paradigmatic tissue-specific stem cell. Am J Pathol 169, 338–346 (2006).

38. Qin, P. et al. Integrated decoding hematopoiesis and leukemogenesis using single-cell sequencing and its medical implication. Cell Discov 7, 2 (2021).

39. de Haan, G. & Van Zant, G. Dynamic Changes in Mouse Hematopoietic Stem Cell Numbers During Aging. Blood 93, 3294–3301 (1999).

40. Morrison, S. J., Wandycz, A. M., Akashi, K., Globerson, A. & Weissman, I. L. The aging of hematopoietic stem cells. Nat Med 2, 1011–1016 (1996).

41. Beaumont, M. A., Zhang, W. & Balding, D. J. Approximate Bayesian computation in population genetics. Genetics 162, 2025–2035 (2002).

42. Koonin, E. V. Splendor and misery of adaptation, or the importance of neutral null for understanding evolution. BMC Biology 14, 114 (2016).

43. Challen, G. A. & Goodell, M. A. Clonal hematopoiesis: mechanisms driving dominance of stem cell clones. Blood 136, 1590–1598 (2020).

44. Yoshizato, T. et al. Somatic Mutations and Clonal Hematopoiesis in Aplastic Anemia. New England Journal of Medicine 373, 35–47 (2015).

45. King, K. Y., Huang, Y., Nakada, D. & Goodell, M. A. Environmental influences on clonal hematopoiesis. Experimental Hematology 83, 66–73 (2020).

46. Florez, M. A. et al. Clonal hematopoiesis: Mutation-specific adaptation to environmental change. Cell Stem Cell 29, 882–904 (2022).

47. Coombs, C. C. et al. Therapy-Related Clonal Hematopoiesis in Patients with Non-hematologic Cancers Is Common and Associated with Adverse Clinical Outcomes. Cell Stem Cell 21, 374–382.e4 (2017).

48. Bolton, K. L. et al. Cancer therapy shapes the fitness landscape of clonal hematopoiesis. Nat Genet 52, 1219–1226 (2020).

49. Wong, T. N. et al. Role of TP53 mutations in the origin and evolution of therapy-related acute myeloid leukaemia. Nature 518, 552–555 (2015).

50. Beura, L. K. et al. Normalizing the environment recapitulates adult human immune traits in laboratory mice. Nature 532, 512–516 (2016).

51. Camell, C. D. et al. Senolytics reduce coronavirus-related mortality in old mice. Science 373, (2021).

52. Matatall, K. A. et al. Chronic Infection Depletes Hematopoietic Stem Cells through Stress-Induced Terminal Differentiation. Cell Rep 17, 2584–2595 (2016).

53. Watson, C. J. et al. The evolutionary dynamics and fitness landscape of clonal hematopoiesis. Science 367, 1449–1454 (2020).

54. Kimura, M. & Ohta, T. The Average Number of Generations until Fixation of a Mutant Gene in a Finite Population. Genetics 61, 763–771 (1969).

55. Challen, G. A., Pietras, E. M., Wallscheid, N. C. & Signer, R. A. J. Simplified murine multipotent progenitor isolation scheme: Establishing a consensus approach for multipotent progenitor identification. Experimental Hematology 104, 55–63 (2021).

56. Sheikh, B. N. et al. MOZ (KAT6A) is essential for the maintenance of classically defined adult hematopoietic stem cells. Blood 128, 2307–2318 (2016).

57. Kobayashi, M. et al. HSC-independent definitive hematopoiesis persists into adult life. Cell Reports 42, 112239 (2023).

58. Abkowitz, J. L., Catlin, S. N., McCallie, M. T. & Guttorp, P. Evidence that the number of hematopoietic stem cells per animal is conserved in mammals. Blood 100, 2665–2667 (2002).

59. Dykstra, B. et al. Long-term propagation of distinct hematopoietic differentiation programs in vivo. Cell Stem Cell 1, 218–229 (2007).

60. Gros, P. & Casanova, J.-L. Reconciling Mouse and Human Immunology at the Altar of Genetics. Annu Rev Immunol 41, 39–71 (2023).

61. Lindsay, S. J., Rahbari, R., Kaplanis, J., Keane, T. & Hurles, M. E. Similarities and differences in patterns of germline mutation between mice and humans. Nat Commun 10, 4053 (2019).

62. Bergeron, L. A. et al. Evolution of the germline mutation rate across vertebrates. Nature 615, 285–291 (2023).

63. Gould, S. J., Lewontin, R. C., Maynard Smith, J. & Holliday, R. The spandrels of San Marco and the Panglossian paradigm: a critique of the adaptationist programme. Proceedings of the Royal Society of London. Series B. Biological Sciences 205, 581–598 (1997).

64. Brayton, C. F., Treuting, P. M. & Ward, J. M. Pathobiology of aging mice and GEM: background strains and experimental design. Vet Pathol 49, 85–105 (2012).

65. Shepherd, B. E. et al. Hematopoietic stem-cell behavior in nonhuman primates. Blood 110, 1806–1813 (2007).

66. Koelle, S. J. et al. Quantitative stability of hematopoietic stem and progenitor cell clonal output in rhesus macaques receiving transplants. Blood 129, 1448–1457 (2017).

67. Shin, T.-H. et al. A macaque clonal hematopoiesis model demonstrates expansion of TET2-disrupted clones and utility for testing interventions. Blood 140, 1774–1789 (2022).

68. Yu, K.-R. et al. The impact of aging on primate hematopoiesis as interrogated by clonal tracking. Blood 131, 1195–1205 (2018).

69. Hsu, J. I. et al. PPM1D Mutations Drive Clonal Hematopoiesis in Response to Cytotoxic Chemotherapy. Cell Stem Cell 23, 700–713.e6 (2018).

70. Heyde, A. et al. Increased stem cell proliferation in atherosclerosis accelerates clonal hematopoiesis. Cell 184, 1348–1361.e22 (2021).

71. Meisel, M. et al. Microbial signals drive pre–leukaemic myeloproliferation in a Tet2– deficient host. Nature 557, 580–584 (2018).

72. Hormaechea-Agulla, D. et al. Chronic infection drives Dnmt3a-loss-of-function clonal hematopoiesis via IFNγ signaling. Cell Stem Cell 28, 1428–1442.e6 (2021).

73. Cheshier, S. H., Morrison, S. J., Liao, X. & Weissman, I. L. In vivo proliferation and cell cycle kinetics of long-term self-renewing hematopoietic stem cells. Proceedings of the National Academy of Sciences 96, 3120–3125 (1999).

74. Wilson, A. et al. Hematopoietic Stem Cells Reversibly Switch from Dormancy to Self-Renewal during Homeostasis and Repair. Cell 135, 1118–1129 (2008).

75. Bernitz, J. M., Kim, H. S., MacArthur, B., Sieburg, H. & Moore, K. Hematopoietic Stem Cells Count and Remember Self-Renewal Divisions. Cell 167, 1296–1309.e10 (2016).

76. Chen, J., Astle, C. M. & Harrison, D. E. Genetic regulation of primitive hematopoietic stem cell senescence. Exp Hematol 28, 442–450 (2000).

77. Ahmad, A. et al. ERCC1-XPF endonuclease facilitates DNA double-strand break repair. Mol Cell Biol 28, 5082–5092 (2008).

78. Nadon, N. L., Strong, R., Miller, R. A. & Harrison, D. E. NIA Interventions Testing Program: Investigating Putative Aging Intervention Agents in a Genetically Heterogeneous Mouse Model. EBioMedicine 21, 3–4 (2017).

79. Bowie, M. B. et al. Hematopoietic stem cells proliferate until after birth and show a reversible phase-specific engraftment defect. J Clin Invest 116, 2808–2816 (2006).

80. Ellis, P. et al. Reliable detection of somatic mutations in solid tissues by laser-capture microdissection and low-input DNA sequencing. Nat Protoc 16, 841–871 (2021).

81. Jones, D. et al. cgpCaVEManWrapper: Simple Execution of CaVEMan in Order to Detect Somatic Single Nucleotide Variants in NGS Data. Curr Protoc Bioinformatics 56, 15.10.1–15.10.18 (2016).

82. Ye, K., Schulz, M. H., Long, Q., Apweiler, R. & Ning, Z. Pindel: a pattern growth approach to detect break points of large deletions and medium sized insertions from paired-end short reads. Bioinformatics 25, 2865–2871 (2009).

83. Kendall, D. G. Stochastic Processes and Population Growth. Journal of the Royal Statistical Society. Series B (Methodological) 11, 230–282 (1949).

84. Challen, G. A., Boles, N., Lin, K. K.-Y. & Goodell, M. A. Mouse hematopoietic stem cell identification and analysis. Cytometry. Part A: the journal of the International Society for Analytical Cytology 75, 14–24 (2009).

85. Lee-Six, H. et al. The landscape of somatic mutation in normal colorectal epithelial cells. Nature 574, 532–537 (2019).

86. Durand, J.-B., Goncalves, P. & Guedon, Y. Computational methods for hidden Markov tree models-an application to wavelet trees. IEEE Transactions on Signal Processing 52, 2551–2560 (2004).

87. Kennedy, S. R. et al. Detecting ultralow-frequency mutations by Duplex Sequencing. Nature protocols 9, 2586–606 (2014).

88. Lai, Z. et al. VarDict: a novel and versatile variant caller for next-generation sequencing in cancer research. Nucleic Acids Res 44, e108 (2016).

89. McLaren, W. et al. The Ensembl Variant Effect Predictor. Genome Biology 17, 122 (2016).

90. Costello, M. et al. Discovery and characterization of artifactual mutations in deep coverage targeted capture sequencing data due to oxidative DNA damage during sample preparation. Nucleic Acids Res 41, e67 (2013).

91. Alexandrov, L. B. et al. The repertoire of mutational signatures in human cancer. Nature 578, 94–101 (2020).

92. Feng, C. G., Weksberg, D. C., Taylor, G. A., Sher, A. & Goodell, M. A. The p47 GTPase Lrg-47 (Irgm1) links host defense and hematopoietic stem cell proliferation. Cell Stem Cell 2, 83– 89 (2008).

93. Lerner, C. & Harrison, D. E. 5-Fluorouracil spares hemopoietic stem cells responsible for long-term repopulation. Exp Hematol 18, 114–118 (1990).

94. Dong, S. et al. Chaperone-mediated autophagy sustains haematopoietic stem-cell function. Nature 591, 117–123 (2021).

95. Martincorena, I. et al. Universal Patterns of Selection in Cancer and Somatic Tissues. Cell 171, 1029–1041.e21 (2017).

96. Campbell, P. et al. Clonal dynamics after allogeneic haematopoietic cell transplantation using genome-wide somatic mutations. Preprint at 10.21203/rs.3.rs-2868644/v1 (2023).

97. Poon, G. Y. P., Watson, C. J., Fisher, D. S. & Blundell, J. R. Synonymous mutations reveal genome-wide levels of positive selection in healthy tissues. Nat Genet 53, 1597–1605 (2021).

98. Flurkey, K., M. Currer, J. & Harrison, D. E. Mouse Models in Aging Research. in The Mouse in Biomedical Research (Second Edition) (eds. Fox, J. G. et al.) 637–672 (Academic Press, Burlington, 2007). doi:10.1016/B978-012369454-6/50074-1.

99. Pritchard, J. K., Seielstad, M. T., Perez-Lezaun, A. & Feldman, M. W. Population growth of human Y chromosomes: a study of Y chromosome microsatellites. Molecular Biology and Evolution 16, 1791–1798 (1999).

100. Csilléry, K., François, O. & Blum, M. G. abc: an R package for approximate Bayesian computation (ABC). Methods in ecology and evolution 3, 475–479 (2012).

101. Lan, S., Palacios, J. A., Karcher, M., Minin, V. N. & Shahbaba, B. An efficient Bayesian inference framework for coalescent-based nonparametric phylodynamics. Bioinformatics 31, 3282–3289 (2015).

102. Moran, P. A. P. Random processes in genetics. Proceedings of the Cambridge Philosophical Society 54, 60–71 (1958).

103. Kingman, J. F. C. On the Genealogy of Large Populations. J. Appl. Probab. **19A**, 27–43 (1982).

104. Griffiths, R. C. & Tavaré, S. Sampling theory for neutral alleles in a varying environment. *Philosophical Transactions of the Royal Society, London*, Series B 344, 403–410 (1994).

105. O’Hagan, A. & Forster, J. Bayesian Inference. vol. 2B (Arnold, London, UK, 2004).

106. Moran, P. A. P. An Introduction to Probability Theory. (Oxford University Press, Oxford, UK, 1968).

